# Wide-scale Geographical Analysis of Genetic Ancestry in the South African Coloured Population

**DOI:** 10.1101/2024.06.13.598620

**Authors:** Imke Lankheet, Rickard Hammarén, Lucía Ximena Alva Caballero, Maximilian Larena, Helena Malmström, Cecile Jolly, Himla Soodyall, Michael de Jongh, Carina Schlebusch

**Author notes:** Corresponding authors: Carina Schlebusch, Imke Lankheet. These authors contributed equally (Imke Lankheet, Rickard Hammarén, Lucía Ximena Alva Caballero).

## Abstract

The South African Coloured (SAC) population, a prominent admixed population in South Africa, reflects centuries of migration, admixture, and historical segregation. Descendants of local Khoe-San and Bantu-speaking populations, European settlers, and enslaved individuals from Africa and Asia, SAC individuals embody diverse ancestries. This study investigates the genetic makeup of SAC individuals, utilizing autosomal genotypes, mitochondrial DNA and Y-chromosome data. We analyze new genotype data for 125 SAC individuals from seven locations. Our analysis, based on a dataset comprising 356 SAC individuals from 22 geographic locations, revealed significant regional variations in ancestry. Khoe-San ancestry predominates in 14 locations, highlighting its lasting influence. Inland regions exhibit higher proportions of Khoe-San ancestry, eastern regions show more Bantu-speaker/West African ancestry, and western/coastal areas, particularly around Cape Town, display increased Asian ancestry. These patterns reflect historical migrations and settlement patterns. Additionally, sex-biased admixture ratios show male-biased admixture from East Africans and Europeans, and female-biased admixture from Khoe-San populations, which is supported by mitochondrial and Y-chromosome data. This research underscores the importance of studying the SAC population to understand South Africa’s historical migrations, providing insights into the complex genetic heritage of South Africans.

## Introduction

The South African Coloured (SAC) population is among the most admixed populations in the world, and SAC individuals trace their genetic roots to local Khoe-San and Bantu-speaking groups, European colonists, and enslaved people from other regions in Africa as well as from Asia. Genetically, Khoe-San populations represent one of the two branches of the earliest population divergence of the human population tree and therefore show high genetic diversity [1, 2, 3]. They also host early diverging mitochondrial and Y-chromosome lineages [4, 5, 6, 7]. Until approximately 2000 years ago, the San ancestors were the only inhabitants of Southern Africa and they practiced hunter-gathering [8]. Around 2000 years ago, East-African pastoralists arrived in Southern Africa, and admixed with the local San populations [1, 2, 9, 10, 11], which gave rise to the Khoekhoe herding groups. Today, Khoe-San is the term used to refer to both populations collectively; the hunter-gatherer San and the herder Khoekhoe [12, 13]. The arrival of East-African pastoralists was followed by the arrival of Bantu-speaking groups practicing agriculture and carrying West African ancestry around 1800 years ago as part of the Bantu expansion [14, 15, 16, 17, 18]. The colonial times introduced both European and Asian ancestries into Southern Africa [19]. In 1652, the Dutch East India company founded a small refueling station that gradually grew over the decades into what became known as the Cape Colony and later on as Cape Town. The Dutch settlers interacted heavily with the local Khoekhoe communities from the very foundation of the colony. They traded for cattle and, as time went by, some Khoekhoe would work on settler farmsteads [20]. There were disproportionately few women among the settlers in the colony which led to formal and informal unions between European men and Khoekhoe women [20].

Over time, non-Europeans in the colony became less accepted, leading to the formation of a distinct community. Sometime after 1700, the term “Cape coloureds” emerged to refer to people of mixed ancestry [21]. The Cape coloureds were descendants of Khoe-San, Bantu-speaking populations, European settlers and enslaved people from the West and East Coast of Africa, the Indian subcontinent, Madagascar, and Indonesia, brought to South Africa during the slave trade period (1658-1806) [20]. The apartheid regime, the institutionalised racial segregation in place from 1948 to the early 1990s, enhanced the unity of the South African Coloured group identity [22, 20].

Currently, the SAC population is the largest admixed population in the country [22]. They constitute more than half of the population of the Western Cape Province today, with large presences in the Northern and Eastern Cape provinces as well, see Figure 2F. The majority of the SAC speak Afrikaans as their first language, 75.8% according to the 2011 South Africa (SA) census, and most SAC individuals identify as Christian. Religion serves as an essential characteristic that differentiates SAC from the Cape Malay population, who practice Islam [21] and are also the result of admixture events between Africans and Asians [23]. Despite the term “Coloured” originating as a construct during the apartheid regime, its usage persists in contemporary South Africa, albeit with varied acceptance.

A number of studies has investigated the genetics of the SAC individuals [24, 25, 26, 11, 27]. They confirm the inferences drawn from historical records: the six main demographic groups that contributed to the genetic pool of the SAC were the Khoe-San, Bantu-speakers/West Africans, East Africans, South Asians/Indians, Southeast Asians and Europeans. A mitochondrial DNA study revealed that the Khoe-San had a large maternal contribution to the SAC (60.0%), while the West Eurasian/European maternal contribution was very limited (4.6%) [25]. However, most of these studies focused on single locations and the majority of locations were close to Cape Town.

In this study, we analyzed new genome-wide data for 125 individuals self-identifying as SAC. The studied individuals are from a wide range of locations, spanning the broad geographic region inhabited by the SAC community, thereby providing a more comprehensive representation of this population. Together with previously published genetic data from SAC individuals, as well as comparative groups, we shed light on the ancestral genetic components present in various SAC populations and identify geographical differences among these components. We also investigate the difference in paternal and maternal contributions for various admixture events through mitogenomes and Y chromosomes, as well as through X-to-autosomal comparisons.

## Materials and Methods

### Sampling and genome-wide SNP typing

Saliva samples were obtained from 152 SAC individuals from seven different sites in South Africa; two in the Eastern Cape Province (Graaff-Reinet (N=45) and Nieu-Bethesda(N=20)), and five in the Western Cape Province (Genadendal (N=29), Greyton (N=16), Kranshoek (N=11), Oudtshoorn (N=17), and Prince Albert (N=14)). Participants donated saliva samples with written informed consent. Sample collection of SAC, Khoe-San and Khoe-San descendent groups were approved by the University of the Witwatersrand Human Research Ethics board, clearance numbers M980553, with renewals M050902, M090576, M1604104. This specific project was approved by the University of the Witwatersrand Human Research Ethics board, clearance number M180655 and the National Ethics review board of Sweden, clearance number Dnr 2021-01448.

The samples were obtained using an Oragene DNA OG-500 kit. DNA was extracted using the prepIT L2P extraction protocol. The extraction of the biological samples and genotyping followed the procedure described in [11]. The data were generated in four genotyping runs on the Illumina Infinium™ H3Africa Consortium Array by the SNP&SEQ Technology Platform in Uppsala, Sweden. Datasets were analyzed using GenomeStudio 2.0.3 and aligned to the Human Genome build version 37 (hg19). A total of 2,267,346 SNP markers were collected in genotyping run 1, 2, and 3, and 2,271,503 SNP markers were collected in genotyping run 4.

### Quality filtering and autosomal dataset merging

The genotype data from 152 SAC individuals was merged with the same dataset as used in [11] [11, 28, 29, 30, 31, 32, 33, 34] as well as with data from additional sources [1, 26, 35, 19]. For further information about the populations included in this study, see Supplementary Table 1. The Petersen dataset was converted to hg37 positions with the LiftOver tool from the Univeristy of California Santa Cruz (UCSC) (https://genome.ucsc.edu/cgi-bin/hgLiftOver). PLINK v1.90b4.9 [36] was used to carry out data processing and quality filtering. Before merging the datasets, duplicate SNPs were removed, only overlapping SNPs between datasets were kept, and C/G and A/T SNPs were eliminated to prevent strand flipping errors. Moreover, 5 individuals with genotyping missingness higher than 15% were excluded (plink –mind 0.15) and SNPs with less than 10% genotyping rate (plink –geno 0.1) were also excluded. Hardy–Weinberg Equilibrium (HWE) was set to 0.00001 (plink –hwe 0.00001) to avoid potential genotyping errors. Once the merging was done, analyses were performed to filter out one individual within each pair of relatives (second-degree or closer) using KING [37]. In total, 22 individuals were removed due to relatedness. Also, SNPs with less than 10% genotyping rate (plink –geno 0.1) were excluded again. To prevent ADMIXTURE and PCA analysis from being negatively affected by linkage disequilibrium (LD) bias, SNPs in LD were removed (plink–indep-pairwise 200 25 0.4). Each of the comparative populations was randomly sub-sampled to 30 individuals per population to avoid a sample-size bias in further analysis. The final dataset comprised 162 382 SNPs and 1203 individuals, of which 356 were SAC individuals and 125 were newly typed SAC individuals. Geographic information of the SAC individuals from previously published data was obtained from their respective publications. Sampling locations are displayed in Supplementary Figure 1.

### Population structure inferences

Unsupervised population structure inference analysis for K = 2 to K = 12 was performed with ADMIXTURE [38] version 1.3.0 using a random seed each time, and repeated 50 times. PONG version 1.5 [39] was used to visualize the results and find the major mode and pairwise similarity. Principal component analysis (PCA) was performed using the program smartpca, from the Eigensoft package (version 7.2.1) [40, 41]. To capture more of the global variation, Uniform Manifold Approximation and Projection for Dimension Reduction (UMAP) was performed on the genotypes directly using the umap-learn python library version 0.5.3.

### Phasing, local ancestry estimation and admixture dating

Phasing was carried out out using SHAPEIT version 2.r837 [42] using the 1000 genomes phase 3 reference genomes [31] and options --states 500 --main 20 --burn 10 --prune 10. Any misaligned sites between the reference dataset and the panel were excluded. Local ancestry estimation was performed using MOSAIC version 1.5.0 compiled and ran under R version 4.3.2 [43], setting source populations to five (-a 5). Admixture dates were gathered from the reported dates from the co-ancestry curves. The origin of each reconstructed ancestry was determined through F_st_ to the reference populations using MOSAIC.

### Formal tests of admixture

The dataset was merged with a chimpanzee genome and f4-statistics were computed using popstats [44, 45] in the format f4(Chimp,SAC,Pop1,Pop2). It was used to test whether SAC individuals were more admixed with Pop1 or Pop2. If the f4-value is significantly negative, it implies gene flow between either SAC and Pop1 (or Chimp and Pop2). If it is significantly positive, it implies gene flow between SAC and Pop2 (or Chimp and Pop1).

### Sex-biased admixture

To test if the admixture was sex-biased, X-chromosome/Autosomal ratios were computed for the SAC individuals. The genotyping data from the individuals from these seven newly sampled sites were merged with a comparative dataset consisting of 20 Central Europeans (CEU), 20 Sri Lankan Tamil (STU), 20 Nigerian Yoruba (YRI), 20 Ethiopian Amhara and 17 Namibian Ju/’hoansi. The data were filtered as described in the section *Quality filtering and autosomal dataset merging*. To avoid differences in chromosome size affecting the admixture proportions, chromosome 1 to 6 were cut to the length of the X-chromosome (180 centiMorgan), and chromosome 7, 10 and 12 were selected as they roughly have the same length (in centiMorgan) as the X-chromosome. For the autosomes, the number of SNPs was downsampled to the number of SNPs found on the X-chromosome (7452 SNPs). Supervised ADMIXTURE (K = 5) was run separately for each of the autosomes and the X-chromosome with 50 iterations each [38]. The results were visualized with Pong [39]. The ADMIXTURE results provided the ancestry proportions on the X-chromosome per individual and per ancestry. Average autosomal proportions were calculated from the ADMIXTURE runs of each of the autosomes, for each individual and each ancestry. Female X-chromosomal proportions were weighed twice, as females have two X-chromosomes and males only one [46]. One individual was removed because we lacked information to determine whether it was male or female, both from the informed consent form as well as plink sex analysis (plink –check-sex). Corrected X and autosomal proportions were bootstrapped (10 000 times) and average X-to-autosomal difference ratios were calculated as in [47] for each of the five ancestries as follows:

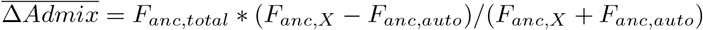

where *F*_*anc,total*_ is the genome-wide admixture proportion for a given ancestry, *F*_*anc,X*_ is the X chromosome admixture proportion for a given ancestry and *F*_*anc,auto*_ is the autosomal admixture proportion for a given ancestry. Negative X-to-autosomal difference ratios are indicative of male-biased admixture for that ancestry, positive X-to-autosomal difference ratios are indicative of a female-biased admixture for that ancestry.

### Uniparental markers

Barcoded primers [48](in preparation) were used to amplify the full mitochondrial sequences from 72 SAC individuals. Using a uniquely barcoded primer combination for every sample, we performed a PCR to amplify the whole mitochondrial genome (30x (98 ^*o*^C, 10 sec; 67 ^*o*^C, 15 min); 4 ^*o*^C) (300 ng DNA, 2.4 nM primers, 200 μM of each dNTP, 1x PCR buffer and 1.25 U Takara LA Taq polymerase in 25 μl reaction). Specificity of PCR products was confirmed on a 1% agarose gel and purified with AMPure PB beads.

Concentrations of the cleaned PCR products were measured (Qubit). Samples were pooled (100 ng/sample). The pool was purified with 0.5x volumes AMPure PB beads. Elution was performed in 10 mM Tris-HCl, pH 8.5. Concentration of the cleaned pool was measured on the Qubit. The full mitochondrial genomes were sequenced on the PacBio Sequel II. Demultiplexing of the sequencing data was performed by Uppsala Genome Centre (UGC) at NGI-SciLifeLab using the SMRT analysis pipeline (www.pacb.com/products-and-services/analytical-software/smrt-analysis/). The full mtDNA sequence reads were mapped to the Revised Cambridge Reference Sequence (rCRS, NCBI accession number: NC 012920.1) to create BAM files. These were converted to FASTA files using DeepVariant (version 1.3.0, settings: –model type=PACBIO) and bcftools consensus (version 1.12). Mitochondrial haplogroups were assigned using HaploGrep3 [49]. All haplogroups were associated with an ancestry, according to literature (see Supplementary Table 4).

Y chromosomal haplogroups were assigned for all 119 males using SNAPPY [50] on the genotyping array data and all haplogroups were associated with an ancestry, according to literature (see Supplementary Table 5).

## Results

In this study, we aim to provide a comprehensive analysis of the genetic ancestry of the South African Coloured (SAC) population by investigating genome-wide data from 356 (125 new) individuals self-identifying as SAC, coming from 22 (7 new) locations in South Africa (Supplementary Figure 1). Building upon previous genetic research, our investigation encompasses a thorough examination of ancestral genetic components within the SAC, for the first time focusing on geographically dispersed SAC groups. Employing a combination of genomic techniques, including analysis of mitogenomes, Y chromosomes, and X-to-autosomal comparisons, we investigate the complexities of admixture events and explore geographic variations in ancestral contributions. Through these approaches, we seek to elucidate the complex genetic make up of SAC and shed light on the historical and demographic factors that have shaped this diverse population.

### Autosomal ancestry contribution in geographically dispersed SAC groups

We created a database consisting of 356 SAC individuals and 847 reference individuals. To capture the major genetic variation between continental groups and to investigate the affiliations of SAC individuals in this genetic space, we applied principal component analysis (PCA) to our dataset. The first principal component separates the out-of-Africa populations from the Khoe-San and West-African populations, while the second principal component represents the variation between Khoe-San and West African-related ancestries (Figure 1A). The SAC individuals are observed scattered in-between these extremes, with some individuals associating more with either Khoe-San, non-African or West African groups. Moreover, PC3 separates East Asian and European ancestry, with South Asians grouping between these two extremes (Supplementary Figure 2). Certain SAC individuals are off-set towards the Asian extreme, suggesting increased ancestry contributions from Asians. Analyzing the average PC values per population (Supplementary Figure 3) reveals a noticeable west-to-east pattern in the PCA. The western locations District Six, Wellington, Genadendal, and Greyton tend to cluster nearer to European populations, while the eastern locations Graaff-Reinet and Nieu-Bethesda show closer proximity to Khoe-San and West-African/Bantu-speaking groups. The three other new locations, Kranshoek, Oudtshoorn, and Prince Albert, which are located geographically in between the previously mentioned groups, also occupy the space in the PCA plot between these groups. Among the three, Kranshoek is closest to Genadendal and Greyton in the PCA plot.

**Figure 1:**
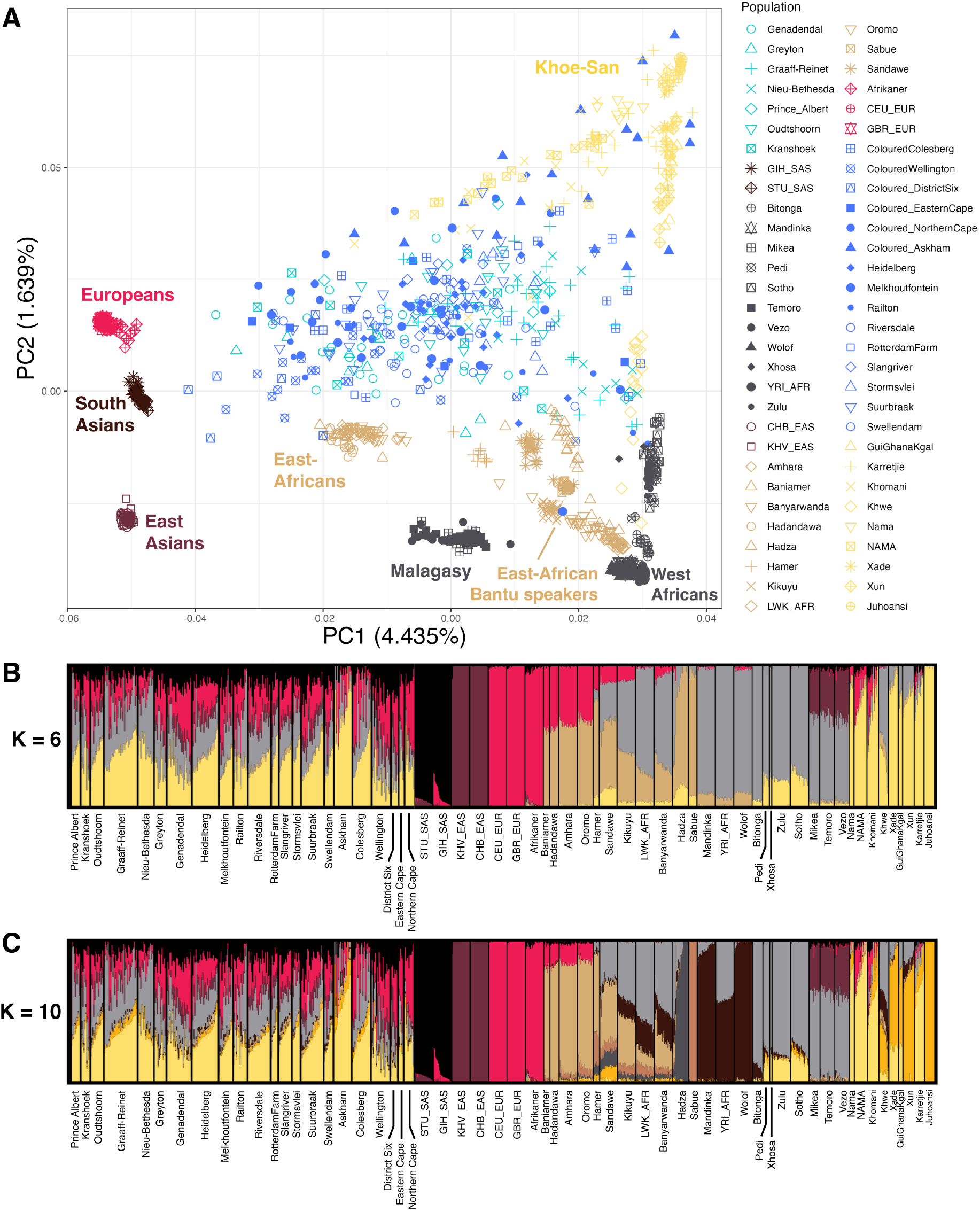
Population structure and genetic affinities of South African Coloured population. Principal component analysis (PCA) and ADMIXTURE results for the populations in our dataset, including 356 SAC individuals. In A, principal component analysis (PCA) results are shown, where PC1 and PC2, are plotted against each other. Labels according to continental groups were added *a posteriori* to help with legibility. The new SAC samples are shown in light blue, the previously published ones in dark blue. For geographical origins of populations, see Supplementary Figure 1. Other PCA projections can be found in Supplementary Figure 2. B and C show ADMIXTURE results, visualized using PONG for K = 6 and K = 10 respectively. ADMIXTURE results for K=2 to K=12 can be found in Supplementary Figure 5. GIH SAS are the Gujarati Indians, STU SAS are the Sri Lankan Tamil, YRI AFR are the Yoruba from Nigeria, CHB EAS are the Han from China, KHV EAS are the Kinh from Vietnam, LWK AFR are the Luhya from Kenya, CEU EUR are Utah residents with Northern and Western European ancestry, and the GBR EUR are the British in England and Scotland.

**Figure 2:**
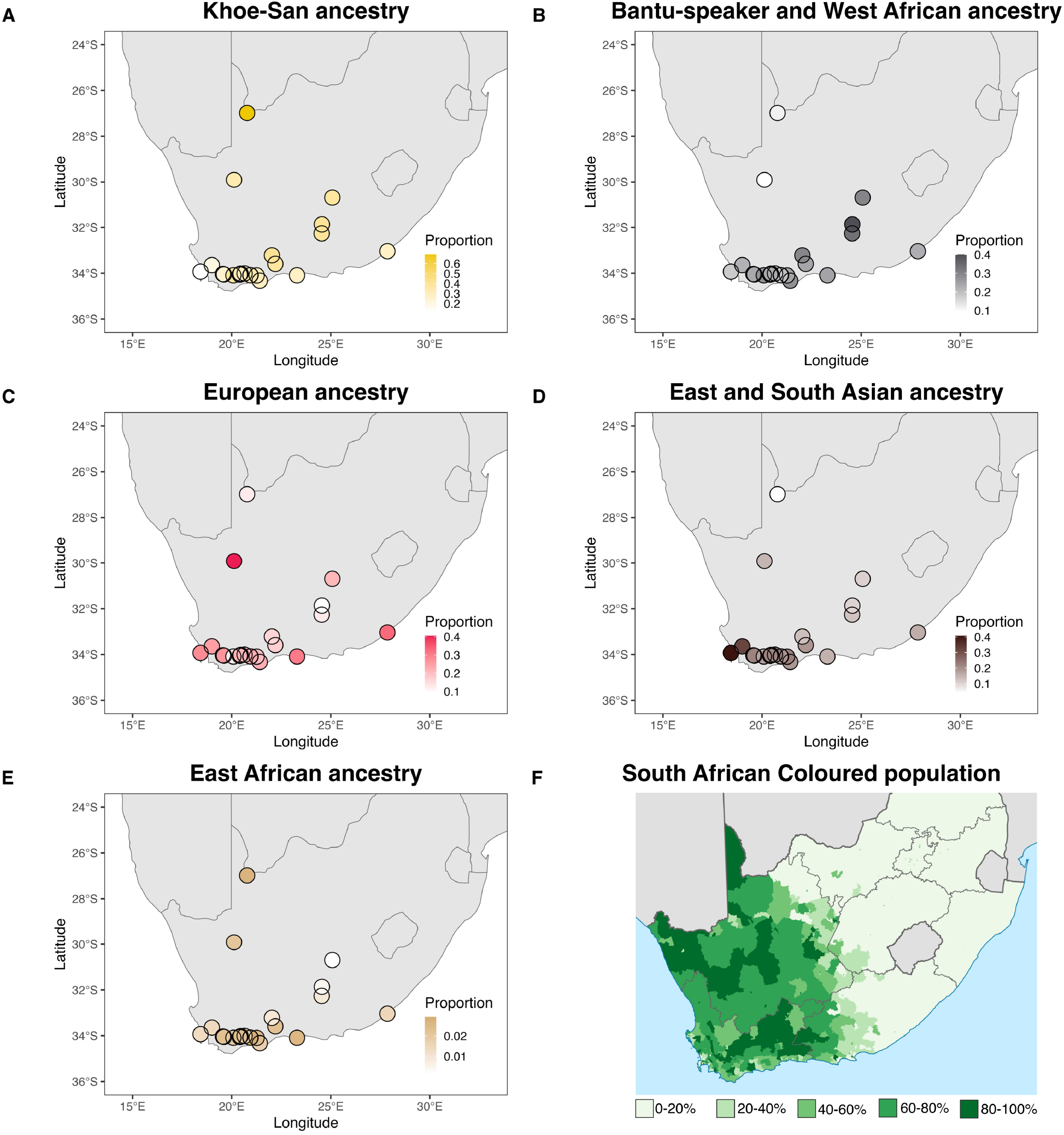
Visualisation of averaged ADMIXTURE derived ancestry proportions from K = 6 plotted by sampling locations. The colour scale is relative to the maximum value of each fraction of admixture. A depicts the component associated to Khoe-San ancestry, the corresponding is shown for B, West African and Bantu-speaker ancestry, C European ancestry, D South Asian and East Asian ancestry combined, and E East African ancestry. In F, the proportion of SAC people among the inhabitants is shown per region in South Africa. Based on the 2011 census. Adapted from https://commons.wikimedia.org/wiki/File:South_Africa_2011_Coloured_population_proportion_map.svg, Public Domain.

Uniform manifold approximation and projection for dimension reduction (UMAP) identifies the major variation in the data and reduces it down to only two dimensions, thus allowing a graphical overview of the variation [51]. Unlike PCA, UMAP aims to capture more of the global variation [51]. The UMAP analysis recapitulates the major continental ancestries within the dataset, with the more drifted out-of-Africa populations forming tightly clustered groups away from each other (Supplementary Figure 4). Khoe-San, SAC, and Bantu-speaker related ancestry populations form a larger group in the center of the UMAP. Most of the SAC are positioned close to the Khoe-San populations but are drawn towards either the European or Bantu-speaker related ancestry. Some SAC individuals cluster firmly with other populations, rather than with the other SAC individuals. Three individuals from District Six, Northern Cape, and Genadendal are closely associated with the European populations. Three other individuals are associated with South Asians, two from Wellington and one from District Six. Additionally, five SAC individuals from various locations group with the South African Bantu-speaking populations.

To further investigate the population structure and ancestral contributions to the SAC populations, we performed unsupervised ADMIXTURE analysis for K = 2 to K = 12 (Supplementary Figure 5). At K = 6 (Figure 1B), we identified components corresponding to major continental and regional groups: Khoe-San, European, West African/Bantu-speakers, East African, East Asian, and South Asian. Compared to K = 6, K = 10 revealed additional clusters: one associated with the East African Hadza, another associated with the East African Sabue, a cluster separating northern Khoe-San from southern Khoe-San populations, and a cluster separating Bantu-speakers from West African non-Bantu Niger-Congo speakers. K = 10 (Figure 1C) had the lowest cross-validation error, see Supplementary Figure 6. Average admixture fractions at K = 6 are shown in Supplementary Table 2. From the 22 locations with SAC individuals, Khoe-San ancestry is predominant at 14 locations, including the new locations of Graaff-Reinet, Nieu-Bethesda, Kranshoek, Oudtshoorn and Prince Albert. Khoe-San ancestry ranges from 12.0% (District Six) to 69.0% (Askham) across all sites, with an average of 33.4%. Based on K = 10 ADMIXTURE results, we can conclude that this observed Khoe-San ancestry is mostly southern Khoe-San rather than northern Khoe-San (yellow vs gold respectively in Figure 1C). This was confirmed by f4-statistics in the form *f4* -(Chimp, SAC, Ju/’hoansi, Karretjie) (Supplementary Figure 7). European ancestry is predominant in seven locations, including Genadendal and Greyton. Generally, European ancestry ranges between 9.2% (Nieu-Bethesda) and 40.5% (Northern Cape) in the studied SAC populations, with an average of 21.7%. In Railton, West-African ancestry constituted the largest proportion (32.8%) while the West-African ancestry was lowest in Northern Cape (9.4%).

At K = 9, ADMIXTURE analysis separates the West African ancestral component into a cluster maximised in West African non-Bantu Niger-Congo speakers (dark-brown) and another in Bantu-speaking populations (light grey)(Supplementary Figure 5). From this K and higher, the component found among the SAC is mostly related to Bantu-speaker ancestry rather than non-Bantu Niger-Congo speakers. In addition, to directly evaluate the genetic affinity of West African/Bantu-speaker ancestry found in SAC, we conducted the test *f4* (Chimp, SAC, YRI AFR, Zulu) (Supplementary Figure 8). All SAC groups, except the Coloured from Askham exhibit greater genetic affinity to the Yorubans relative to the South African Bantu-speaking Zulu.

F4-statistics were computed to assess the genetic affinity of the Asian component in the SAC population (*f4* (Chimp, SAC, CHB EAS, GIH SAS)), where CHB EAS are the Han Chinese (East Asian) and GIH SAS are the Gujarati Indians (South Asians) (Supplementary Figure 9). Positive values for most SAC groups imply more genetic affinity with South Asians rather than East Asians. The ADMIXTURE results also highlight an additional interesting aspect about the ancestry of the studied SAC populations, namely the presence of a genetic component shared with the Malagasy populations. From K = 4 to K = 10, the genetics of the Malagasy populations (Mikea, Temoro, Vezo) can be explained as being comprised mainly of two clusters; a West African cluster (grey) (∼ 60%), and a East Asian cluster (∼ 40%). However, from K = 11, the Malagasy populations get their own cluster (royal blue), with some minor West African and East Asian contributions. This genetic cluster can also be observed in the various SAC populations, at low percentages. The average percentage of this component across all the studied SAC locations is 5.8%. Positive values for f4-statistics in the form *f4* (Chimp, SAC, Malagasy, South Asian) (Supplementary Figure 10) for all SAC groups imply they possess greater genetic affinity with South Asians relative to Malagasy people.

As the ADMIXTURE at K = 6 captures best the diversity in ancestries in the SAC, the average ADMIXTURE derived ancestral fractions for K = 6 were plotted on a map of the southern part of South Africa, to investigate spatial patterns of the different major ancestries (Figure 2). Khoe-San ancestry is larger towards the inland regions and towards the east, while Bantu-speaker ancestry proportions are higher in the most eastern localities. The combined East and South Asian ancestry is highest close to Cape Town and decreases with increasing distance. East African ancestry is smaller than the other ancestries (0.1-2.9%), but geographical differences can be observed with higher East African ancestries along coastal regions and along the Gariep river valley (northern-most point). The European-related ancestry is highest along the coast, with the exception of the Northern Cape site.

### Admixture dating

SAC individuals trace their ancestry to major ancestral groups that might have admixed during different time periods. We employed a local ancestry estimation method to discern the mosaic composition of the genomes of SAC individuals, delineating which segments most likely originated from each parental population. We employed a 5-way admixture model in MOSAIC allowing for all reference populations as parental populations, meaning that MOSAIC uses the reference populations to construct five ancestries that best describe the haplotypes observed in the target. Each constructed ancestry is compared to the source populations through F_st_. The constructed ancestries typically reflect the major continental ancestries that we get from ADMIXTURE, see the 1-F_st_ plots in Supplementary Figures 14 to 57. However, in some cases several ancestries belong to the same continental ancestry. One such case is Oudtshoorn, where the fourth and fifth ancestries are closest to the Khomani and Karretjie respectively, both southern Khoe-San groups.

Subsequently, using this information, we retrieved the admixture dates of these parental populations from the co-ancestry curves as generated by MOSAIC (Figure 3). Most of the dates fall within less than ten generations, overlapping with the time period since European colonisation (1650 onward). Nine SAC populations display admixture dates that are above 50 generations ago (corresponding to 1450 years, assuming a generation time of 29 years [52]). Seven of these older admixture dates are associated with Khoe-San and East-African ancestry, two of them with European and South Asian ancestry.

**Figure 3:**
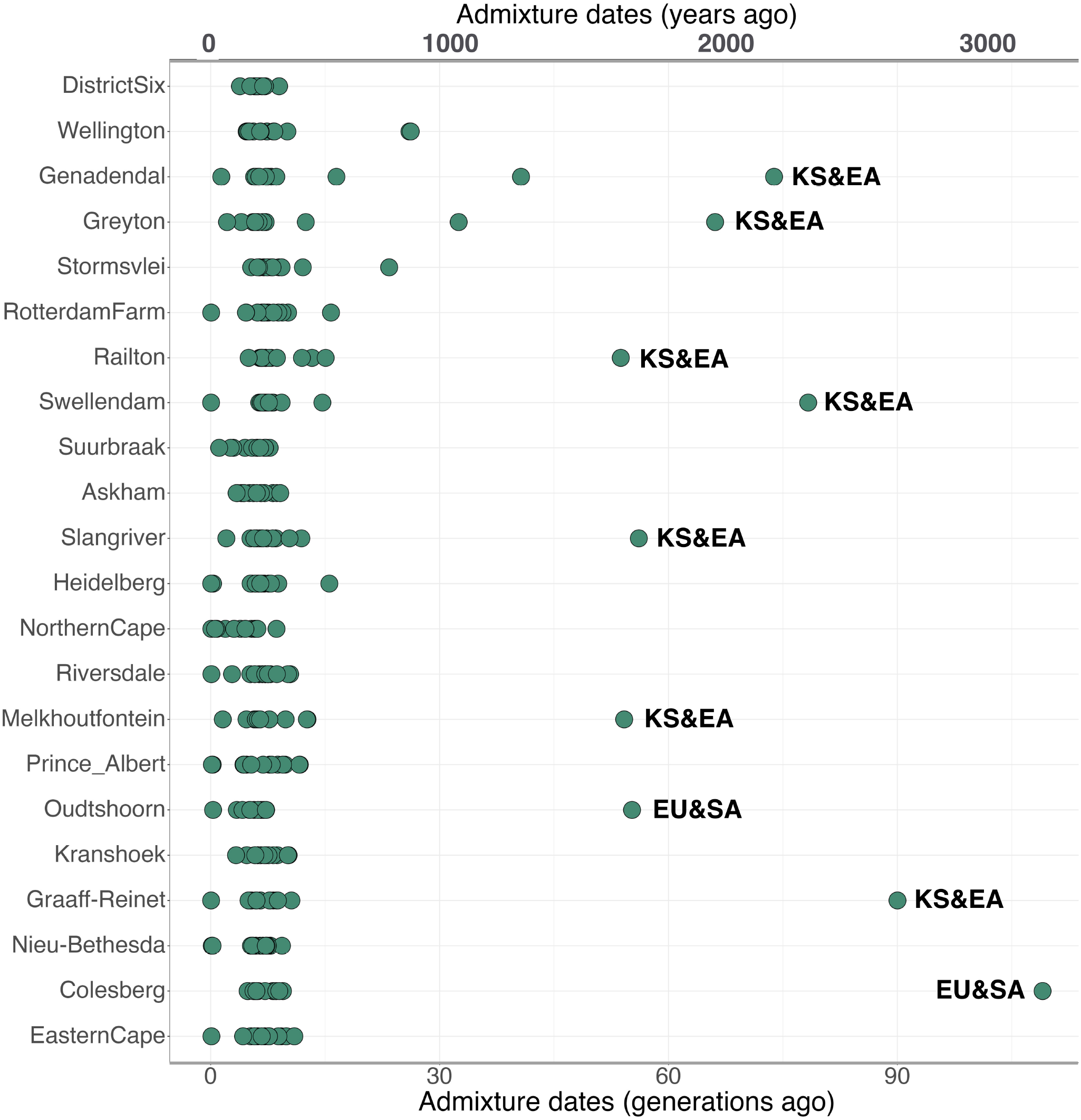
Inferred admixture dates for the 5-way admixture scenario for the 22 SAC populations using all reference populations as putative sources. Dots labeled with ”EA & KS” indicate admixture events between Khoe-San and East African constructed ancestries, as determined by F_st_. Dots labeled with ”EU & SA” indicate admixture events between European and South Asian constructed ancestries. Sites are shown from West (high) to East (low) on the y-axis. X-axis on top shows the time in years, x-axis at the bottom shows time in generations ago.

### Patterns of sex-biased admixture

Previous studies have shown that the admixture events that shaped the SAC population were sex-biased [25, 11], indicating that the extent of male and female genetic contribution from different admixing populations may have varied. Here, we investigate the sex-biased nature of the admixture events shaping the SAC populations further by performing supervised ADMIXTURE for the autosomes and X-chromosome (Supplementary Figure 11) and looking at the t::Admix ratios of East-African, European, Khoe-San, Asian and West-African ancestry (Figure 4A). Negative X-to-autosomal t::Admix ratios are indicative of male-biased admixture for that ancestry, positive t::Admix ratios are indicative of a female-biased admixture for that ancestry. We observe negative t::Admix ratio values with 95% confidence intervals not overlapping zero for East-African and European ancestries (−0.0177 and -0.0259 respectively), indicating male-biased admixture from East-Africans and Europeans. We observe a positive t::Admix ratio with 95% confidence intervals not overlapping zero for Khoe-San ancestry (0.0365), indicating female-biased ancestry from Khoe-San people. For both Asian and West African ancestries, 95% confidence intervals overlap zero and are therefore not significantly differing from zero, thus indicating non-significant sex-biased admixture from these ancestries. The same analysis was also performed per site (Supplementary Figure 12), and although all Δ Admix ratios associated with European ancestry are negative and all those associated with Khoe-San ancestry are positive, the 95% confidence intervals often overlap zero, due to smaller sample sizes.

**Figure 4:**
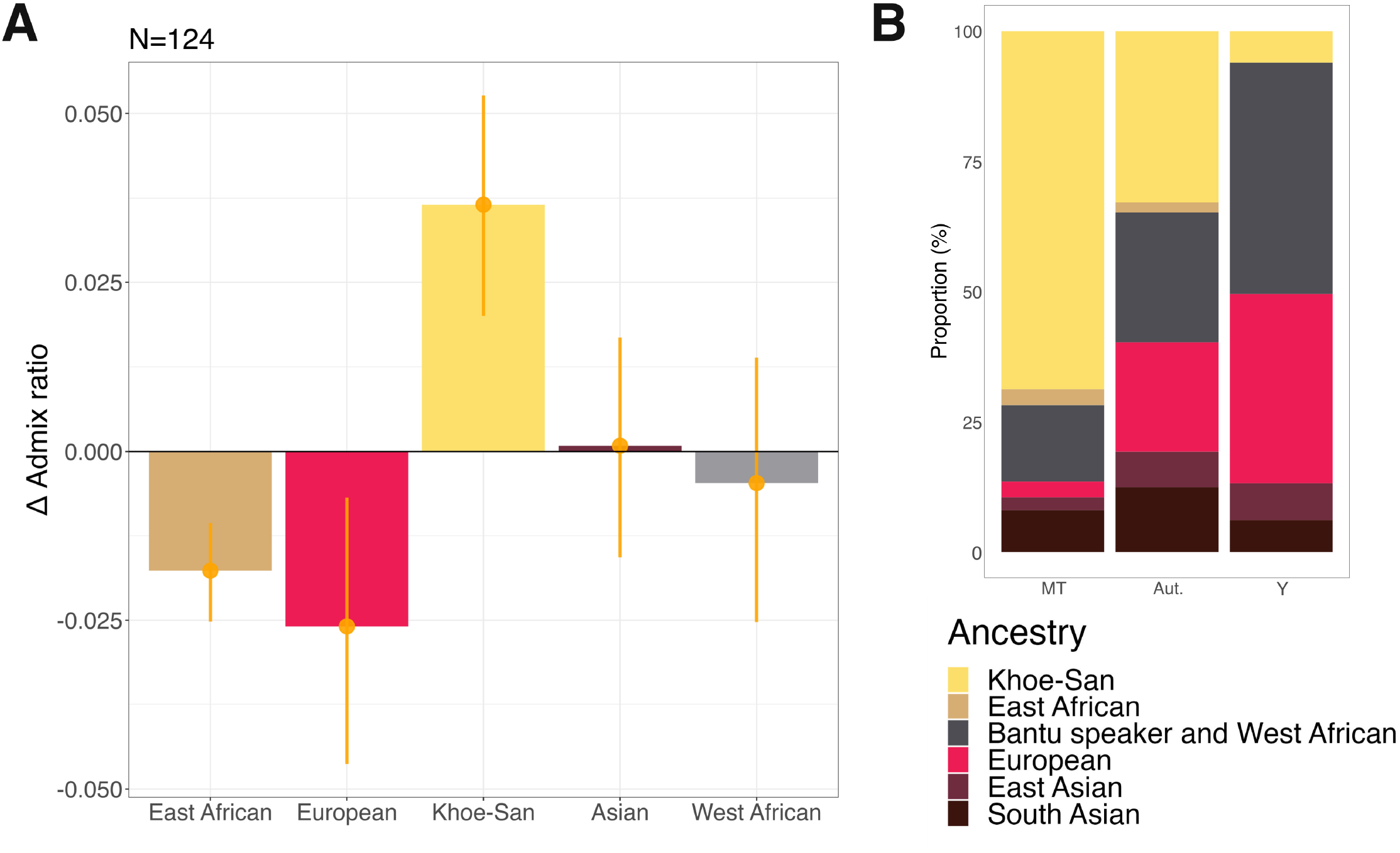
Average sex-biased admixture among the SAC people. In A, t::Admix ratios for each of the five ancestries, averaged over the seven investigated sites are shown. X and autosomal proportions were bootstrapped (10 000 times) and average X-to-autosomal difference ratios were calculated for each of the five ancestries, as well as standard deviations. The error bars indicate the 95% confidence interval. Negative X-to-autosomal difference ratios are indicative of male-biased admixture for that ancestry, positive X-to-autosomal difference ratios are indicative of a female-biased admixture for that ancestry. Results from sex-biased admixture analyses per site can be found in Supplementary Figure 12. In B) ancestries associated with the mitochondrial and Y genomes, as well as the autosomal proportions are shown for all studied SAC individuals.

These signals of sex-biased admixture are further supported with data from the uni-parental markers of these individuals. The mitochondrial genome and the Y-chromosome allow for the study of maternal and paternal lineages separately in a population. We generated novel mitochondrial DNA sequences for 72 SAC individuals and determined the Y-chromosome haplogroups for 67 newly genotyped SAC individuals using SNAPPY [50]. We combined these data with the individuals from the reference datasets. Our results show that mitochondrial haplogroups associated with Khoe-San ancestry are more frequent in the SAC populations than the Y chromosome haplogroups associated to Khoe-San ancestry (Figure 4B). The opposite pattern is observed for West-African and European associated mitochondrial and Y chromosome haplogroups. Haplogroups associated with East-African ancestry become less frequent as we move from mitochondrial genomes to Y chromosomes, but fractions are low (less than 0.031). For both Asian ancestries, no clear pattern can be observed. Supplementary Figure 13 shows the associations of the mitochondrial genomes, autosomes and Y chromosomes to the six different ancestries for each of the separate sites. Large differences in continental distributions can be observed between sites such as Genadendal, Graaff-Reinet and Askham (numbers of indiviudals used for each analysis can be found in Supplementary Table 8). Genadendal, located in the west, generally shows more European ancestry in autosomes, and more mitochondrial and Y chromosomal haplogroups associated with Europeans. This contrasts with the locations of Graaff-Reinet and Askham, situated more to the east and north, respectively. Elevated Khoe-San ancestry can be observed at Askham for all three genetic markers (MT, autosomes and Y), whereas elevated Bantu-speaker ancestry is evident for all markers in Graaff-Reinet.

## Discussion

In this study, we have analyzed genome-wide data from 125 SAC individuals, coming from seven different locations in South Africa. Combining this information with previously published population genetic data, our investigation encompasses a thorough examination of ancestral genetic components within the SAC, spanning a wide geographic distribution. We have investigated the complexities of admixture events that shaped the SAC population and explored potential geographic variations in ancestral contributions. We find evidence of geographical stratification of genetic ancestries in agreement with historical information.

### Ancestry proportions in the South African Coloured population

Our analysis of the general genetic background of the SAC population through PCA and ADMIXTURE (Figure 1) supports the previously identified ancestral components: Khoe-San, West African and Bantu-speaker, European, East African, South and East Asian [24, 25, 26, 11]. We identified heterogeneity within the SAC, ancestry proportions differing substantially across individuals (Figure 1B and C). Average continental ancestry is generally comparable across sites, albeit with some regional variation.

Results from the ADMIXTURE analysis, Figure 1B & C, align with previous studies [26, 11, 1] indicating that the primary genetic ancestry found among the SAC people is Khoe-San. ADMIXTURE at K = 8 (Supplementary Figure 5) splits the northern Khoe-San from the southern Khoe-San populations, thereby revealing for the first time that the Khoe-San ancestry in the SAC is mostly southern Khoe-San-related (Nama, Karretjie, and Khomani). ADMIXTURE at K = 9 separates the West African ancestral component into a component maximised in West African non-Bantu Niger-Congo speakers (dark brown) and another in Bantu-speaking populations (light grey) (Supplementary Figure 5). The analysis at K = 10 reveals low contribution (0.4-3.3%) of the West African ancestry in the SAC. Slaves were brought to the Cape colony from the West African kingdom of Dahomey and from Angola in 1658, and they were a part of the founding population of the SAC [20]. Thus it is interesting to see that the ADMIXTURE analysis only reports low West African ancestry contributions among the SAC populations, and around 22.5% (minimum 7.6, maximum 39.5) of the Bantu-speaker-associated ancestry. According to the ADMIXTURE analysis, these initial enslaved individuals seem to have contributed a small but consistent amount of ancestry to the SAC communities. However, the f4-statistics in the form *f4* (Chimp, SAC, YRI AFR, Zulu) indicate a greater genetic influence from the West-African Yoruba (Supplementary Figure 8). This mismatch with the ADMIXTURE results has been observed in other studies as well, and is called “neighbour repulsion” [53], where the neighbouring populations (Zulu in this case) received independent gene flow from an external source after their split from the West-Africans. In Afrikaners, the West African non-Bantu Niger-Congo speaker associated component contributes more than the Bantu-speaker associated component [19]. This difference in ancestry contributions based on ADMIXTURE likely reflects different patterns of historical admixture for the SAC and Afrikaner populations. West African admixture into Afrikaners likely occurred during the early phases of founding of the colony, with slaves of West African origin, while most of the West African component in the SAC groups was most likely contributed through continued admixture with Bantu-speakers in the contact zone towards the east.

District Six and Wellington have relatively high South Asian ancestry contributions (Supplementary Table 2), likely due to the specific social dynamics at these sites. The District Six community was formed by formerly enslaved people, merchants, and immigrants. Cape Malays, brought as part of the slave trade, composed an essential portion of the founding community, along with the Xhosa people. Afrikaners composed only a small part of the residents of District Six until apartheid laws declared it a ”whites-only” area in 1966, causing many people to be forcibly relocated [54]. Today, more than 90% of its inhabitants are SAC [55]. Similarly to District Six, Wellington was founded in 1699 as an agricultural town. Until the first part of the 20th century, it was mainly composed of SAC residents, many of whom were Muslims and of Asian descent [56].

Our ADMIXTURE analysis at K = 6 also highlights the genetic contributions of Asian populations to the SAC population. With the average South Asian contribution at 12.1%, it is roughly twice as large as the contribution from East Asians. F4-values in the form *f4* (Chimp, SAC, CHB EAS, GIH SAS) are positive for most SAC groups, supporting more admixture from South Asians (Supplementary Figure 9). Thus, we conclude that most of the Asian slaves were brought from South Asia, and to a lesser extent from East Asia. This corresponds to the historical record stating that the Dutch East India Company imported slaves from Indonesia to South Africa [57].

Through ADMIXTURE and subsequent f4-statistics, we also elucidate for the first time the contribution of Malagasy populations to the SAC population. At K = 11, the Malagasy populations get their own cluster (royal blue), with some minor West-African and East-Asian contributions (Supplementary Figure 5). This Malagasy population genetic cluster can also be observed in the various SAC populations, with an average percentage of across all the studied SAC locations of 5.8%. We computed f4-statistics in the form *f4* (Chimp, SAC, Vezo, GIH SAS)(Supplementary Figure 10) and show that there is less genetic affinity of the SAC to the Malagasy, when compared to the South Asian population. This is not to say that no admixture occurred with Malagasy people, just that South Asians have made a larger contribution compared to the Malagasy contribution. The Malagasy ancestry found in the SAC population is also consistent with the historical record that the Dutch East India Company imported slaves from Madagascar to South Africa [57].

### Regional differences in observed ancestry proportions

We collected data from seven new locations to further identify regional variations in SAC ancestries. The ancestry proportions at K = 6 on a map of South Africa (Figure 2) reveal various interesting trends. The Bantu-speaker ancestry shows higher contributions in the East, and lower contributions in the West. This can largely be attributed to the dominant presence of Bantu-speaking groups in the eastern regions of South Africa, which marks the historical limit of the Bantu expansion [20]. The high prevalence of Khoe-San ancestry in the SAC in the inland regions and toward the east reflects the influence of the Cape colony and the increased admixture from Europeans and enslaved people from Asia and Madagascar in the areas closer to the coast and to Cape Town. In the Northern Cape region, Khoe-San ancestry is high (33.4-69.0%), and Bantu-speaker and West African ancestry is low (9.4-11.6%). The SAC of the Northern Cape can be traced back to the Nama herder groups who resided in Namaqualand (South Africa) and Namibia, local San hunter-gatherer groups, and to European settlers who moved into these interior areas. Thus, the Nama people likely contributed to the high Khoe-San genetic ancestry in the SAC individuals in the Northern Cape. This is supported by MOSAIC analyses, which find low F_st_ values for the Nama as a source population for the SAC at Askham and Northern Cape (Supplementary Figures 14 and 22). The low Bantu-speaker (and West African ancestry) component in Askham and the Northern Cape site points to limited admixture with Bantu-speakers. The distribution pattern for combined Asian ancestries and, to a large extent, European ancestry exhibits an interesting contrast. In the Cape region, high contributions from Asian and European ancestries can be observed, gradually decreasing as one moves eastwards. This phenomenon finds its roots in the historical influx of European settlers into the Cape Colony, with its centre and entry point at the Cape of Good Hope (current-day Cape Town), accompanied by enslaved people from Asia and other parts of Africa. Although East African ancestry proportions are very low in comparison to the other ancestries, it is higher along coastal regions and along the Gariep river valley (northern-most point) correlating with the past distribution of Khoekhoe herder groups (vs. San hunter-gatherer groups) [8, 58].

### Dating admixture events in the SAC population

Since the SAC individuals trace their ancestries to various continental and sub-continental sources, we set out to investigate when these populations admixed. We identify that most of the admixture dates fall within less than ten generations, aligning with the anticipated timeframe for the formation of the Cape Colony. The admixture dating using MOSAIC (Figure 3) also identified several admixture events that can be correlated with the formation of the Khoekhoe with the arrival of East African pastoralists in southern Africa [59, 60, 61, 11]. This can be seen in a few dates that are very old (longer than 50 generations ago). These exact dates should, however, be viewed with caution as they are based on small fractions of ancestry, have deep time estimates and have parental source groups that might be distant from actual source groups. This uncertainty is reflected in the co-ancestry graphs that the dates are inferred from, in which the Khoe-San vs. East African admixture estimates are the least robust of the analyses, Supplementary Figures 14 to 57. The East African ancestry contribution is the smallest according to the ADMIXTURE results (Supplementary Table 2) and minor ancestries are problematic for the proper fitting of co-ancestry curves. Two of the admixture dates older than 50 generations ago can be attributed to European and South Asian ancestries. This reflects admixture events happening outside the African continent before these ancestries were introduced during colonial times, possibly related to Eurasian trade routes such as the Silk Road (200 BCE - 1450 AD).

The admixture events with Bantu-speakers (unlabeled in Figure 3) mostly occurred during and after colonial times. This indicates that most of the admixture between Khoe-San and Bantu-speakers also occurred after colonial times, due to the disruptions and population mobility that the colonial times instigated. Even the admixture events between Asian and West-African/Bantu-speaker ancestries are all between 3.9 and 11.7 generations ago, also corresponding to the colonial period. Malagasy populations are known to be the result of an admixture event between Austronesian and Bantu sources around 20 to 32 generations ago [35]. These sources are supported by the ADMIXTURE analysis in this study (Figure 1B&C). However, we do not observe the same generation time-frame for the admixture event between Asian and West-African/Bantu-speaker ancestries in the SAC individuals, possibly indicating that most of these ancestries came from Asian, West-Africans, and Bantu-speakers directly, and not from Malagasy populations. This observation fits with the small Malagasy contributions observed at K = 11 (Supplementary Figure 5) and is in line with what has been observed in other SAC populations [11].

### Sex-biased nature of admixture events in the Coloured

From historical records, we know that there were disproportionately few women among the European settlers, especially in the period before 1688 [20]. A previous genetic study concluded that Khoekhoe women constituted the majority of the maternal contribution for the SAC groups [25]. Moreover, additional investigations into the sex-biased nature of the admixture events shaping the SAC using X-chromosome and autosomes inferred a male-biased influence from East Africans, Asians, and Europeans, and a female-bias from Khoe-San and West-African individuals [11]. In the current study, we also observe a male-biased admixture from East Africans and Europeans, and a female-biased admixture from Khoe-San. The Asian ancestry shows a very small female bias and the West-African ancestry a male bias. However, both of these trends are not statistically significant. The investigation of sex-biased patterns at individual sites (Supplementary Figure 12) highlights the heterogeneous nature of the SAC population. Although non-significant, West-African sex-biased admixture ratios are female-biased in some sites (Genadendal, Greyton, Oudtshoorn, and Kranshoek), while being male-biased in other, more northeastern sites (Graaff-Reinet, Nieu-Bethesda and Prince Albert). The sex-biased admixture in the SAC is supported by the findings from the uni-parental markers; mitochondrial genomes and Y chromosomes. Mitochondrial haplogroups associated with Khoe-San ancestry are more frequent in the SAC populations than the Y chromosome haplogroups associated to Khoe-San ancestry (Figure 4B). The opposite pattern can be observed for West-African and European associated mitochondrial and Y chromosome haplogroups. Since the mitochondrial genome is inherited through the female line and Y chromosomes completely through the male line, we observe them as the extremes when it comes to differences between the contribution of the two sexes, whereas the autosomal ancestries are observed somewhere in the middle of these two. We also note the regional variation between the male and female contributions from different populations across the sites (Supplementary Figure 13), again highlighting the genetic heterogeneity of the SAC population.

## Conclusions

In this study, we analyzed new genotype array data for 125 South African Coloured individuals and built upon research to describe the genetics of one the most admixed populations in the world, the SAC. The Khoe-San people, especially the southern Khoe-San, played a major role in the foundation of the SAC, with their ancestry contribution ranging from 12.0-69.0% across all investigated sites. We also identified a considerable variation in ancestry contributions between different individuals. By adding genetic data from seven new geographically dispersed sites, we were able to better investigate geographical differentiation in ancestry proportions and we identified higher Khoe-San contribution in inland regions and toward the east, and higher Bantu-speaker contributions in eastern regions, whereas the Asian ancestry is higher in western regions. Near Cape Town and in the Western Cape province, the non-African ancestry is especially high, reflecting the historically greater density of European colonists and slaves in those locations. We infer that the admixture events shaping the SAC were in many ways sex-biased; mainly female-biased from Khoe-San people and male-biased from both East Africans and Europeans. Altogether, this study highlights the intricate admixture history and diverse ancestry of the SAC population.

## Availability of data and materials

All data generated or analyzed during this study are included in this published article, its supplementary information files and publicly available repositories. The generated genotype data is available for academic research use through the European Genome-Phenome Archive with accession number EGAD50000000513 (152 individuals) and Data Access Committee EGAC50000000240. Scripts are available at https://github.com/imkelankheet/heet/South-African-Coloured-project.

## Acknowledgements

We are grateful to all subjects who participated in this research. We also would like to thank Gustav and Lili Radlov, Chris and Marie Heese, Andrew and Anneke Fraser-Jones, Harry and Belinda Gordon, Dr. Morley, Monica Thomson, Michelle Moodie, Poem Mooney, Dr Judy Maguire, Maria Johnson and the family Simmers for help during field collection. We thank Michael Salter-Townshend for support with MOSAIC-related discussions. We are grateful for Cesar Fortes-lima for his help with the EGA data upload. The genotyping was performed by the SNP&SEQ Technology Platform (Uppsala, Sweden). The facility is part of the National Genomics Infrastructure supported by the Swedish Research Council for Infrastructures and Science for Life Laboratory (NGI-SciLifeLab), Sweden. The SNP&SEQ Technology Platform is also supported by the Knut and Alice Wallenberg Foundation. The computation and data handling were enabled by resources provided by the Swedish National Infrastructure for Computing (SNIC) at Uppmax partially funded by the Swedish Research Council through grant agreement no. 2018-05973. This project was supported by funding to CS from the European Research Council (ERC) under the European Union’s Horizon 2020 research and innovation programme (grant agreement No. 759933), the Knut and Alice Wallenberg foundation, the Leakey foundation and the Erik Philip Sorensson foundation. An authorized NIH Data Access Committee (DAC) granted data access to Carina Schlebusch for the controlled-access genetic data analyzed in this study that were previously deposited by Scheinfeldt et al. (2019) in the NIH dbGAP repository (dbGaP accession code: phs001780.v1.p1; project approval date: 2019-05-17), as well data previously deposited by Martin et al. (2017) in the NIH dbGAP repository (dbGaP accession code: phs001753; project approval date: 2019-10-25).

