## Supplementary for "Wide-scale Geographical Analysis of Genetic Ancestry in the South African Coloured Population"

Supplementary Table 1: List of populations included in the dataset

| Population | Latitude | Longitude | Published in | Language | Subsistence | Geographical area |
|---|---|---|---|---|---|---|
| Amhara | 10 | 39 | [32] | Afro-Asiatic | Agriculturalist | EastAfrica |
| Baniamer | 19.15 | 35.66 | [34] | Afro-Asiatic | Agriculturalist | EastAfrica |
| Banyarwanda | 2.16 | 33.69 | [32] | Niger-Kordofanian | Agriculturalist | EastAfrica |
| Bitonga | -24.25 | 34.78 | [30] | Niger-Kordofanian | Agriculturalist | SouthernAfrica |
| CEU_EUR | 49.89 | 5.05 | [31] | Indo-European | Agriculturalist | Europe |
| CHB_EAS | 39.88 | 116.41 | [31] | NA | NA | EastAsia |
| ColouredColesberg | -30.7 | 25.08 | [1] | Indo-European | Agriculturalist | SouthernAfrica |
| ColouredWellington | -33.64 | 19 | [1] | Indo-European | Agriculturalist | SouthernAfrica |
| Coloured_Askham | -26.98 | 20.78 | [1] | Indo-European | Agriculturalist | SouthernAfrica |
| GBR_EUR | 51.51 | -0.13 | [31] | Indo-European | Agriculturalist | Europe |
| GIH_SAS | 22.75 | 73.09 | [31] | Indo-European | Agriculturalist | SoutheastAsia |
| GuiGhanaKgal | -23.65 | 24.67 | [1] | Khoisan | Hunter-Gatherer | Khoisan |
| Hadandawa | 20.8 | 35.53 | [34] | Afro-Asiatic | Agriculturalist | EastAfrica |
| Hadza | -3.11 | 33.25 | [34] | Khoisan | Agriculturalist | EastAfrica |
| Hamer | 4.84 | 36.52 | [34] | Afro-Asiatic | Agriculturalist | EastAfrica |
| Heidelberg | -34.08 | 20.96 | [11] | Indo-European | NA | SouthernAfrica |
| Ju/'hoansi | -23.54 | 24.18 | [1] | Khoisan | Hunter-Gatherer | SouthernAfrica |
| Karretjie | -30.71 | 25.1 | [1] | Khoisan | Agriculturalist | SouthernAfrica |
| Khomani | -26.94 | 20.66 | [1] | Khoisan | Agriculturalist | SouthernAfrica |
| KHV_EAS | 10.51 | 106.67 | [31] | NA | NA | EastAsia |
| Khwe | -18.2 | 22.16 | [1] | Khoisan | Hunter-Gatherer | SouthernAfrica |
| Kikuyu | -2.06 | 37.68 | [32] | Niger-Kordofanian | Agriculturalist | EastAfrica |
| LWK_AFR | 0.6 | 34.57 | [31] | Niger-Kordofanian | Agriculturalist | EastAfrica |
| Mandinka | 13.32 | -16.16 | [32] | Niger-Kordofanian | Agriculturalist | WestAfrica |
| Melkhoutfontein | -34.33 | 21.42 | [11] | Indo-European | NA | SouthernAfrica |
| Oromo | 9.25 | 36.89 | [32] | Afro-Asiatic | Agriculturalist | EastAfrica |
| Pedi | -23.84 | 29.98 | [17] | Niger-Kordofanian | Agriculturalist | SouthernAfrica |
| Railton | -34.03 | 20.43 | [11] | Indo-European | NA | SouthernAfrica |
| Riversdale | -34.09 | 21.26 | [11] | Indo-European | NA | SouthernAfrica |
| RotterdamFarm | -34.06 | 20.41 | [11] | Indo-European | NA | SouthernAfrica |
| Sabue | 7.1 | 35.55 | [34] | Nilo-Saharan | Agriculturalist | EastAfrica |
| Sandawe | -7.16 | 35.42 | [34] | Khoisan | Agriculturalist | EastAfrica |
| Slangriver | -34.08 | 20.94 | [11] | Indo-European | NA | SouthernAfrica |
| Sotho | -29.35 | 25.38 | [32] | Niger-Kordofanian | Agriculturalist | SouthernAfrica |
| Stormsvlei | -34.09 | 20.09 | [11] | Indo-European | NA | SouthernAfrica |
| STU_SAS | 7.2 | 80.86 | [31] | NA | NA | SoutheastAsia |
| Suurbraak | -34.01 | 20.65 | [11] | Indo-European | NA | SouthernAfrica |
| Swellendam | -34.02 | 20.45 | [11] | Indo-European | NA | SouthernAfrica |
| Wolof | 15.8 | -16.52 | [32] | Niger-Kordofanian | Agriculturalist | WestAfrica |
| Xade | -22.34 | 23.01 | [28] | Khoisan | Hunter-Gatherer | SouthernAfrica |
| Xhosa | -32.72 | 26.88 | [17] | Niger-Kordofanian | Agriculturalist | SouthernAfrica |
| Xun | -14.63 | 17.67 | [1] | Khoisan | Hunter-Gatherer | SouthernAfrica |
| YRI_AFR | 8.01 | 3.98 | [31] | Niger-Kordofanian | Agriculturalist | WestAfrica |
| Zulu | -30.4 | 29.5 | [32] | Niger-Kordofanian | Agriculturalist | SouthernAfrica |
| Afrikaner | -28.82 | 24.99 | [19] | Indo-European | Agriculturalist | SouthernAfrica |
| Nama | -29 | 17.1 | [33] | Khoisan | Pastoralist | SouthernAfrica |
| NAMA | -22.7 | 17.11 | [1] | Khoisan | Pastoralist | SouthernAfrica |
| Temoro | -22.03 | 47.91 | [35] | NA | Agriculturalist | SouthernAfrica |
| Vezo | -23.54 | 43.75 | [35] | NA | Pastoralist | SouthernAfrica |
| Mikea | -21.65 | 43.86 | [35] | NA | Hunter-Gatherer | SouthernAfrica |
| Coloured_NorthernCape | -29.91 | 20.12 | [26] | Indo-European | NA | SouthernAfrica |
| Coloured_EasternCape | -32.04 | 26.86 | [26] | Indo-European | NA | SouthernAfrica |
| Coloured_DistrictSix | -33.93 | 18.43 | [26] | Indo-European | NA | SouthernAfrica |
| Nieu-Bethesda | -31.87 | 24.55 | This study | Indo-European | NA | SouthernAfrica |
| Oudtshoorn | -33.59 | 22.2 | This study | Indo-European | NA | SouthernAfrica |
| Prince Albert | -33.22 | 22.03 | This study | Indo-European | NA | SouthernAfrica |
| Genadendal | -34.03 | 19.56 | This study | Indo-European | NA | SouthernAfrica |
| Graaff-Reinet | -32.26 | 24.54 | This study | Indo-European | NA | SouthernAfrica |
| Greyton | -34.05 | 19.61 | This study | Indo-European | NA | SouthernAfrica |
| Kranshoek | -34.09 | 23.3 | This study | Indo-European | NA | SouthernAfrica |

Supplementary Table 2: Admixture proportions at K = 6 for the 22 SAC populations. Newly investigated sites are denoted in bold.

| Site | European | East African | East Asian | West African | Khoe-San | South Asian |
|---|---|---|---|---|---|---|
| Colesberg | 0.210 | 0.001 | 0.033 | 0.300 | 0.382 | 0.071 |
| DistrictSix | 0.279 | 0.013 | 0.163 | 0.179 | 0.120 | 0.244 |
| EasternCape | 0.326 | 0.014 | 0.058 | 0.227 | 0.272 | 0.099 |
| NorthernCape | 0.405 | 0.020 | 0.046 | 0.094 | 0.334 | 0.097 |
| Wellington | 0.249 | 0.013 | 0.107 | 0.220 | 0.194 | 0.213 |
| Askham | 0.124 | 0.027 | 0.009 | 0.116 | 0.690 | 0.031 |
| **Genadendal** | 0.279 | 0.026 | 0.089 | 0.200 | 0.252 | 0.151 |
| **Graaff-Reinet** | 0.128 | 0.009 | 0.042 | 0.337 | 0.398 | 0.084 |
| **Greyton** | 0.265 | 0.023 | 0.076 | 0.236 | 0.254 | 0.144 |
| Heidelberg | 0.185 | 0.025 | 0.069 | 0.238 | 0.360 | 0.121 |
| **Kranshoek** | 0.302 | 0.025 | 0.049 | 0.244 | 0.275 | 0.101 |
| Melkhoutfontein | 0.224 | 0.017 | 0.044 | 0.278 | 0.289 | 0.145 |
| **Nieu-Bethesda** | 0.092 | 0.004 | 0.043 | 0.400 | 0.400 | 0.059 |
| **Oudtshoorn** | 0.170 | 0.023 | 0.077 | 0.241 | 0.373 | 0.113 |
| **Prince Albert** | 0.161 | 0.009 | 0.050 | 0.296 | 0.401 | 0.080 |
| Railton | 0.215 | 0.021 | 0.058 | 0.328 | 0.272 | 0.103 |
| Riversdale | 0.203 | 0.024 | 0.075 | 0.245 | 0.299 | 0.150 |
| RotterdamFarm | 0.160 | 0.020 | 0.097 | 0.216 | 0.377 | 0.127 |
| Slangriver | 0.225 | 0.029 | 0.063 | 0.196 | 0.353 | 0.131 |
| Stormsvlei | 0.125 | 0.020 | 0.058 | 0.280 | 0.398 | 0.115 |
| Suurbraak | 0.223 | 0.018 | 0.087 | 0.178 | 0.343 | 0.148 |
| Swellendam | 0.215 | 0.023 | 0.086 | 0.232 | 0.308 | 0.132 |
| Minimum value | 0.092 | 0.001 | 0.009 | 0.094 | 0.120 | 0.031 |
| Maximum value | 0.405 | 0.029 | 0.163 | 0.400 | 0.690 | 0.244 |
| Average | 0.217 | 0.018 | 0.067 | 0.240 | 0.334 | 0.121 |

Supplementary Table 3: Admixture proportions at K = 10 for the 22 SAC populations. Newly investigated sites are denoted in bold.

| Site | Bantu speaker | Sabue-related | Northern San | European European | East Asian | Hadza--related | Southern San | East African | South Asian | West African |
|---|---|---|---|---|---|---|---|---|---|---|
| Colesberg | 0.293 | 0.001 | 0.061 | 0.200 | 0.033 | 0.002 | 0.328 | 0.003 | 0.065 | 0.009 |
| DistrictSix | 0.148 | 0.002 | 0.011 | 0.273 | 0.163 | 0.006 | 0.109 | 0.011 | 0.240 | 0.033 |
| EasternCape | 0.214 | 0.002 | 0.029 | 0.314 | 0.058 | 0.005 | 0.250 | 0.016 | 0.093 | 0.016 |
| NorthernCape | 0.076 | 0.005 | 0.025 | 0.385 | 0.046 | 0.005 | 0.324 | 0.021 | 0.090 | 0.017 |
| Wellington | 0.198 | 0.007 | 0.014 | 0.240 | 0.107 | 0.002 | 0.182 | 0.011 | 0.208 | 0.025 |
| Askham | 0.097 | 0.004 | 0.165 | 0.096 | 0.009 | 0.002 | 0.552 | 0.026 | 0.026 | 0.019 |
| **Genadendal** | 0.183 | 0.004 | 0.014 | 0.262 | 0.089 | 0.005 | 0.245 | 0.028 | 0.145 | 0.019 |
| **Graaff-Reinet** | 0.331 | 0.004 | 0.048 | 0.114 | 0.042 | 0.002 | 0.358 | 0.009 | 0.078 | 0.009 |
| **Greyton** | 0.220 | 0.005 | 0.021 | 0.249 | 0.076 | 0.006 | 0.237 | 0.025 | 0.139 | 0.018 |
| Heidelberg | 0.222 | 0.007 | 0.031 | 0.163 | 0.069 | 0.004 | 0.340 | 0.025 | 0.115 | 0.018 |
| **Kranshoek** | 0.228 | 0.003 | 0.027 | 0.284 | 0.049 | 0.004 | 0.255 | 0.030 | 0.096 | 0.020 |
| Melkhoutfontein | 0.260 | 0.008 | 0.023 | 0.210 | 0.044 | 0.004 | 0.271 | 0.014 | 0.140 | 0.020 |
| **Nieu-Bethesda** | 0.395 | 0.004 | 0.052 | 0.084 | 0.043 | 0.003 | 0.353 | 0.001 | 0.052 | 0.010 |
| **Oudtshoorn** | 0.224 | 0.006 | 0.038 | 0.149 | 0.077 | 0.004 | 0.346 | 0.024 | 0.108 | 0.019 |
| **Prince Albert** | 0.295 | 0.004 | 0.047 | 0.148 | 0.050 | 0.001 | 0.364 | 0.009 | 0.073 | 0.004 |
| Railton | 0.313 | 0.005 | 0.016 | 0.198 | 0.058 | 0.003 | 0.260 | 0.026 | 0.098 | 0.018 |
| Riversdale | 0.235 | 0.007 | 0.023 | 0.186 | 0.075 | 0.004 | 0.283 | 0.024 | 0.145 | 0.013 |
| RotterdamFarm | 0.208 | 0.005 | 0.023 | 0.140 | 0.097 | 0.006 | 0.367 | 0.019 | 0.120 | 0.010 |
| Slangriver | 0.173 | 0.009 | 0.016 | 0.202 | 0.064 | 0.005 | 0.353 | 0.026 | 0.124 | 0.023 |
| Stormsvlei | 0.270 | 0.006 | 0.035 | 0.105 | 0.058 | 0.003 | 0.375 | 0.021 | 0.110 | 0.011 |
| Suurbraak | 0.160 | 0.004 | 0.031 | 0.202 | 0.087 | 0.003 | 0.324 | 0.023 | 0.142 | 0.018 |
| Swellendam | 0.207 | 0.005 | 0.021 | 0.198 | 0.086 | 0.007 | 0.296 | 0.022 | 0.127 | 0.027 |
| Minimum value | 0.076 | 0.001 | 0.011 | 0.084 | 0.009 | 0.001 | 0.109 | 0.001 | 0.026 | 0.004 |
| Maximum value | 0.395 | 0.009 | 0.165 | 0.385 | 0.163 | 0.007 | 0.552 | 0.030 | 0.240 | 0.033 |
| Average | 0.225 | 0.005 | 0.035 | 0.200 | 0.067 | 0.004 | 0.308 | 0.019 | 0.115 | 0.017 |

Supplementary Table 4: Mitochondrial DNA haplogroup assignment by ancestry

| Haplogroup | Most prevalent ancestry | Reference |
|---|---|---|
| B4a | East Asian | [25] |
| B4b | East Asian | [25] |
| B5b | East Asian | [62] |
| E1a | South Asian | [63] |
| H | European | [25] |
| H1a | European | [25] |
| H1c | European | [25] |
| J1c | European | [64] |
| L0a | Bantu-speaker | [6] |
| L0d | Khoe-San | [65];[66]; |
| L0f | East African | [67] |
| L1b | Bantu-speaker | [68] |
| L1c | Bantu-speaker | [6] |
| L2a | Bantu-speaker | [6] |
| L3 | Bantu-speaker | [69] |
| L3d | Bantu-speaker | [70] |
| L3e | Bantu-speaker | [6] |
| L4b | East African | [71] |
| L5a | East African | [71] |
| M1a | European | [72] |
| M6a | South Asian | [69] |
| M18 | South Asian | [73] |
| M2a | South Asian | [73] |
| M2b | South Asian | [73] |
| M33 | South Asian | [73] |
| M42 | South Asian | [74] |
| M5a | South Asian | [73] |
| U2a | South Asian | [75] |
| U7a | South Asian | [75] |

Supplementary Table 5: Y-chromosome haplogroup assignment by ancestry

| Y-chr haplogroup | Most prevalent ancestry | Reference |
|---|---|---|
| A1b | Khoe-San | [76] |
| A0-T | Khoe-San | [77] |
| B2a | Bantu-speaker and West African | [78] |
| B2b | Bantu-speaker and West African | [76] |
| C | East Asian | [79] |
| E1b | Bantu-speaker and West African | [76] |
| E2b | Bantu-speaker and West African | [76] |
| E2 | Bantu-speaker and West African | [80] |
| G2a | European | [69] |
| G2b | West Asian | [81] |
| H | South Asian | [69] |
| I1 | European | [69] |
| I1a | European | [69] |
| I2 | European | [69] |
| I2a | European | [69] |
| J | European | [69] |
| J2a | European | [69] |
| J2b | European | [69] |
| L1a | South Asian | [82] |
| N1c | East Asian | [69] |
| O1a | East Asian | [69] |
| R1a | European | [69] |
| R1b | European | [83] |
| R2 | South Asian | [69] |
| R2a | South Asian | [69] |
| T | European | [84] |

Supplementary Table 6: Mitochondrial ancestries at the 16 sites for which mitochondrial sequences were available. Newly investigated sites are denoted in bold.

| Site | European | East African | East Asian | West African | Khoe-San | South Asian |
|---|---|---|---|---|---|---|
| Askham | 0 | 0.052 | 0 | 0 | 0.947 | 0 |
| Colesberg | 0 | 0 | 0 | 0.277 | 0.722 | 0 |
| **Genadendal** | 0.154 | 0.077 | 0 | 0.077 | 0.615 | 0.077 |
| **Graaff-Reinet** | 0 | 0 | 0.061 | 0.091 | 0.758 | 0.091 |
| **Greyton** | 0.091 | 0.182 | 0 | 0.182 | 0.545 | 0 |
| Heidelberg | 0 | 0 | 0 | 0 | 0.9 | 0.1 |
| Melkhoutfontein | 0 | 0 | 0 | 0.1 | 0.8 | 0.1 |
| **Nieu-Bethesda** | 0 | 0 | 0 | 0.211 | 0.789 | 0 |
| Railton | 0 | 0 | 0 | 0.375 | 0.5 | 0.125 |
| Riversdale | 0 | 0 | 0.083 | 0.083 | 0.75 | 0.083 |
| RotterdamFarm | 0 | 0.143 | 0 | 0.143 | 0.571 | 0.143 |
| Slangriver | 0 | 0 | 0 | 0.125 | 0.625 | 0.25 |
| Stormsvlei | 0 | 0 | 0 | 0.4 | 0.6 | 0 |
| Suurbraak | 0 | 0.091 | 0.091 | 0.182 | 0.545 | 0.091 |
| Swellendam | 0.182 | 0 | 0 | 0.091 | 0.636 | 0.091 |
| Wellington | 0.1 | 0 | 0.05 | 0.35 | 0.45 | 0.05 |

Supplementary Table 7: Y chromosome ancestries at the 22 sites. Newly investigated sites are denoted in bold.

| Site | European | East African | East Asian | West-African | Khoe-San | South Asian |
|---|---|---|---|---|---|---|
| Askham | 0.166 | 0 | 0 | 0.5 | 0.333 | 0 |
| Colesberg | 0.2 | 0 | 0 | 0.6 | 0.2 | 0 |
| DistrictSix | 0.375 | 0 | 0 | 0.5 | 0 | 0.125 |
| EasternCape | 0.285 | 0 | 0.142 | 0.428 | 0.142 | 0 |
| **Genadendal** | 0.666 | 0 | 0.111 | 0 | 0 | 0.222 |
| **Graaff-Reinet** | 0.181 | 0 | 0.045 | 0.727 | 0 | 0.045 |
| **Greyton** | 1 | 0 | 0 | 0 | 0 | 0 |
| Heidelberg | 0 | 0 | 0 | 0.666 | 0.333 | 0 |
| **Kranshoek** | 0 | 0 | 0.333 | 0.666 | 0 | 0 |
| Melkhoutfontein | 0.25 | 0 | 0 | 0.75 | 0 | 0 |
| **Nieu-Bethesda** | 0.25 | 0 | 0 | 0.375 | 0.375 | 0 |
| NorthernCape | 0.333 | 0 | 0.166 | 0.166 | 0 | 0.333 |
| **Oudtshoorn** | 0.444 | 0 | 0.111 | 0.444 | 0 | 0 |
| **Prince Albert** | 0.111 | 0 | 0 | 0.777 | 0.111 | 0 |
| Railton | 0.5 | 0 | 0 | 0 | 0.5 | 0 |
| Riversdale | 0.6 | 0 | 0 | 0.2 | 0 | 0.2 |
| RotterdamFarm | 0.4 | 0 | 0.2 | 0.2 | 0 | 0.2 |
| Slangriver | 0.5 | 0 | 0 | 0.5 | 0 | 0 |
| Stormsvlei | 0 | 0 | 0 | 0.5 | 0 | 0.5 |
| Suurbraak | 0 | 0 | 1 | 0 | 0 | 0 |
| Swellendam | 0.5 | 0 | 0.125 | 0.375 | 0 | 0 |
| Wellington | 0 | 0 | 0 | 0.333 | 0.333 | 0.333 |

Supplementary Table 8: Number of individuals used at each site to make inferences about mitochondrial, autosomal and Y chromsomal ancestries.

| Site | Mitochondria | Autosomal | Y chromosome |
|---|---|---|---|
| Askham | 18 | 19 | 12 |
| Colesberg | 18 | 20 | 5 |
| District Six | 0 | 8 | 8 |
| Eastern Cape | 0 | 6 | 6 |
| Genadendal | 13 | 26 | 9 |
| Graaff-Reinet | 33 | 35 | 22 |
| Greyton | 11 | 13 | 7 |
| Heidelberg | 10 | 28 | 3 |
| Kranshoek | 0 | 10 | 3 |
| Melkhoutfontein | 10 | 15 | 4 |
| Nieu-Bethesda | 19 | 17 | 8 |
| Northern Cape | 0 | 11 | 6 |
| Oudtshoorn | 0 | 14 | 9 |
| Prince Albert | 0 | 10 | 9 |
| Railton | 8 | 15 | 2 |
| Riversdale | 12 | 24 | 5 |
| RotterdamFarm | 7 | 8 | 5 |
| Slangriver | 8 | 14 | 2 |
| Stormsvlei | 10 | 9 | 2 |
| Suurbraak | 11 | 24 | 1 |
| Swellendam | 11 | 10 | 8 |
| Wellington | 20 | 20 | 3 |
| All 22 sites | 219 | 356 | 139 |

### Supplementary Figures

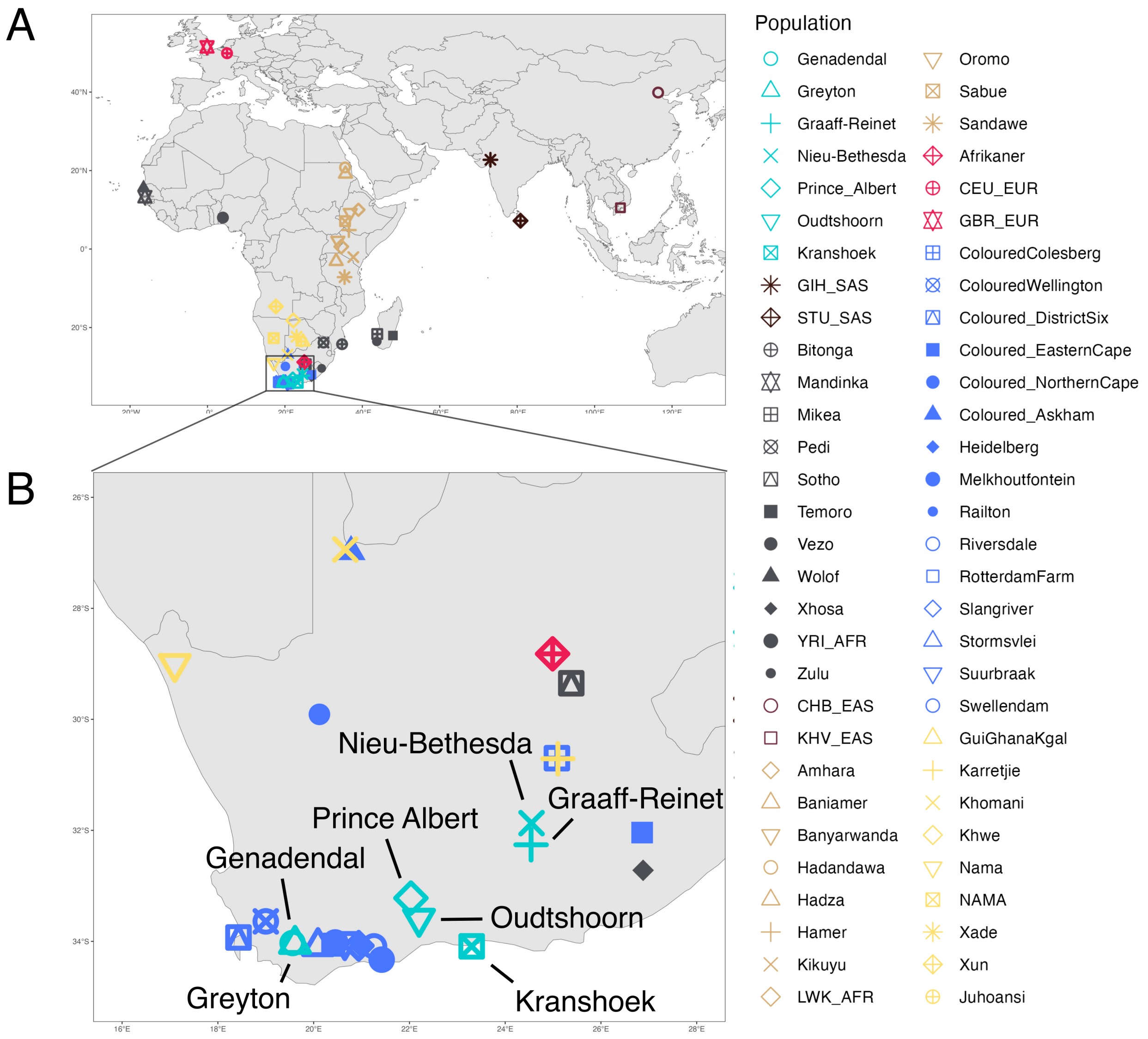


Supplementary Figure 1: Geographic distribution of sampling locations. In A, locations of all populations including reference populations are shown. In B, a zoom-in of the South African region where the new samples are from is shown. The new SAC locations are shown in light blue, the previously published locations in dark blue.

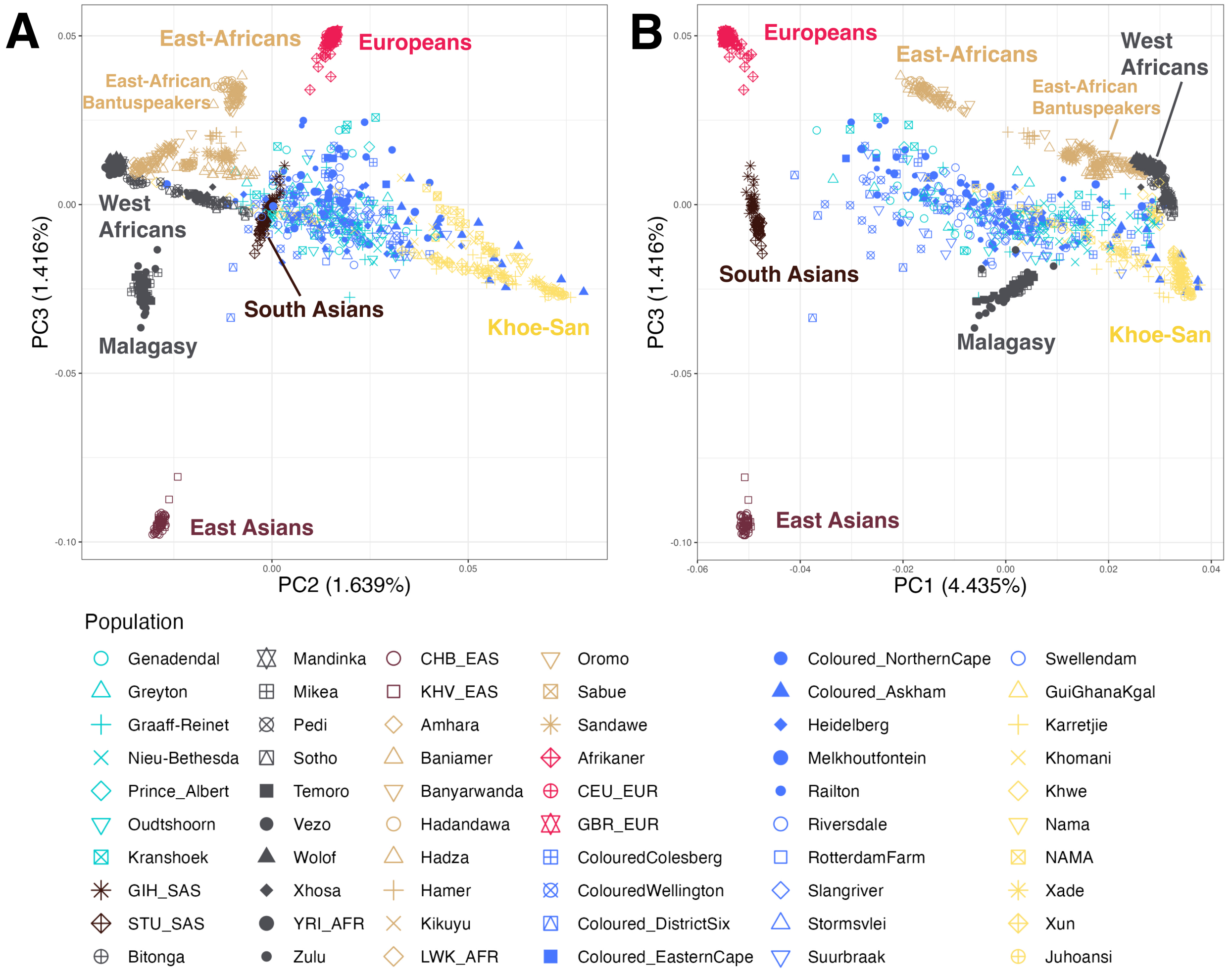


Supplementary Figure 2: Principal Component analysis of the dataset. Within parenthesis is the PC loading. PC3 is plotted against PC2 and PC1, A and B respectively. The new SAC samples are shown in light blue, the previously published SAC samples in dark blue.

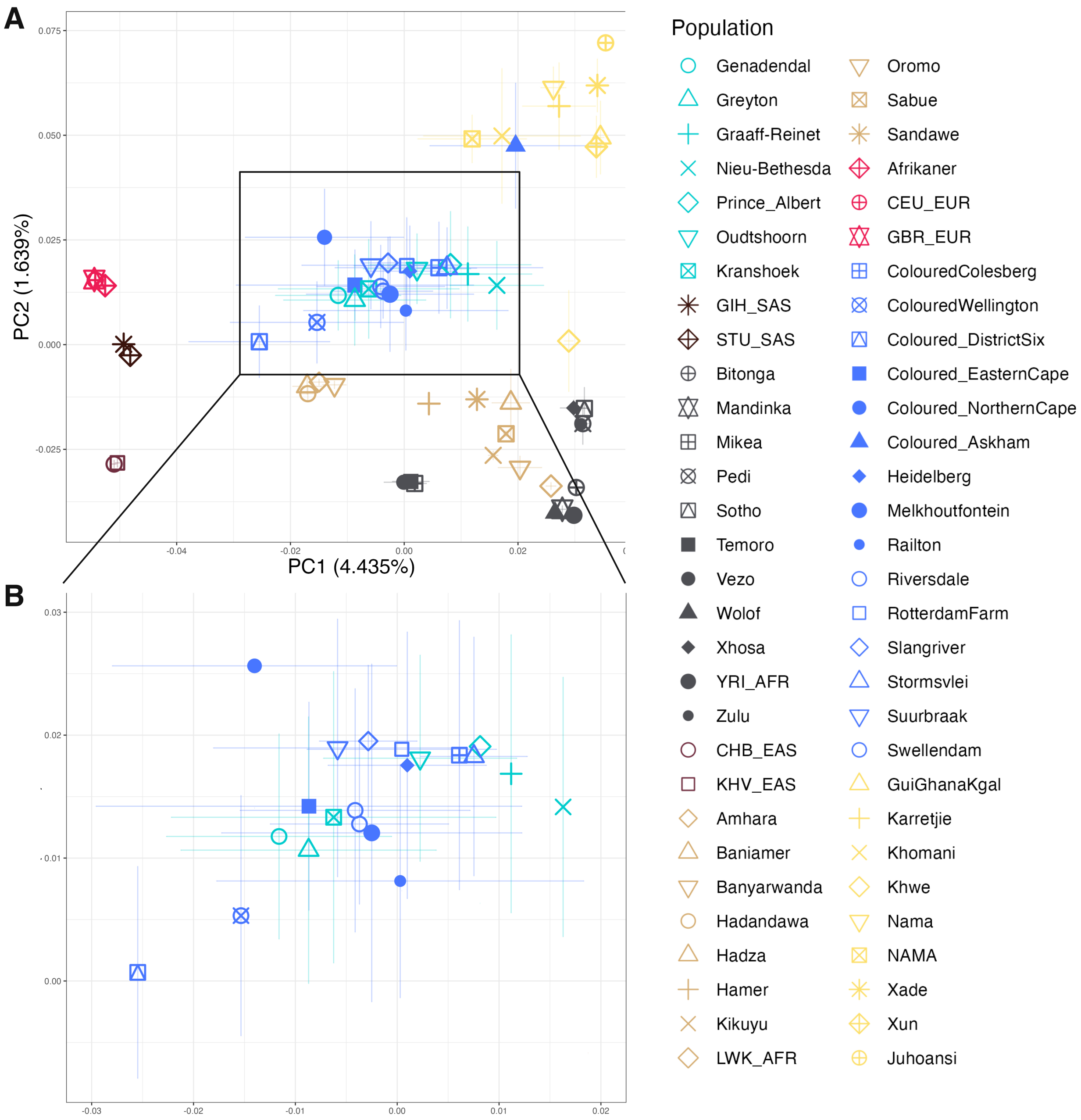


Supplementary Figure 3: Principal Component analysis depicting the average and standard deviations of the PC values of the populations. On the axes, within parenthesis is the PC loading. In B, a zoom in of the plot with the SAC populations is shown. The new SAC locations are shown in light blue, the previously published locations in dark blue.

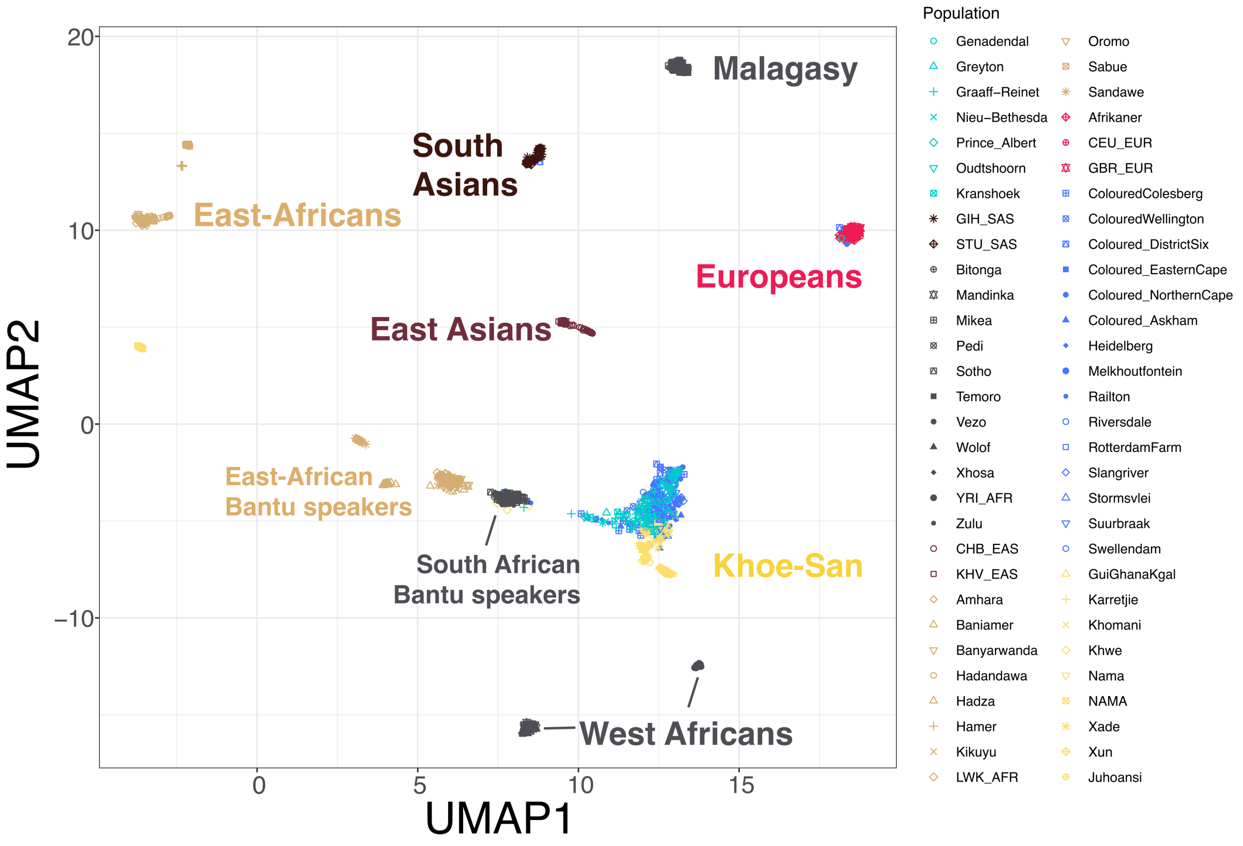


Supplementary Figure 4: Uniform Manifold Approximation and Projection for dimension reduction (UMAP) of the populations in the dataset. Projections are based on genotype calls. Colours indicate continental ancestry. Labels according to continental groups were added *a posteriori* to help with legibility.

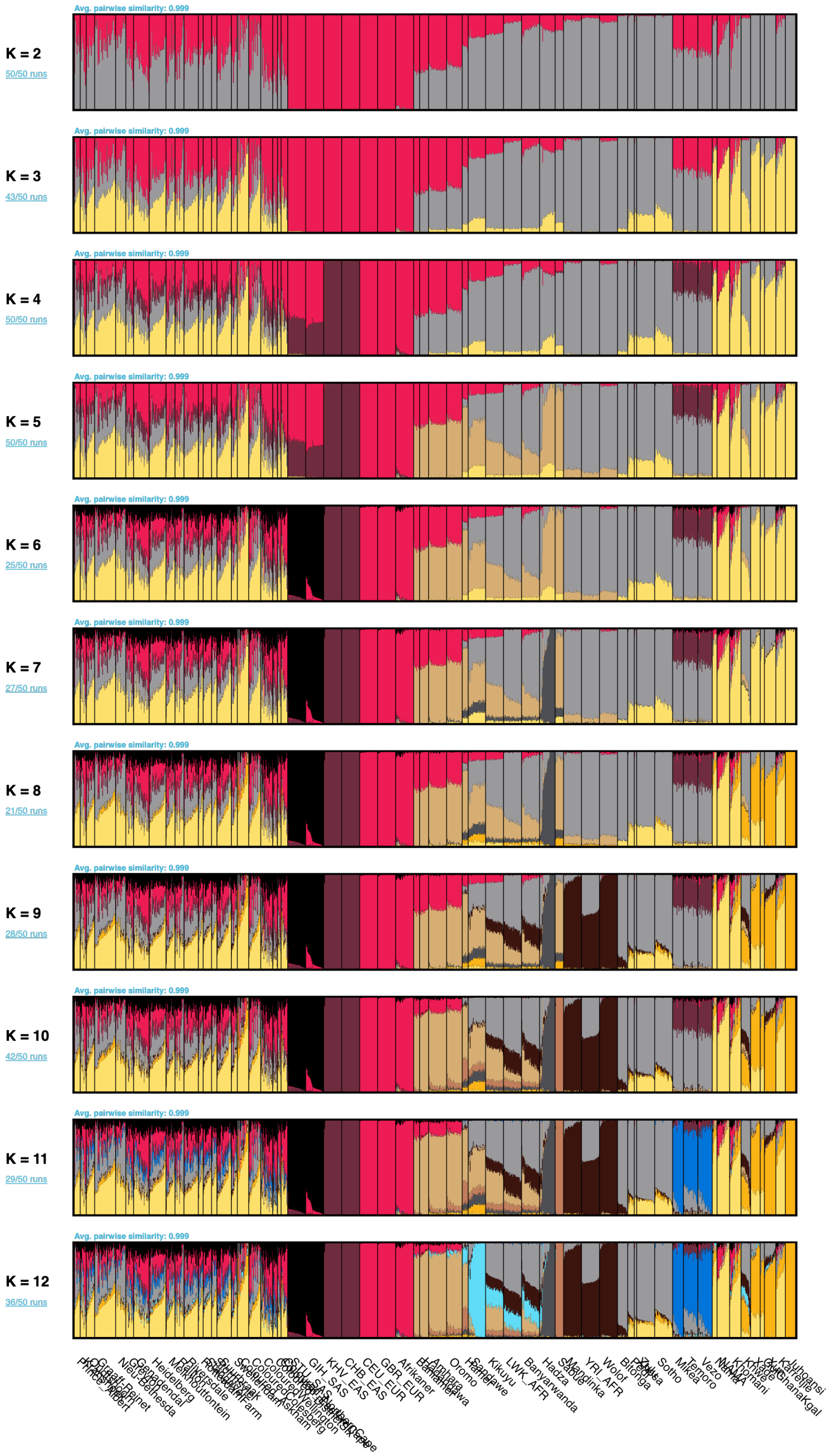


Supplementary Figure 5: PONG visualization of unsupervised ADMIXTURE analysis with 50 iterations for the full dataset, for ancestries K = 2 to K = 12. The best identified K through cross validation was K = 10 (Supplementary Figure 6).

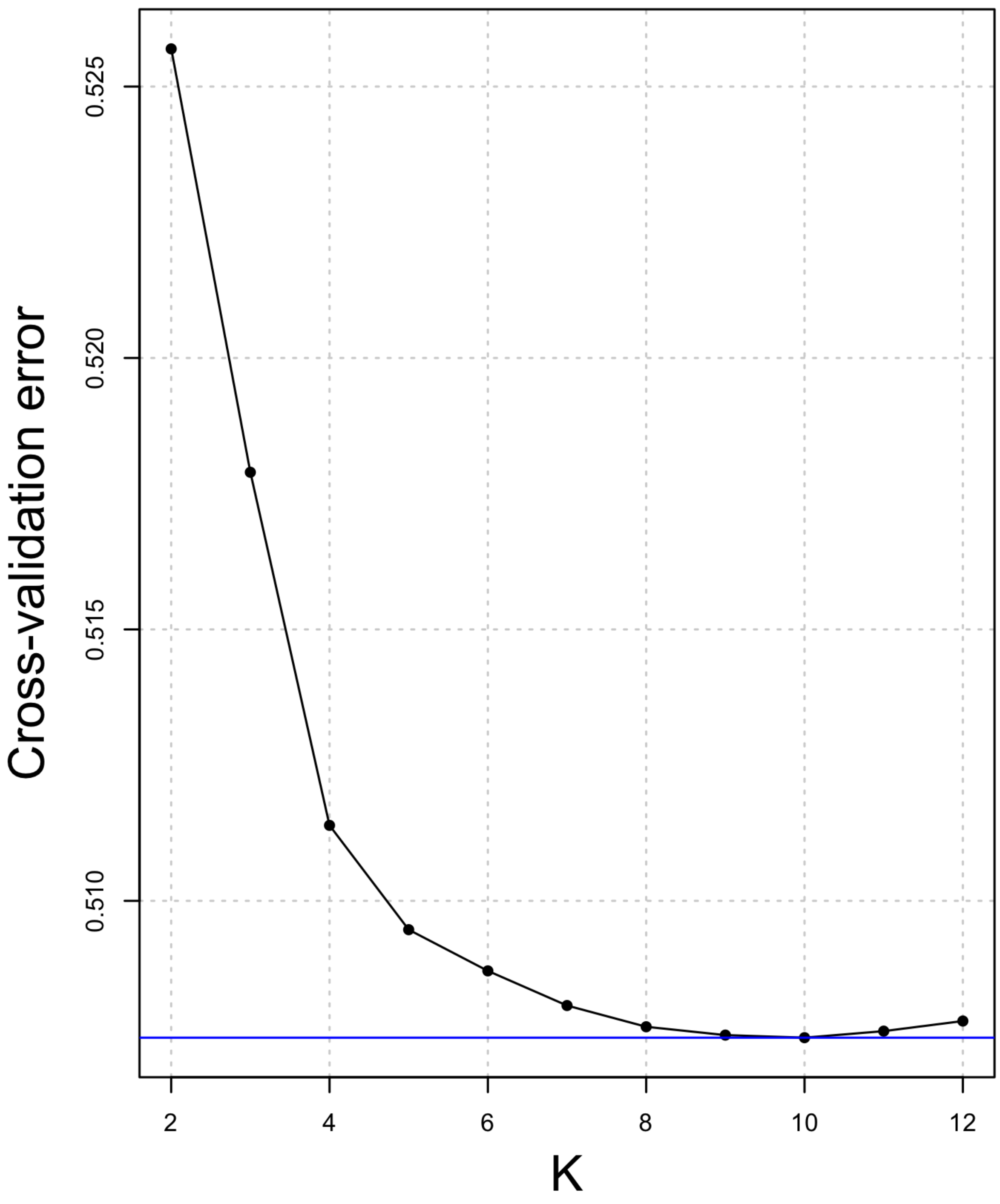


Supplementary Figure 6: Cross validation (CV) error for K = 2 to K =12, averaged over the 50 repetitions. The K with the lowest CV error was K = 10 (horizontal blue line).

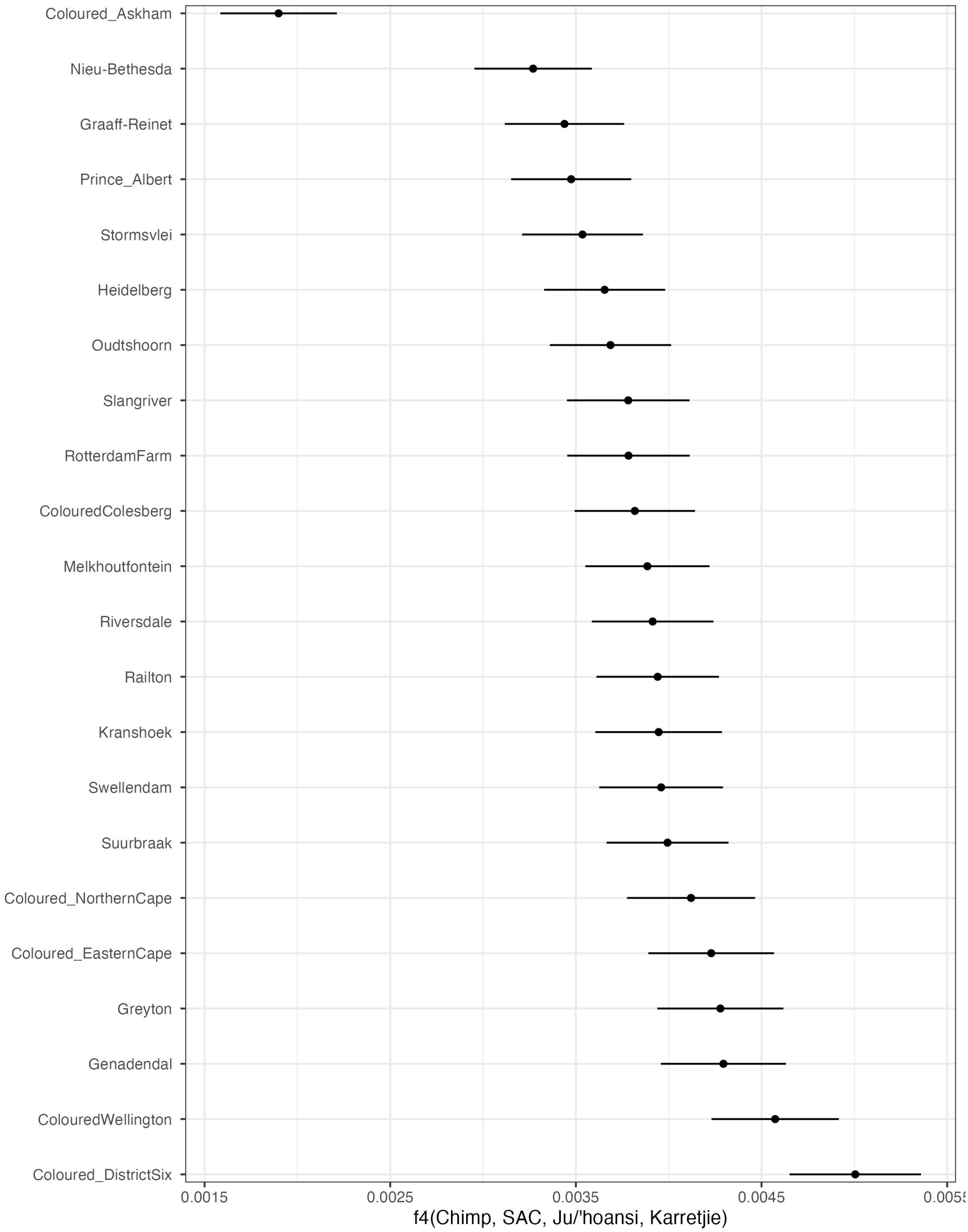


Supplementary Figure 7: Values of admixture f4-statistic in the form *f4* (Chimp, SAC, Ju/'hoansi, Karretjie).

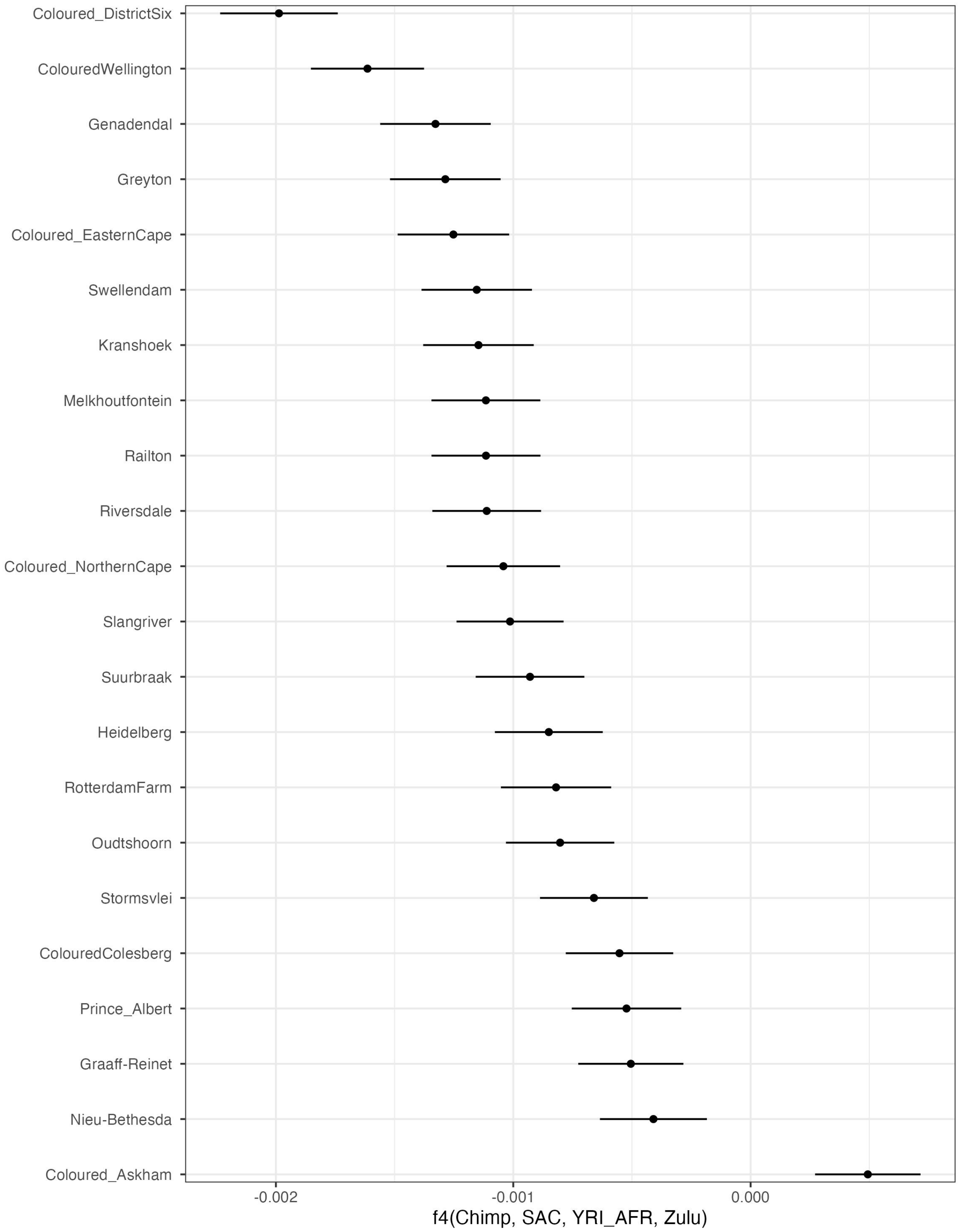


Supplementary Figure 8: Values of admixture f4-statistic in the form *f4*(Chimp, SAC, YRI_AFR, Zulu).

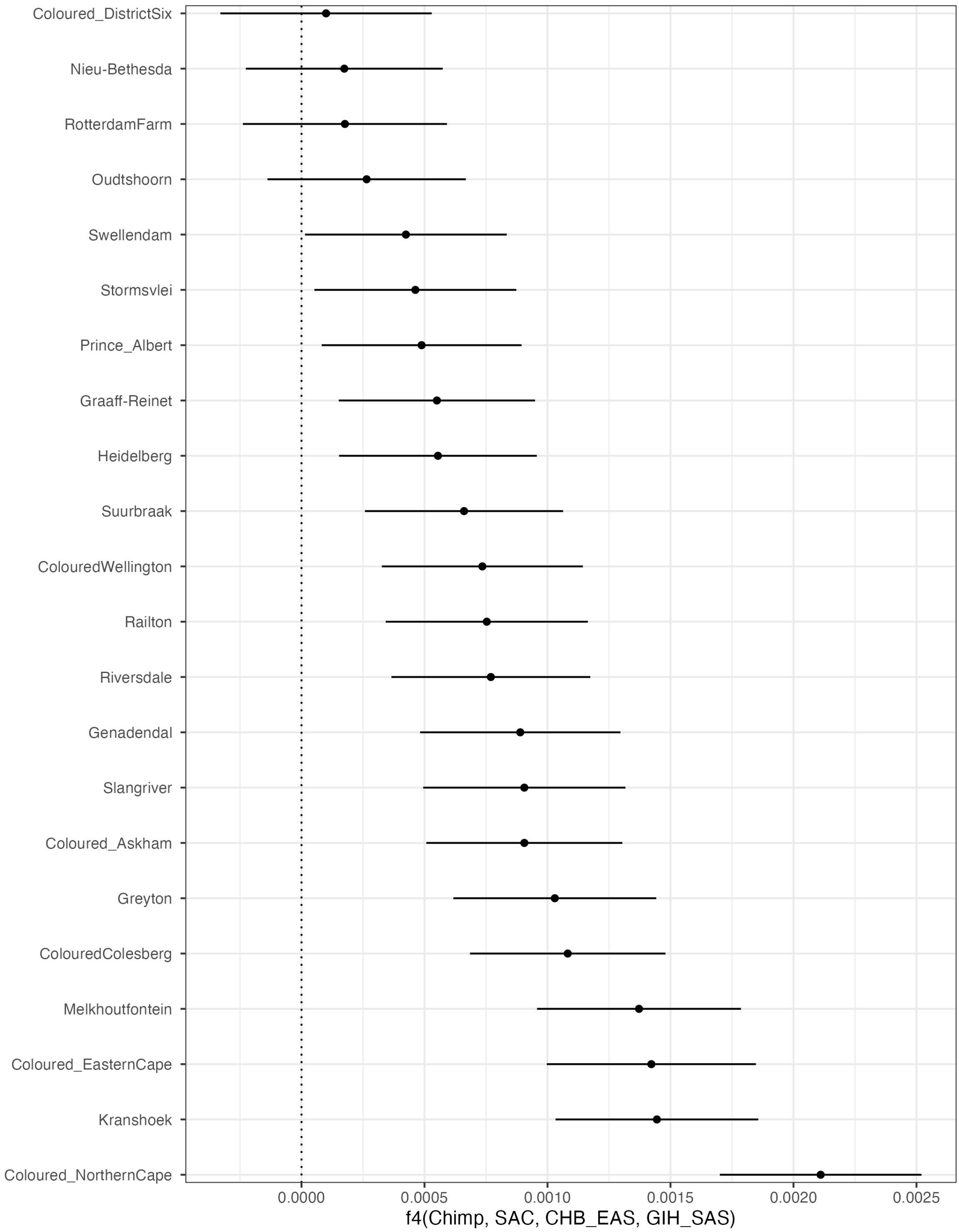


Supplementary Figure 9: Values of admixture f4-statistic in the form *f4* (Chimp, SAC, CHB_EAS, GIH_SAS).

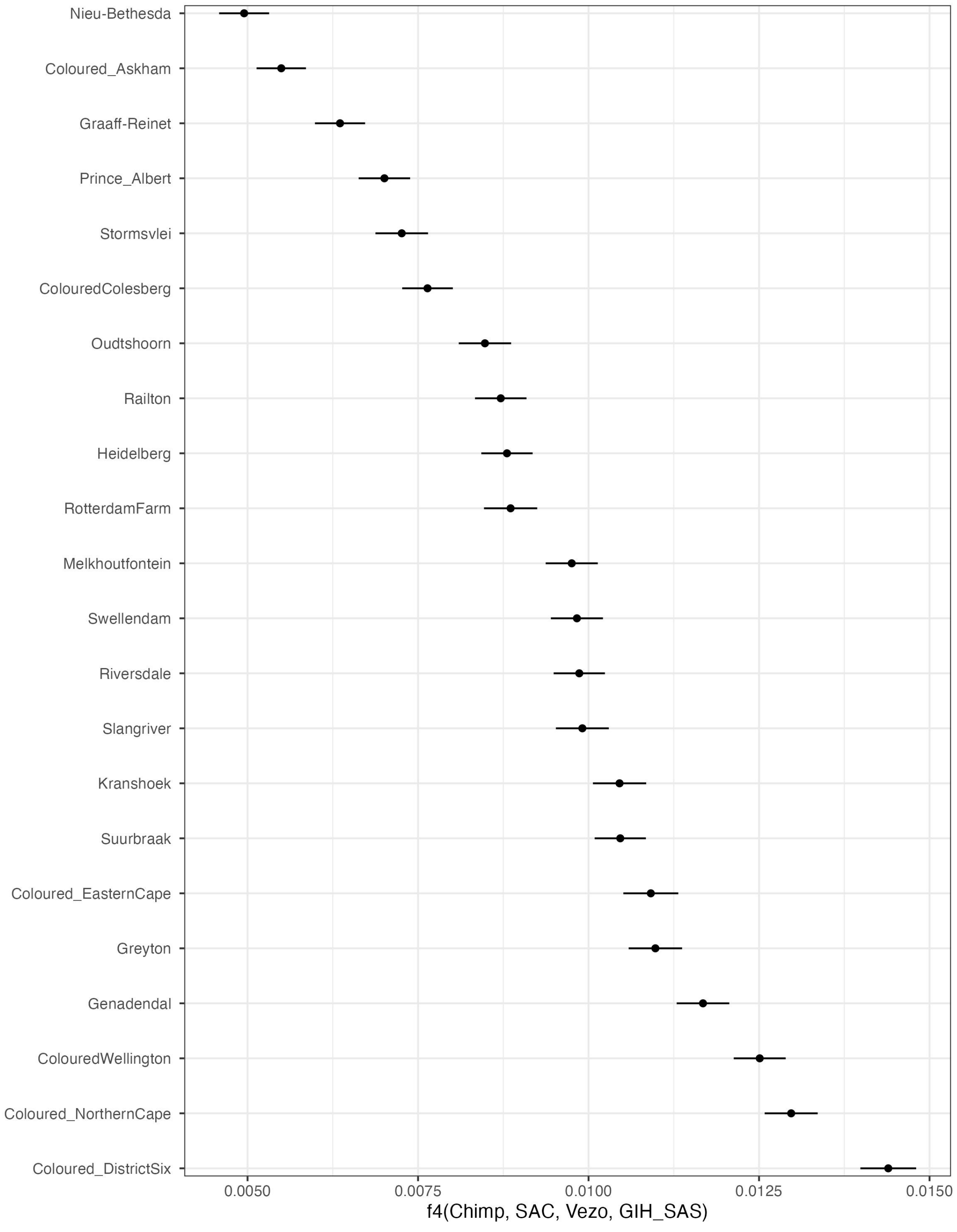


Supplementary Figure 10: Values of admixture f4-statistic in the form *f4*(Chimp, SAC, Vezo, GIH_SAS).

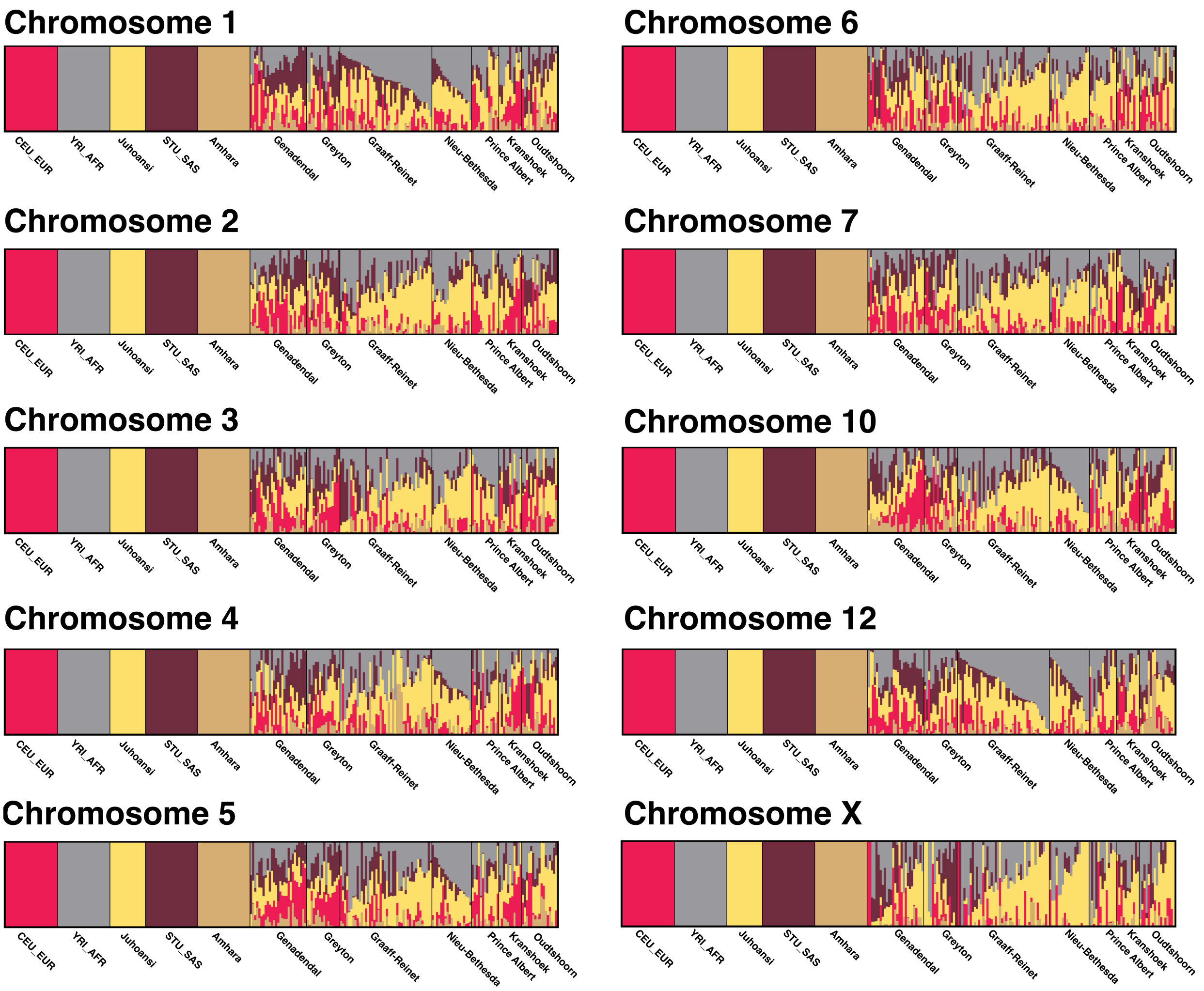


Supplementary Figure 11: Supervised ADMIXTURE results per chromosome, visualized using PONG for K = 5. These ADMIXTURE results were used to calculate the ΔAdmix ratios depicted in Figure 4. Each ADMIXTURE run is based on 13,000 SNPs.

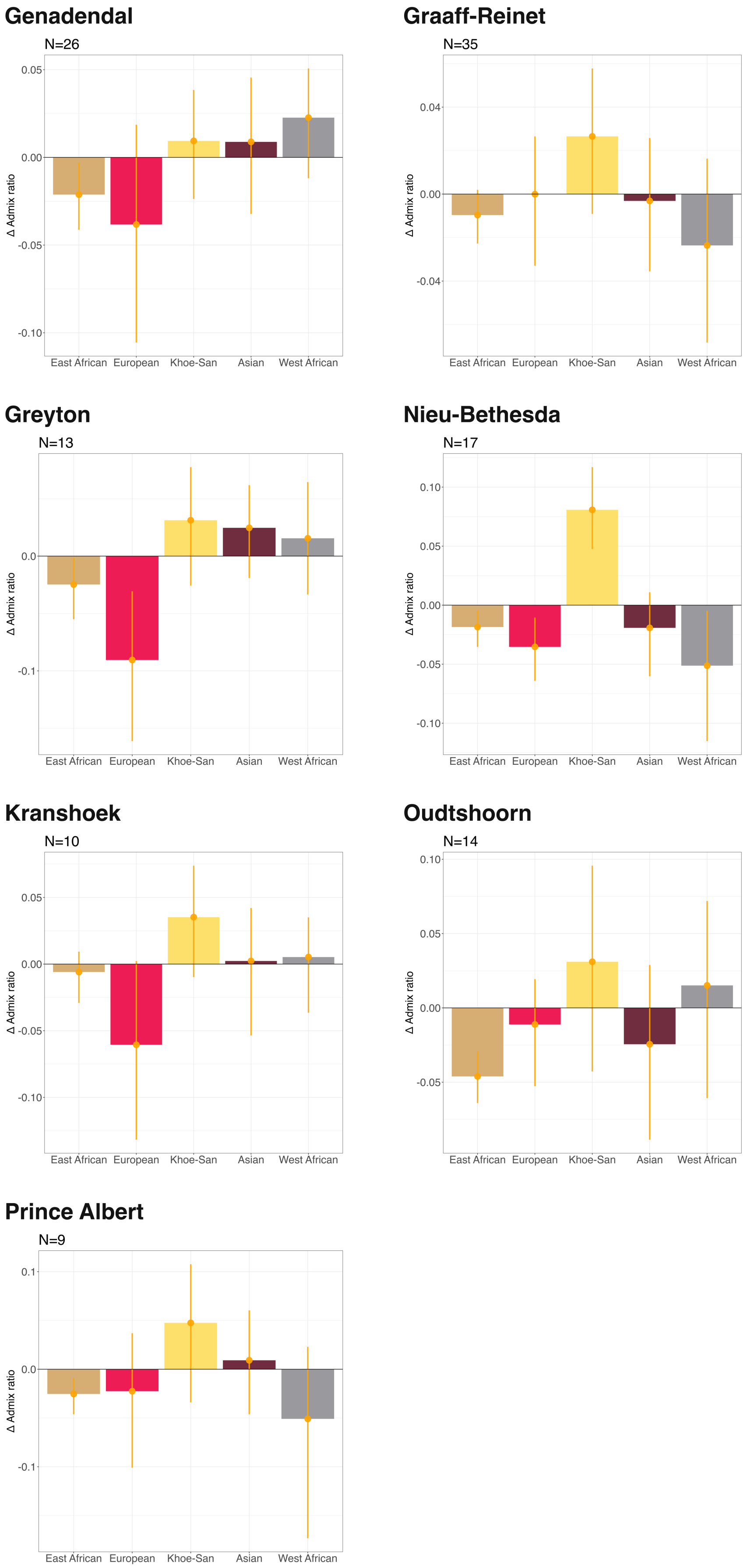


Supplementary Figure 12: ΔAdmix ratios for each of the seven ancestries, shown separately for the seven investigated sites. X and autosomal proportions were bootstrapped (10 000 times) and average X-to-autosomal difference ratios were calculated for each of the five ancestries, as well as standard deviations. The error bars indicate the 95% confidence interval. Negative X-to-autosomal difference ratios are indicative of male-biased admixture for that ancestry, positive X-to-autosomal difference ratios are indicative of a female-biased admixture for that ancestry.

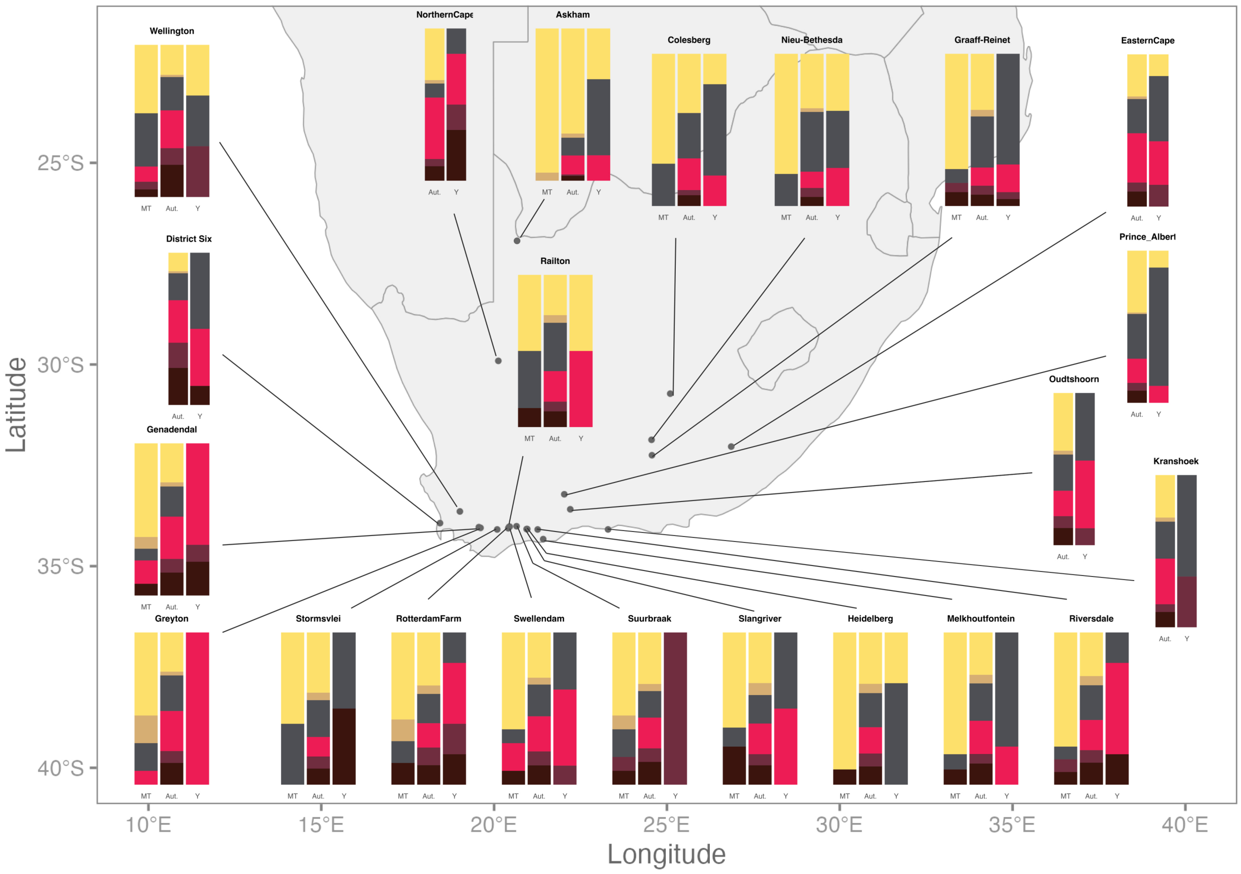


Supplementary Figure 13: Maternal, autosomal, and paternal ancestries at each location with SAC individuals. Where no information about maternal or paternal ancestries was available, these are not shown.

#### MOSAIC results

The following section (supplementary Figures 14 to 57) contains the 1-Fst values followed by the co-ancestry curves for each of the SAC populations in the higher density dataset. The ancestries from left to right correspond to the ancestry numbers (1-5) in the co-ancestry curves. Each ancestry is plotted against itself and each of the other ancestries.

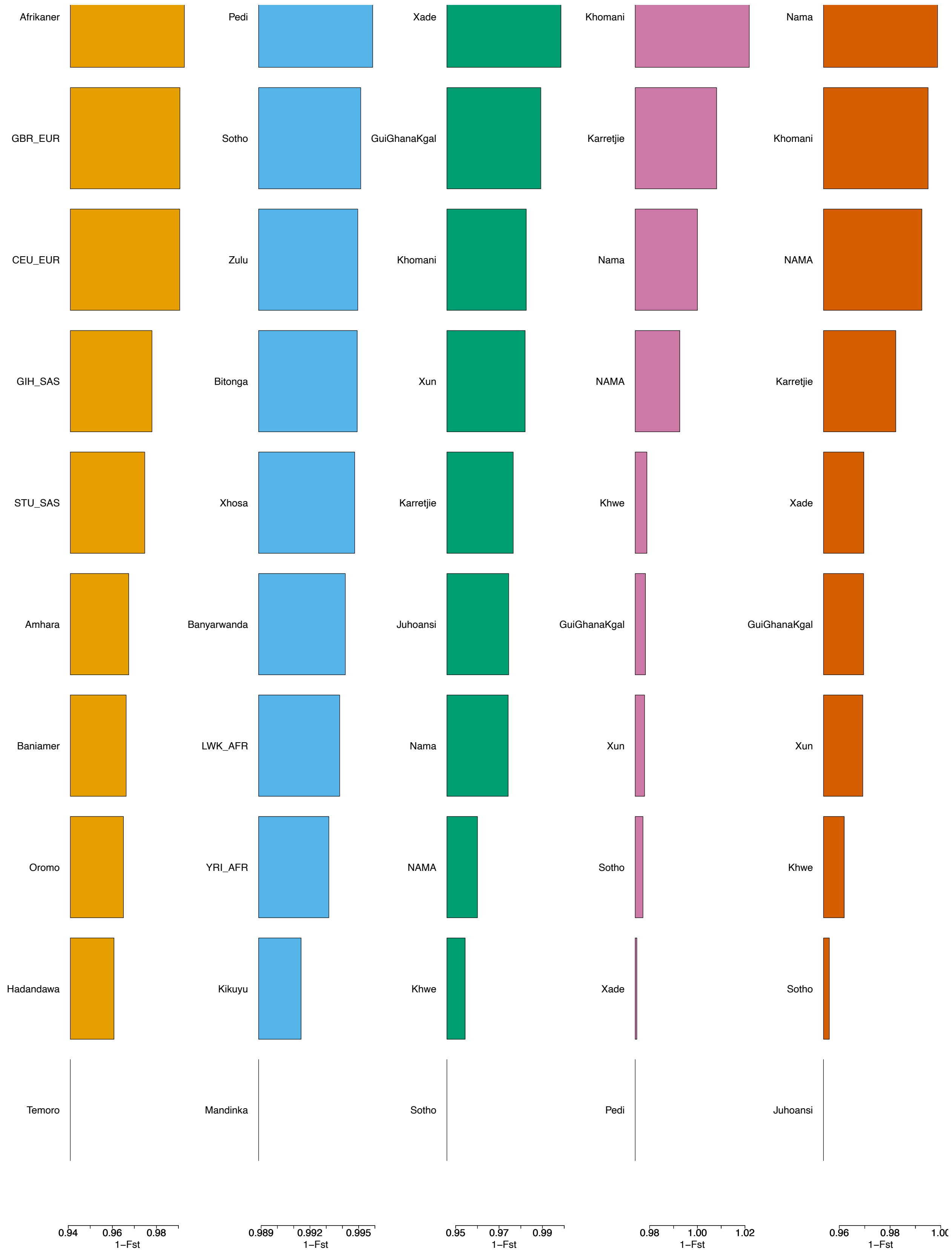


Supplementary Figure 14: 1-$F_{st}$ values from MOSAIC for the 5 constructed ancestries for Askham.

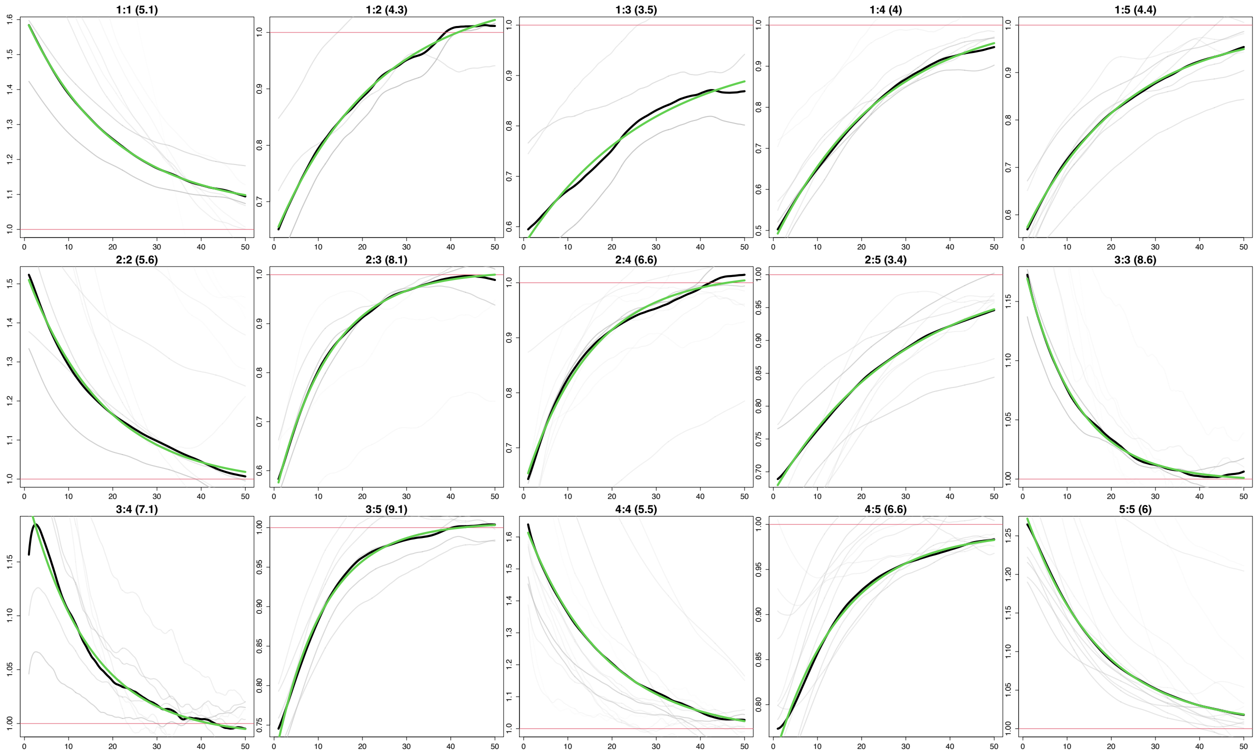


Supplementary Figure 15: Co-Ancestry curves from MOSAIC for the 5 constructed ancestries for Askham. The y-axis depicts the relative probability of jointly copying two chunks from the donor populations and x-axis is the genetic distance in centimorgans. The black lines represent the empirical coancestry curves across all target individuals, the light grey are per individual, and the green lines represent the fitted single-event coancestry curve. Above each plot, the number of the populations being examined is noted as a:b, with the number of generations since the admixture event in parentheses behind it. The number represents the order the ancestries appear in in the 1-$F_{st}$ plot.

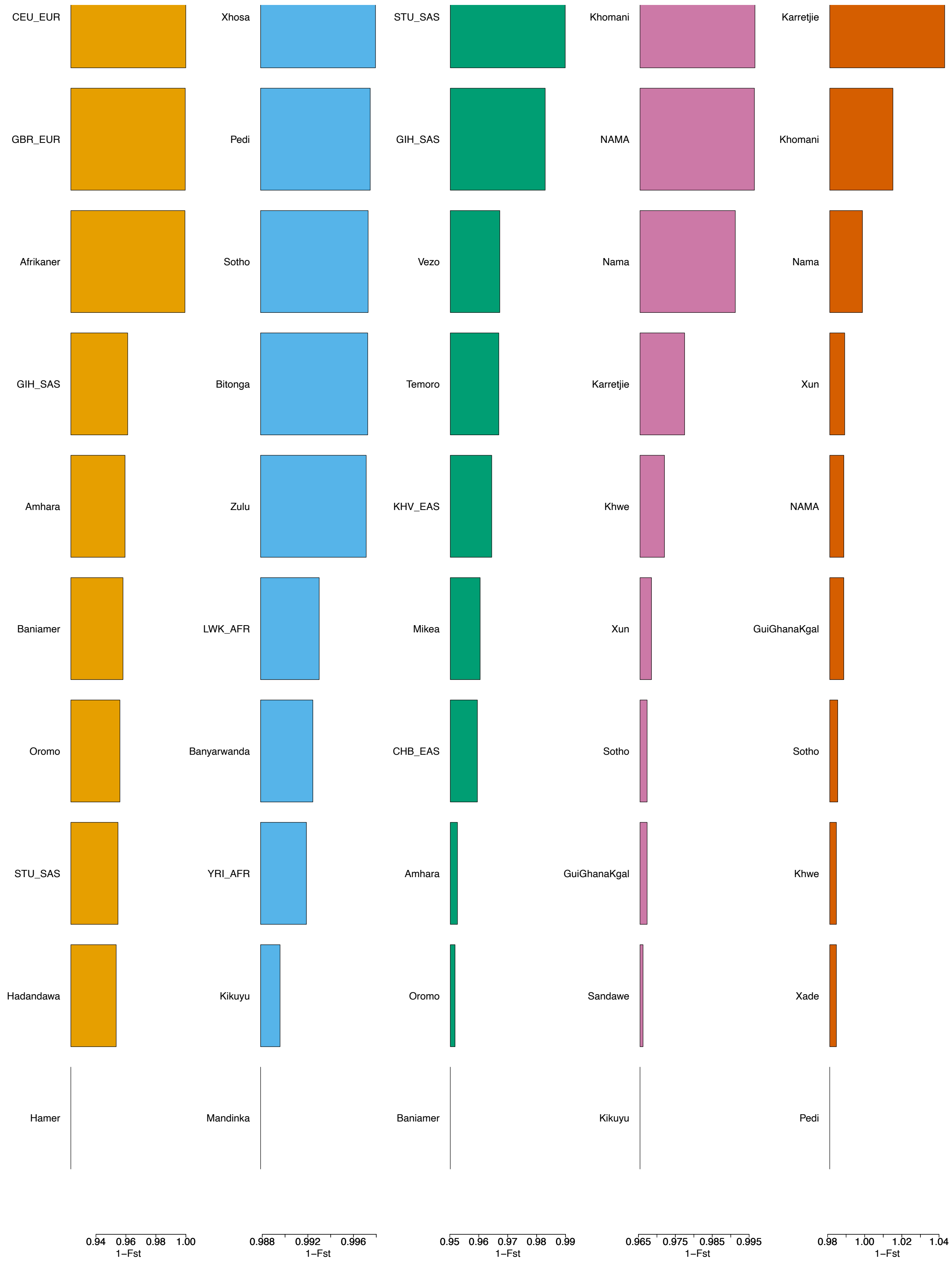


Supplementary Figure 16: 1-$F_{st}$ values from MOSAIC for the 5 constructed ancestries for ColouredColesberg.

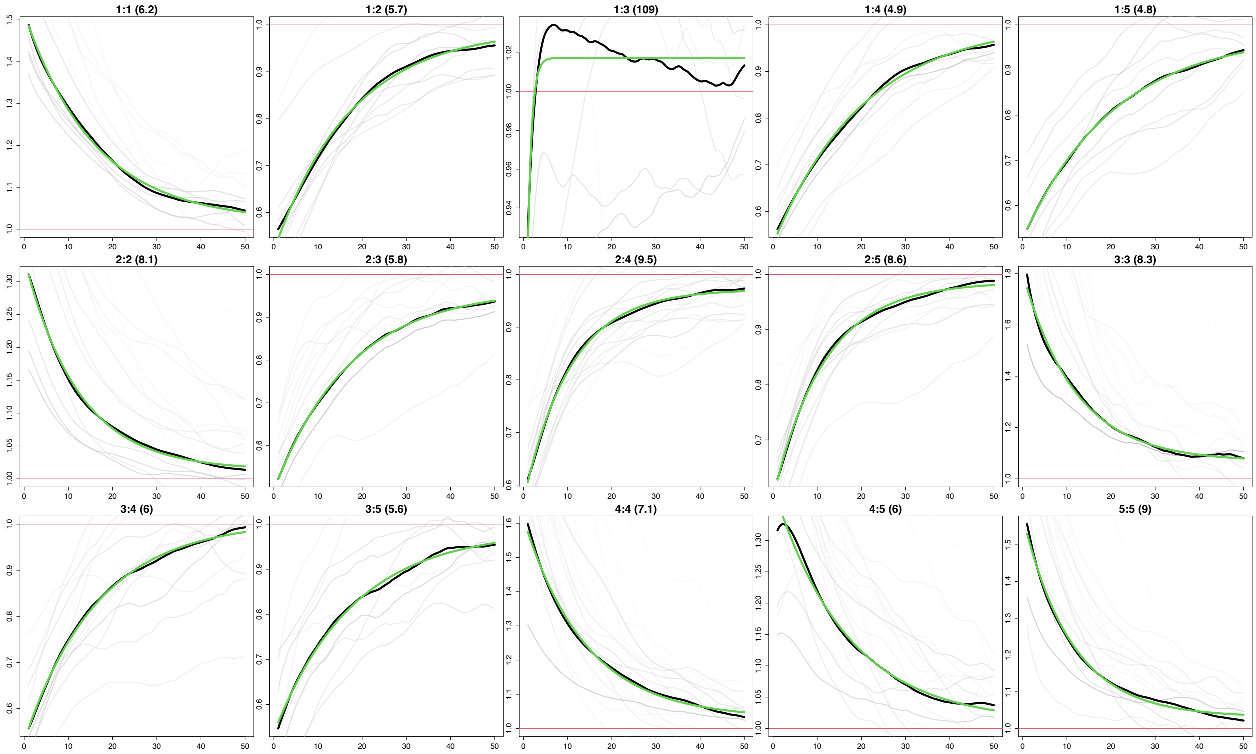


Supplementary Figure 17: Co-Ancestry curves from MOSAIC for the 5 constructed ancestries for Coloured-Colesberg.The y-axis depicts the relative probability of jointly copying two chunks from the donor populations and x-axis is the genetic distance in centimorgans. The black lines represent the empirical coancestry curves across all target individuals, the light grey are per individual, and the green lines represent the fitted single-event coancestry curve. Above each plot, the number of the populations being examined is noted as a:b, with the number of generations since the admixture event in parentheses behind it. The number represents the order the ancestries appear in in the 1-$F_{st}$ plot.

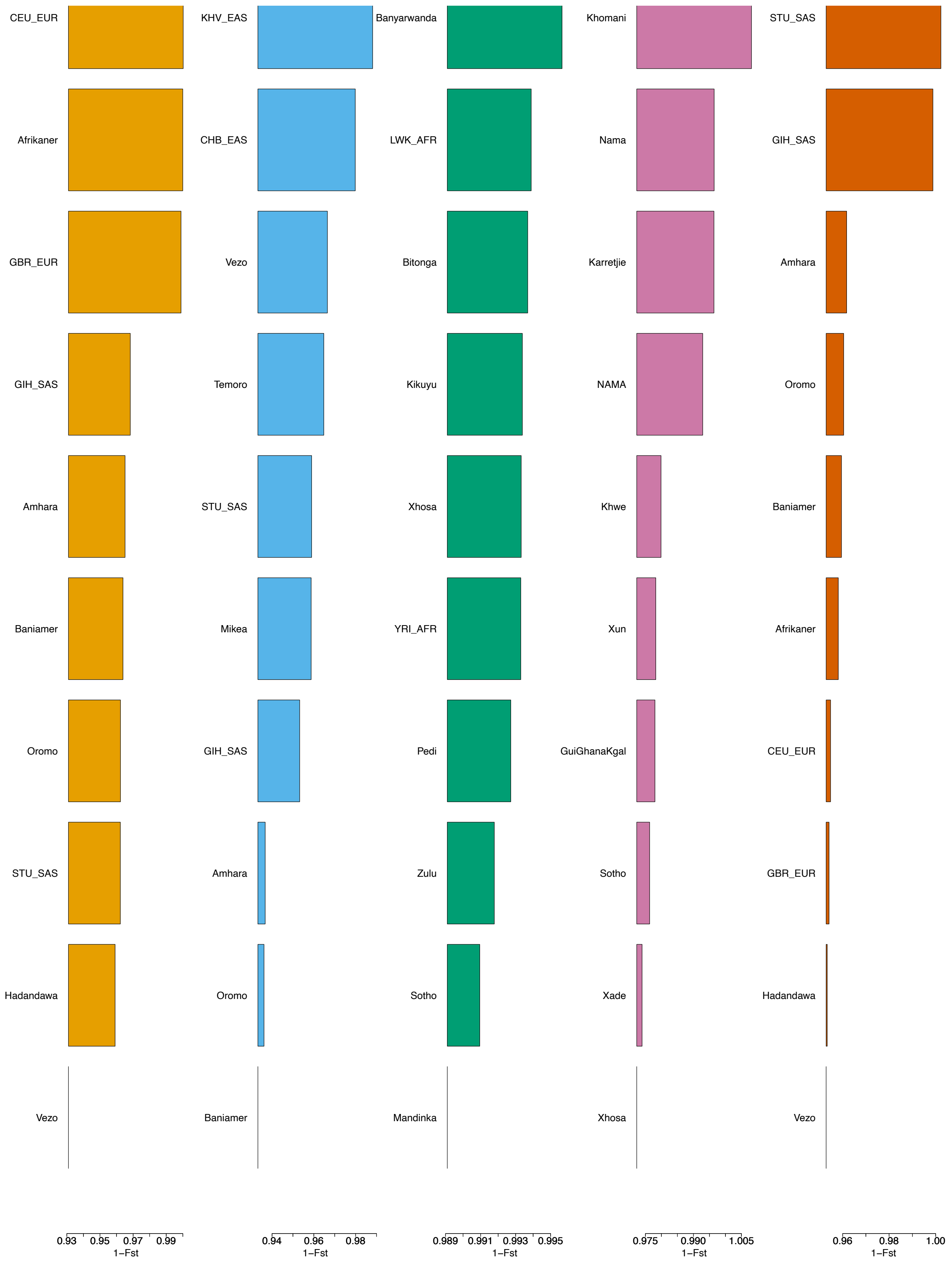


Supplementary Figure 18: 1-$F_{st}$ values from MOSAIC for the 5 constructed ancestries for Coloured District Six.

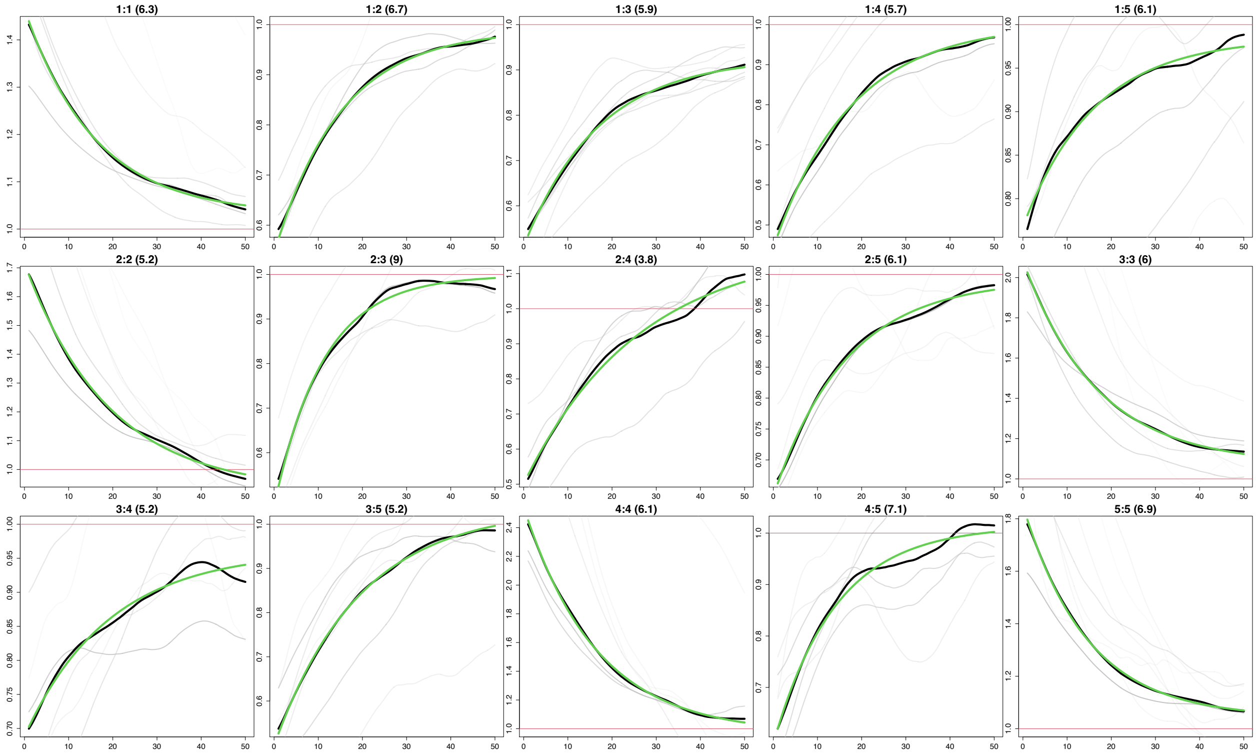


Supplementary Figure 19: Co-Ancestry curves from MOSAIC for the 5 constructed ancestries for Coloured District Six.The y-axis depicts the relative probability of jointly copying two chunks from the donor populations and x-axis is the genetic distance in centimorgans. The black lines represent the empirical coancestry curves across all target individuals, the light grey are per individual, and the green lines represent the fitted single-event coancestry curve. Above each plot, the number of the populations being examined is noted as a:b, with the number of generations since the admixture event in parentheses behind it. The number represents the order the ancestries appear in in the 1-$F_{st}$ plot.

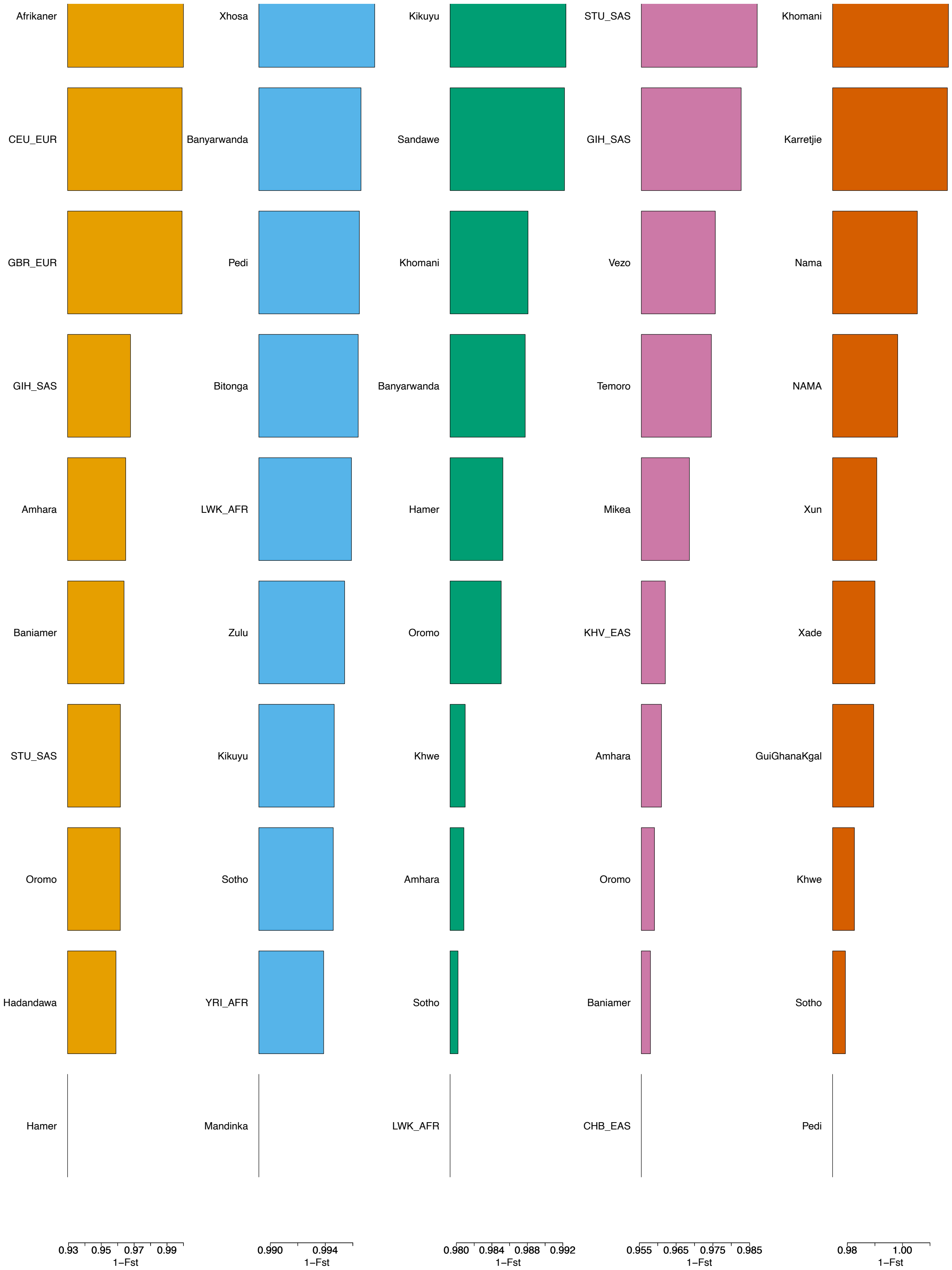


Supplementary Figure 20: 1-$F_{st}$ values from MOSAIC for the 5 constructed ancestries for Coloured Eastern Cape.

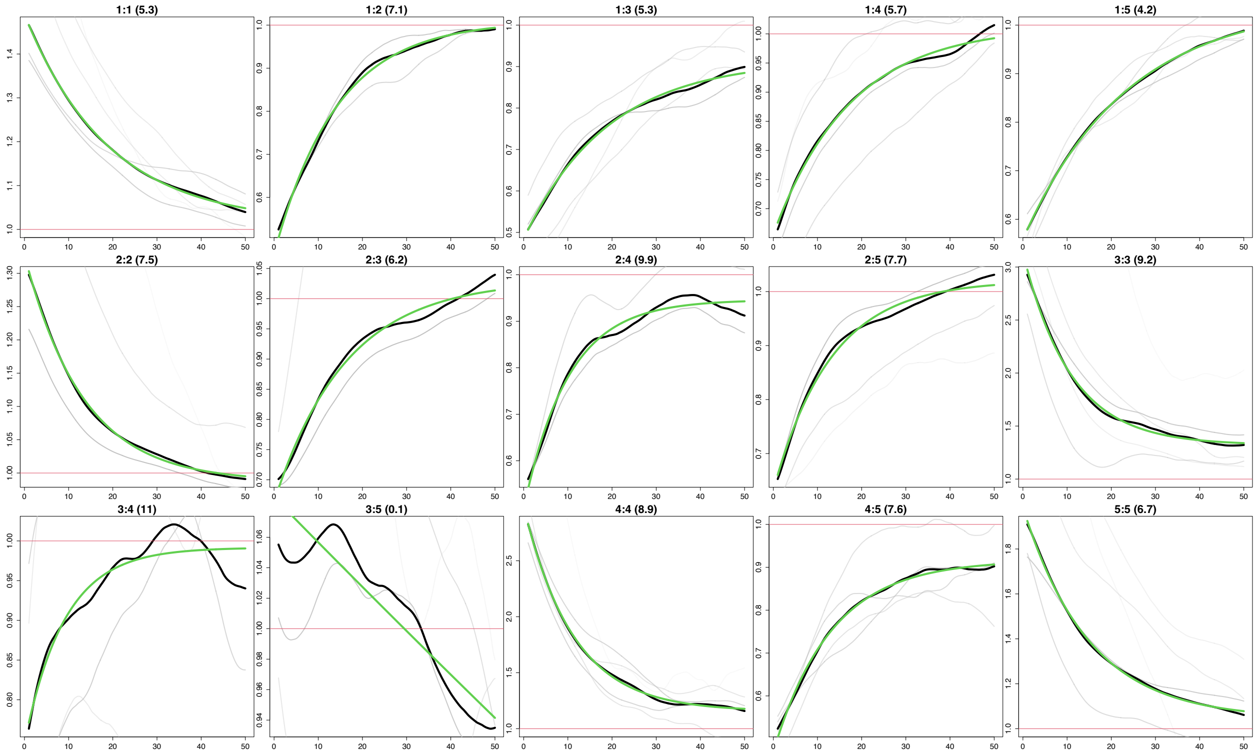


Supplementary Figure 21: Co-Ancestry curves from MOSAIC for the 5 constructed ancestries for Coloured Eastern Cape.The y-axis depicts the relative probability of jointly copying two chunks from the donor populations and x-axis is the genetic distance in centimorgans. The black lines represent the empirical coancestry curves across all target individuals, the light grey are per individual, and the green lines represent the fitted single-event coancestry curve. Above each plot, the number of the populations being examined is noted as a:b, with the number of generations since the admixture event in parentheses behind it. The number represents the order the ancestries appear in in the 1-$F_{st}$ plot.

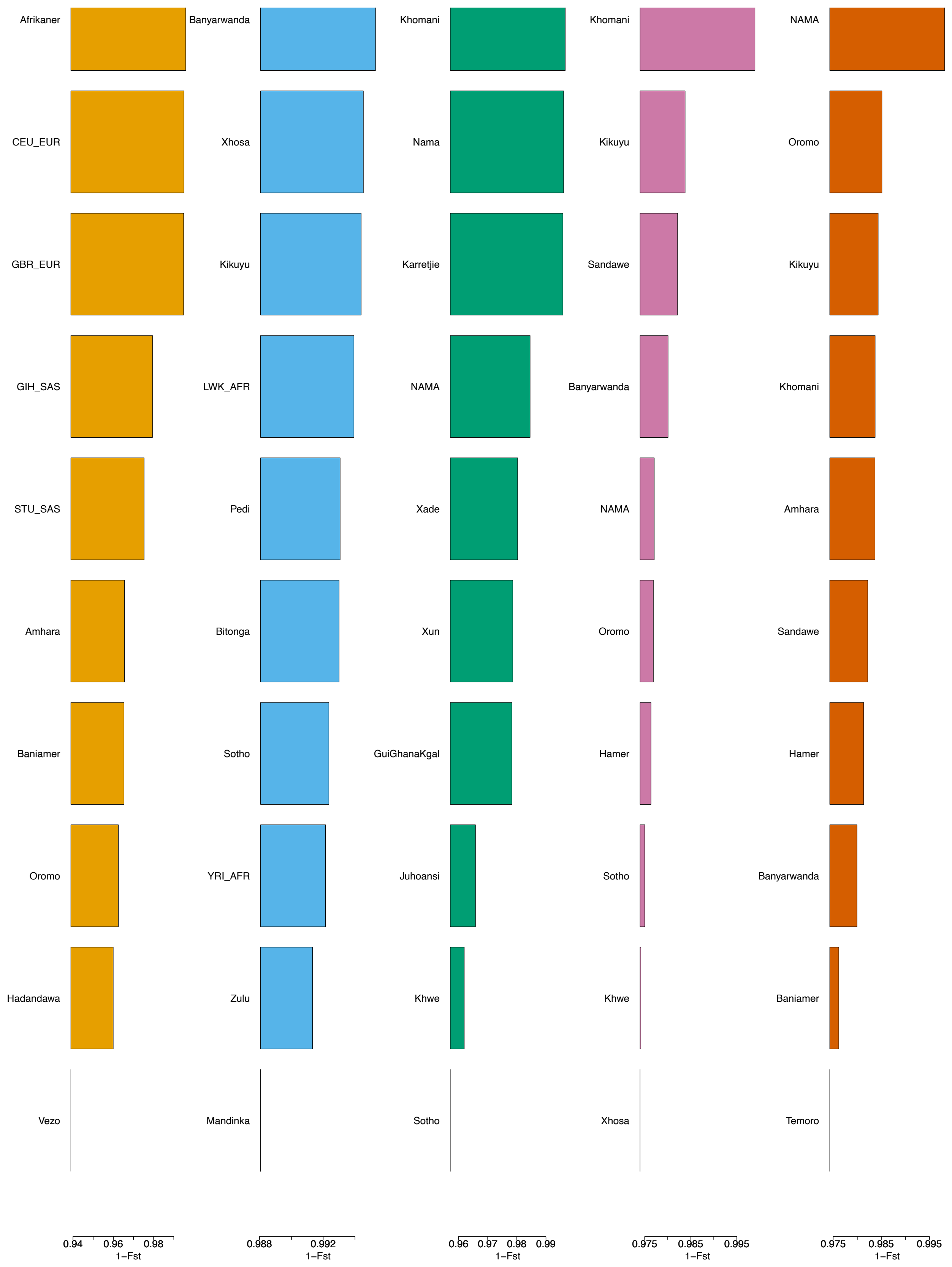


Supplementary Figure 22: 1-$F_{st}$ values from MOSAIC for the 5 constructed ancestries for Coloured Northern Cape.

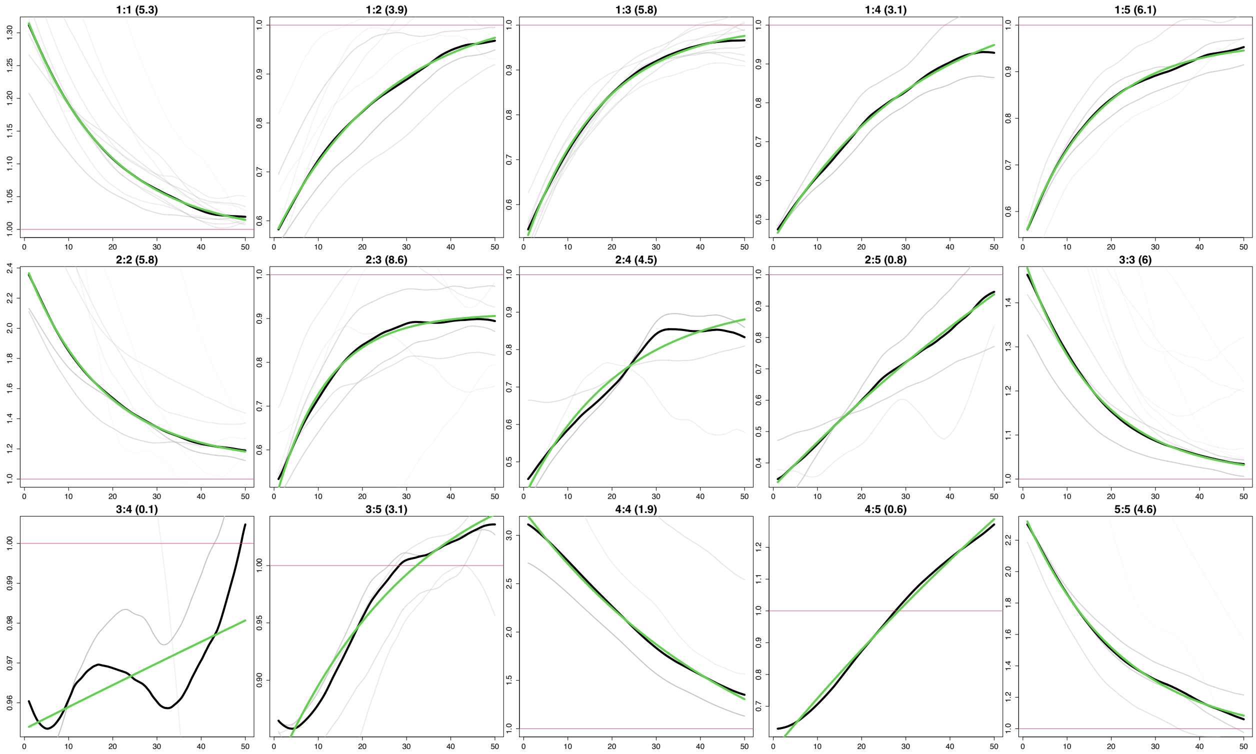


Supplementary Figure 23: Co-Ancestry curves from MOSAIC for the 5 constructed ancestries for Coloured Northern Cape. The y-axis depicts the relative probability of jointly copying two chunks from the donor populations and x-axis is the genetic distance in centimorgans. The black lines represent the empirical coancestry curves across all target individuals, the light grey are per individual, and the green lines represent the fitted single-event coancestry curve. Above each plot, the number of the populations being examined is noted as a:b, with the number of generations since the admixture event in parentheses behind it. The number represents the order the ancestries appear in in the 1-$F_{st}$ plot.

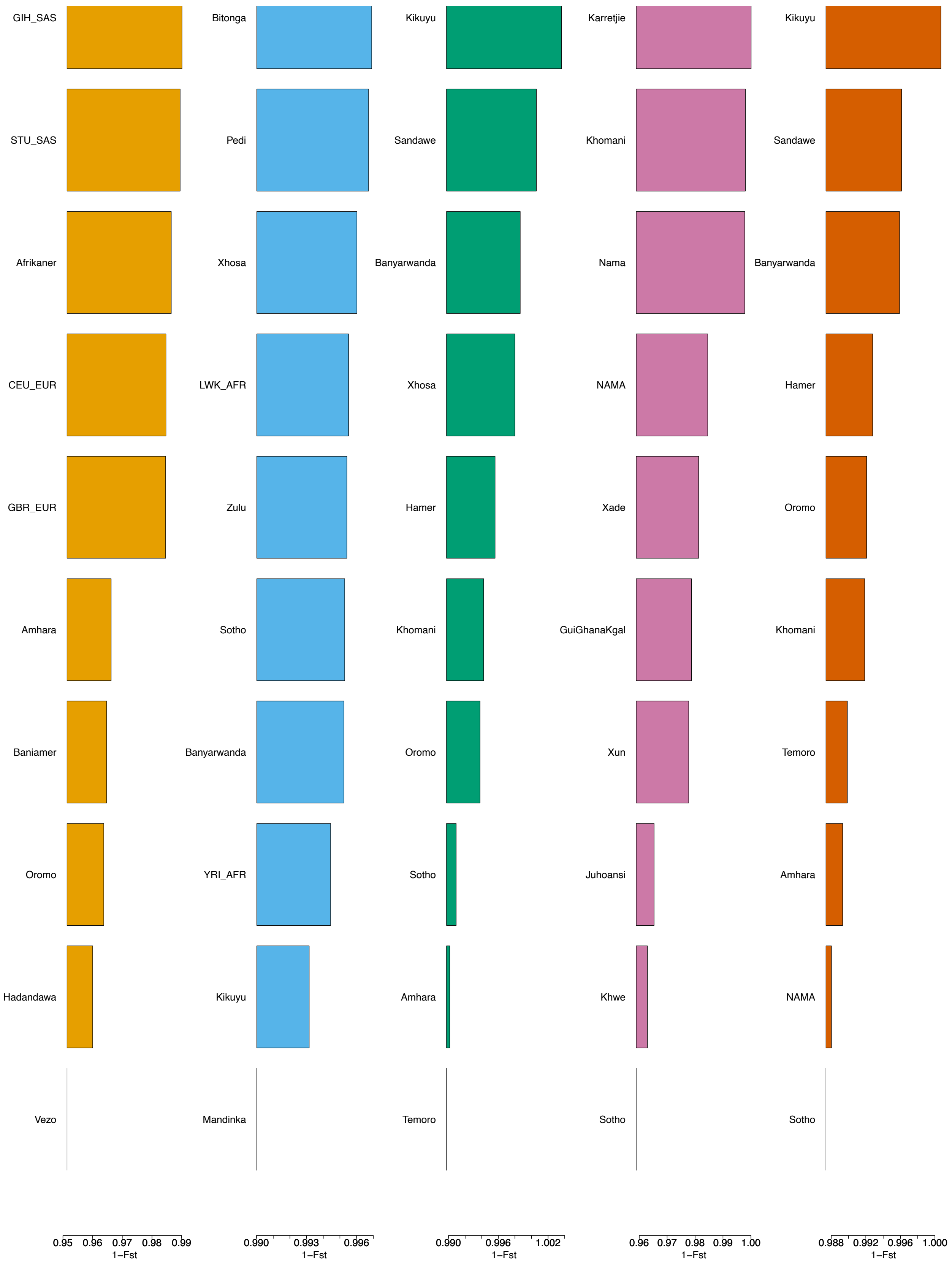


Supplementary Figure 24: 1-$F_{st}$ values from MOSAIC for the 5 constructed ancestries for ColouredWellington.

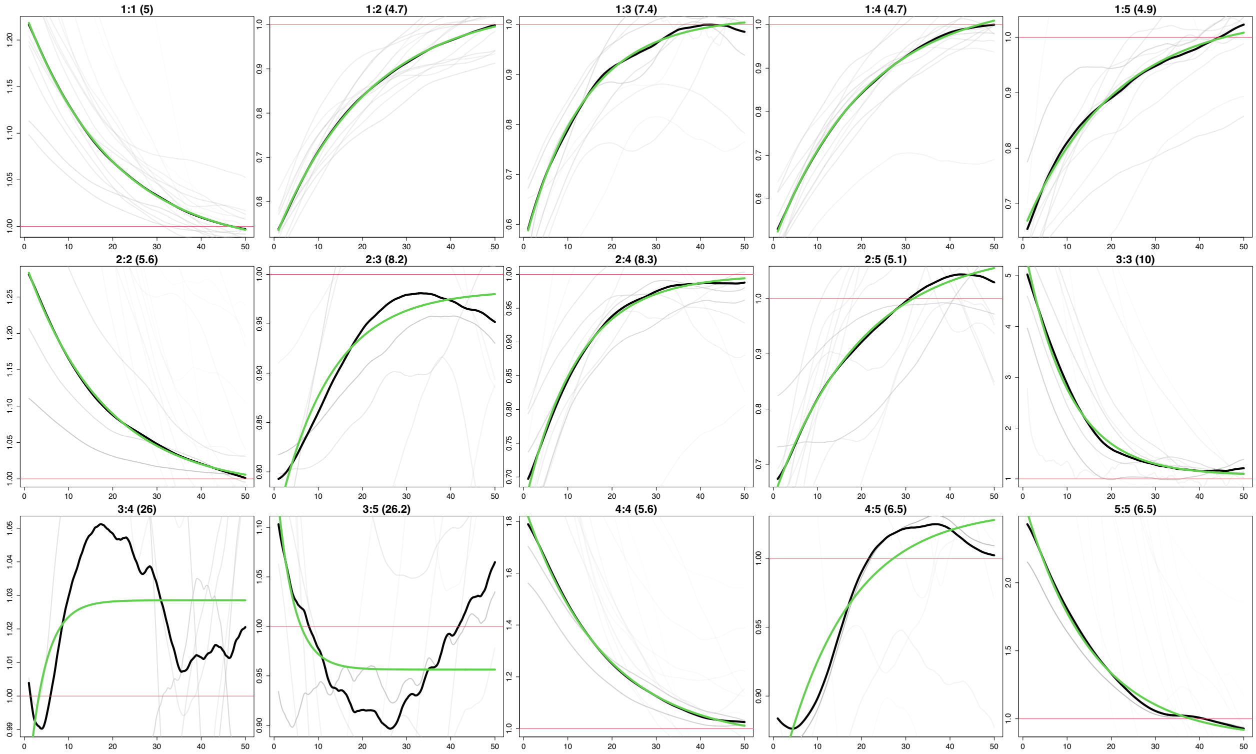


Supplementary Figure 25: Co-Ancestry curves from MOSAIC for the 5 constructed ancestries for Coloured-Wellington. The y-axis depicts the relative probability of jointly copying two chunks from the donor populations and x-axis is the genetic distance in centimorgans. The black lines represent the empirical coancestry curves across all target individuals, the light grey are per individual, and the green lines represent the fitted single-event coancestry curve. Above each plot, the number of the populations being examined is noted as a:b, with the number of generations since the admixture event in parentheses behind it. The number represents the order the ancestries appear in in the 1-$F_{st}$ plot.

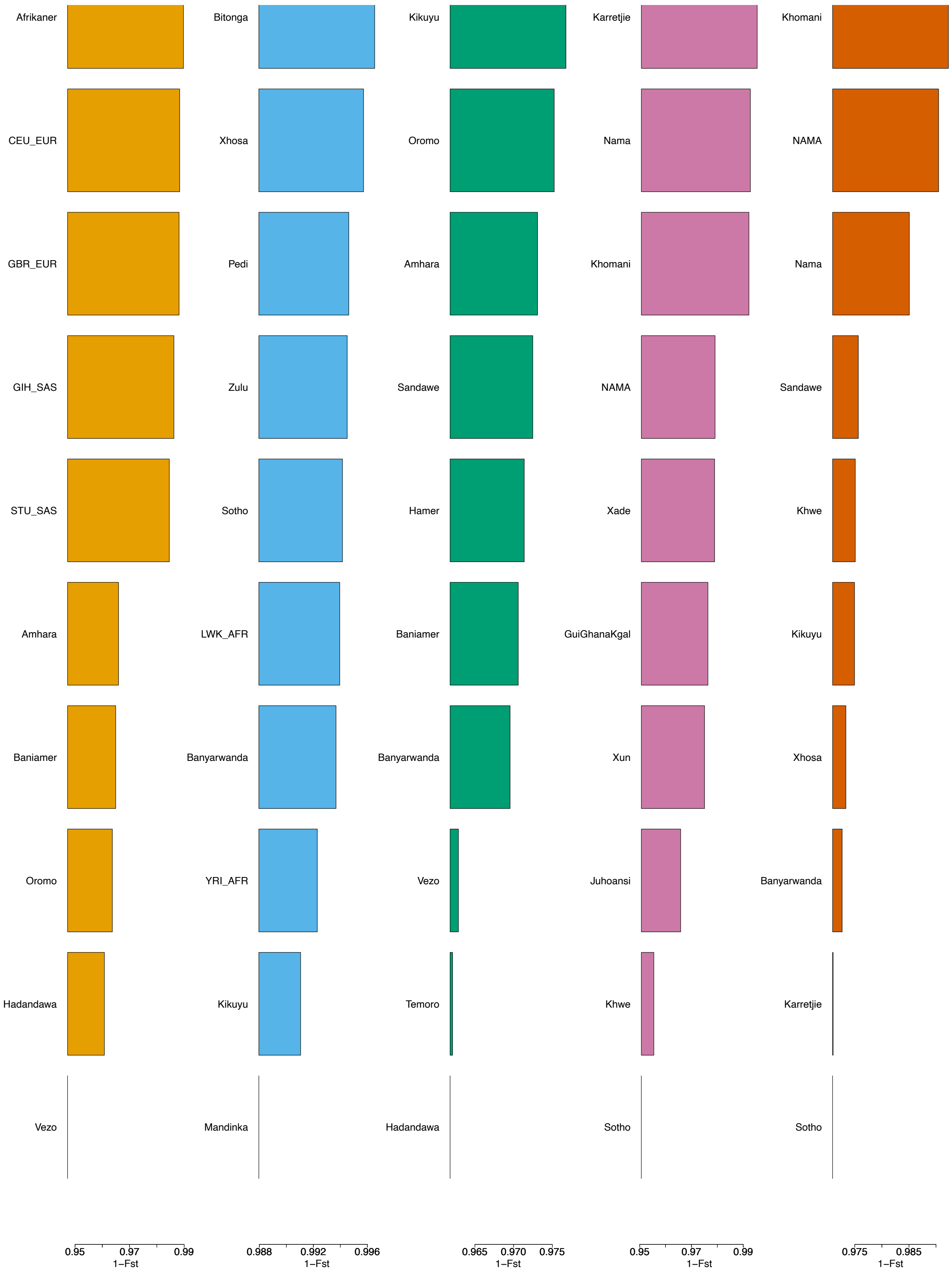


Supplementary Figure 26: 1-$F_{st}$ values from MOSAIC for the 5 constructed ancestries for Genadendal.

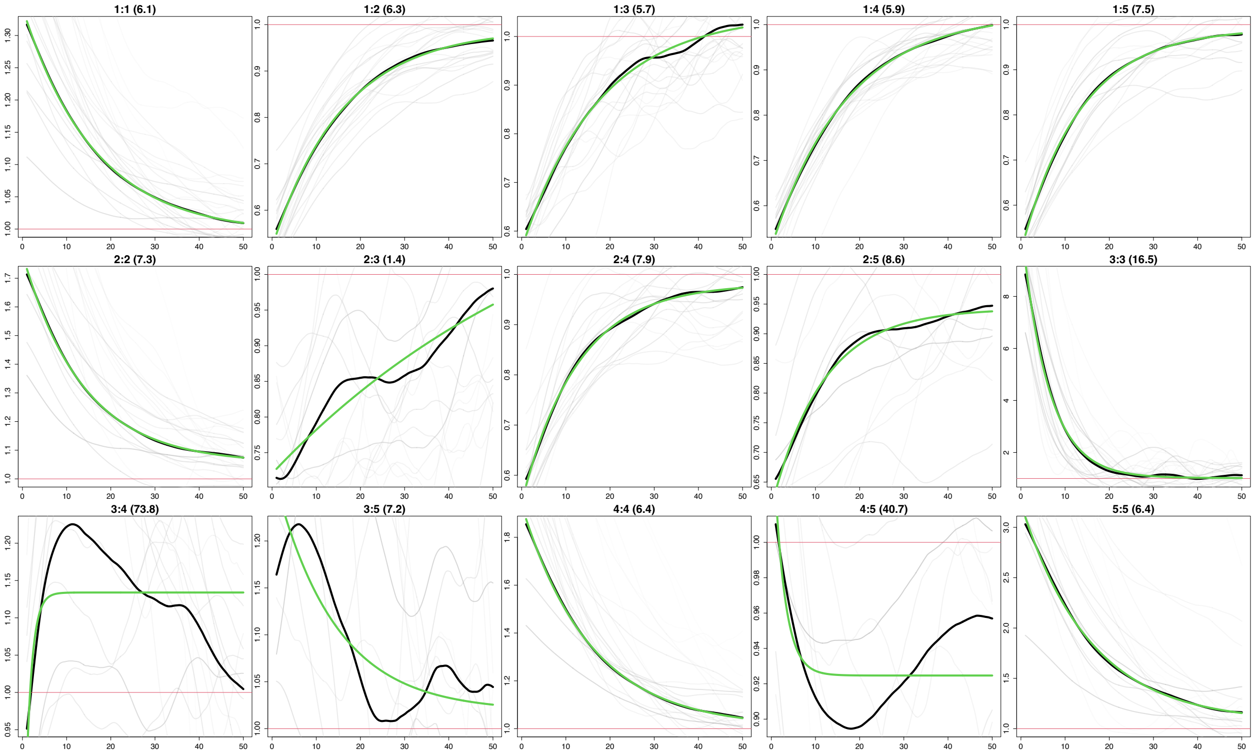


Supplementary Figure 27: Co-Ancestry curves from MOSAIC for the 5 constructed ancestries for Genadendal. The y-axis depicts the relative probability of jointly copying two chunks from the donor populations and x-axis is the genetic distance in centimorgans. The black lines represent the empirical coancestry curves across all target individuals, the light grey are per individual, and the green lines represent the fitted single-event coancestry curve. Above each plot, the number of the populations being examined is noted as a:b, with the number of generations since the admixture event in parentheses behind it. The number represents the order the ancestries appear in in the 1-$F_{st}$ plot.

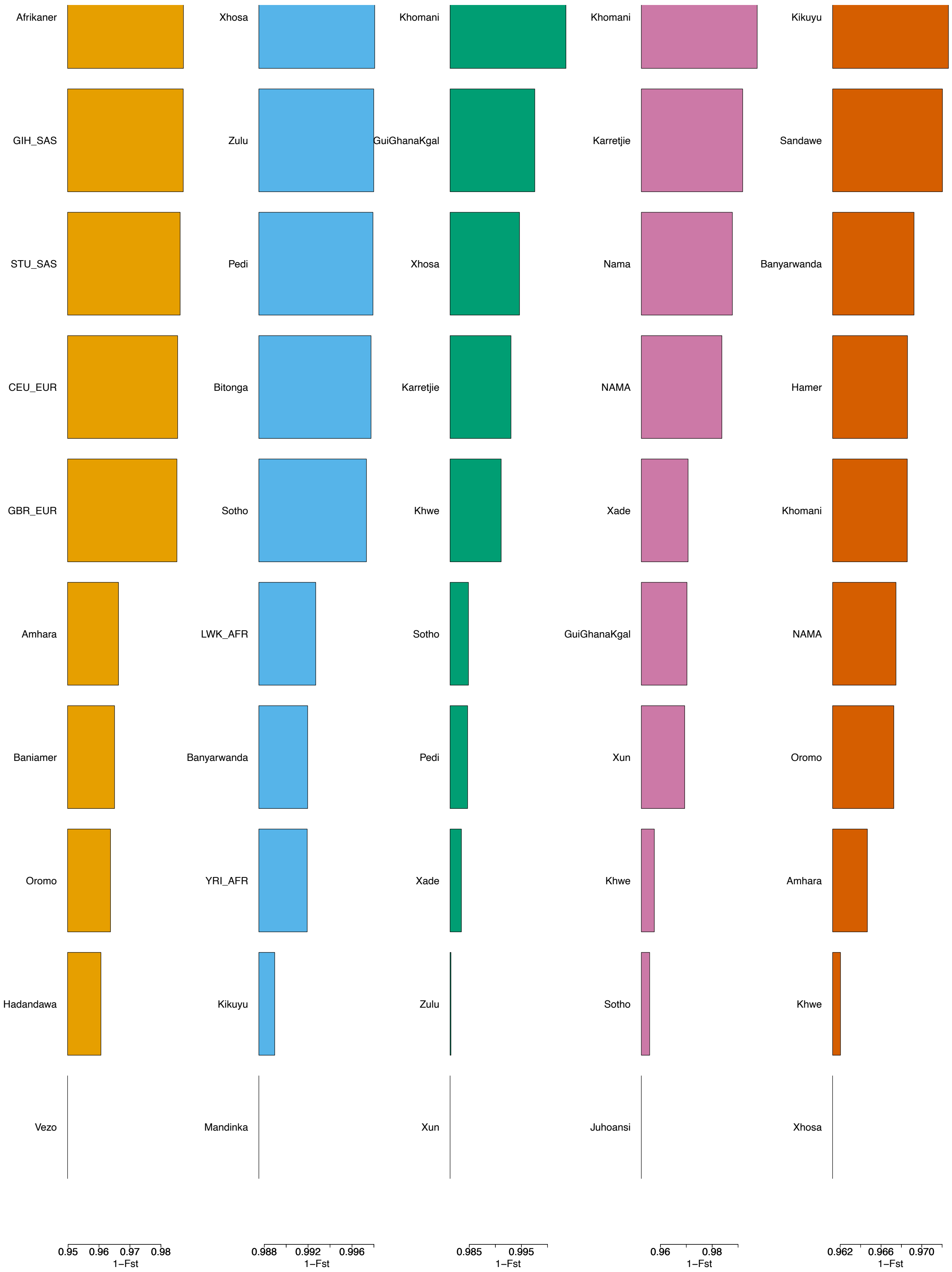


Supplementary Figure 28: 1-$F_{st}$ values from MOSAIC for the 5 constructed ancestries for Graaff-Reinet.

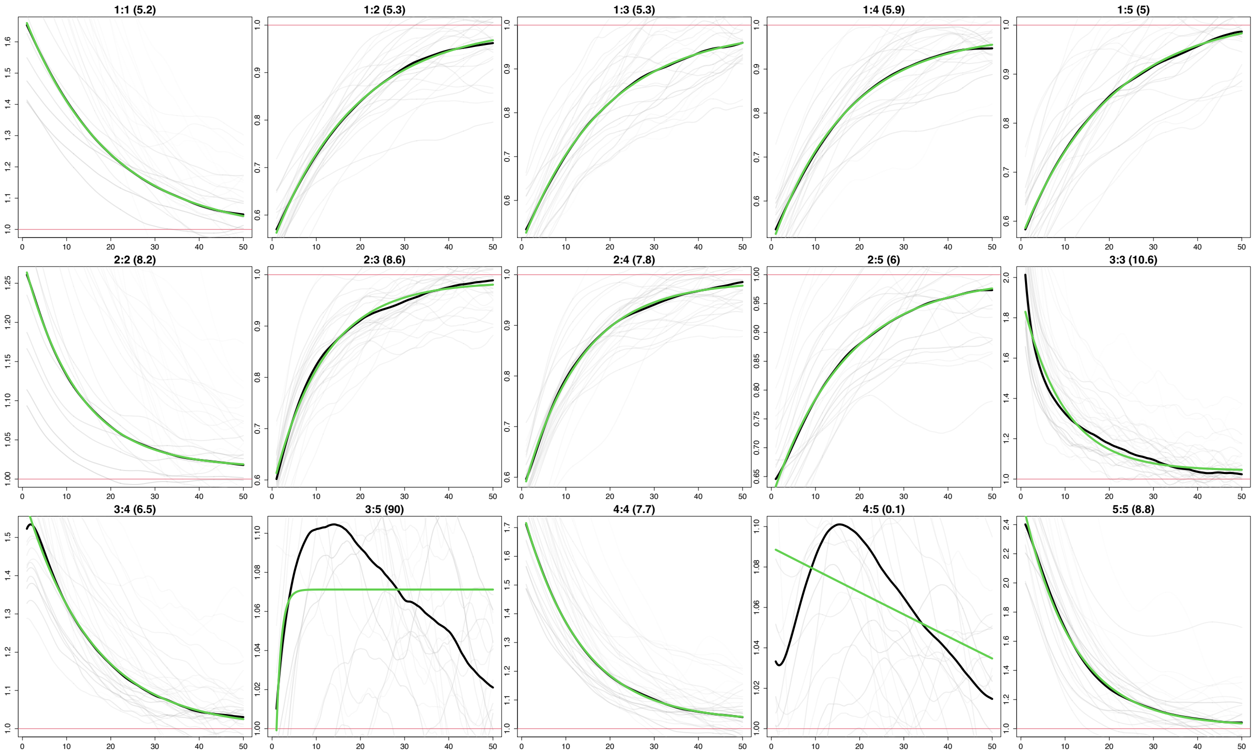


Supplementary Figure 29: Co-Ancestry curves from MOSAIC for the 5 constructed ancestries for Graaff-Reinet. The y-axis depicts the relative probability of jointly copying two chunks from the donor populations and x-axis is the genetic distance in centimorgans. The black lines represent the empirical coancestry curves across all target individuals, the light grey are per individual, and the green lines represent the fitted single-event coancestry curve. Above each plot, the number of the populations being examined is noted as a:b, with the number of generations since the admixture event in parentheses behind it. The number represents the order the ancestries appear in in the 1-$F_{st}$ plot.

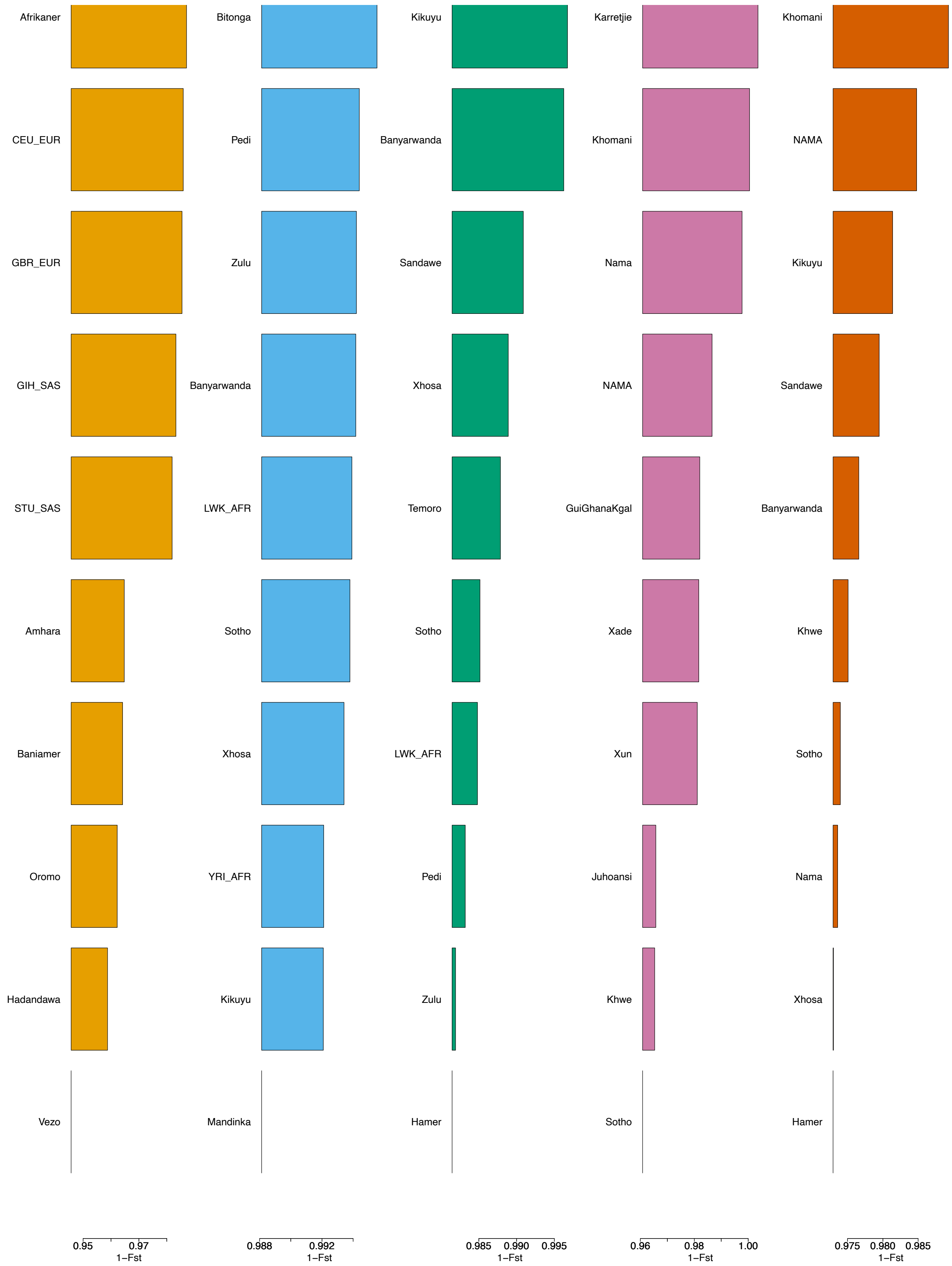


Supplementary Figure 30: 1-$F_{st}$ values from MOSAIC for the 5 constructed ancestries for Greyton.

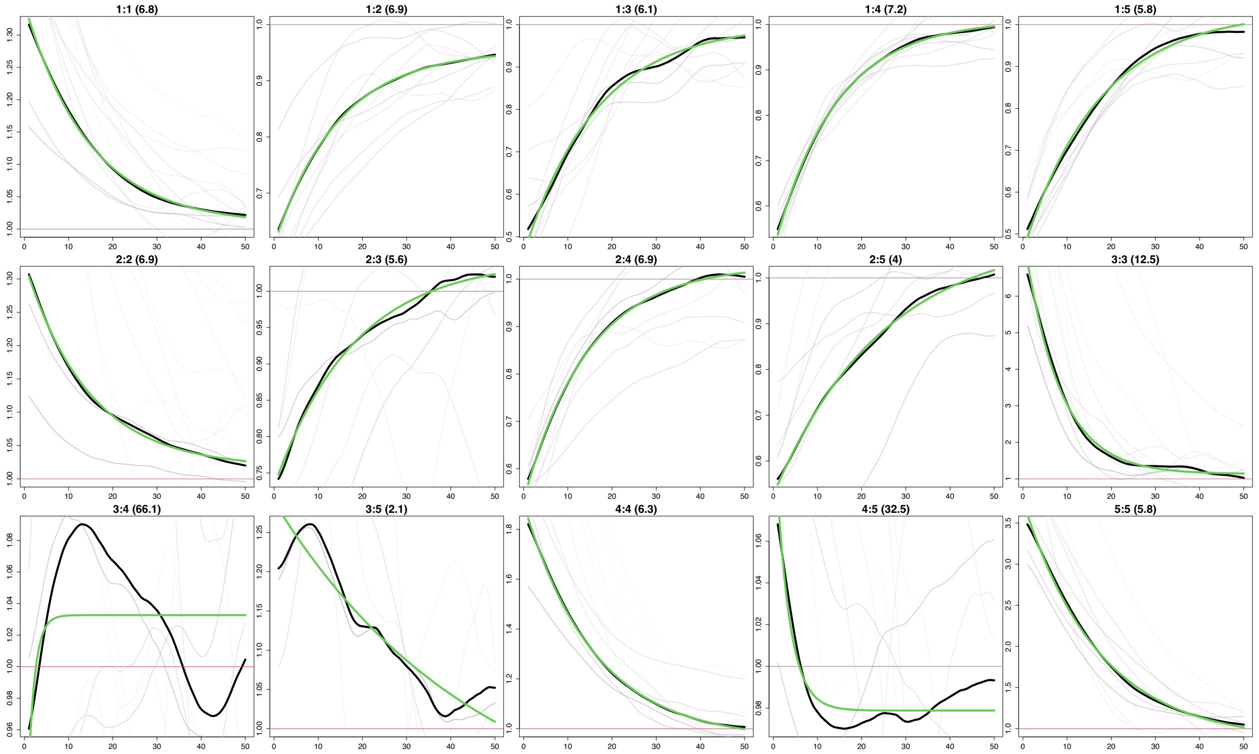


Supplementary Figure 31: Co-Ancestry curves from MOSAIC for the 5 constructed ancestries for Greyton. The y-axis depicts the relative probability of jointly copying two chunks from the donor populations and x-axis is the genetic distance in centimorgans. The black lines represent the empirical coancestry curves across all target individuals, the light grey are per individual, and the green lines represent the fitted single-event coancestry curve. Above each plot, the number of the populations being examined is noted as a:b, with the number of generations since the admixture event in parentheses behind it. The number represents the order the ancestries appear in in the 1-$F_{st}$ plot.

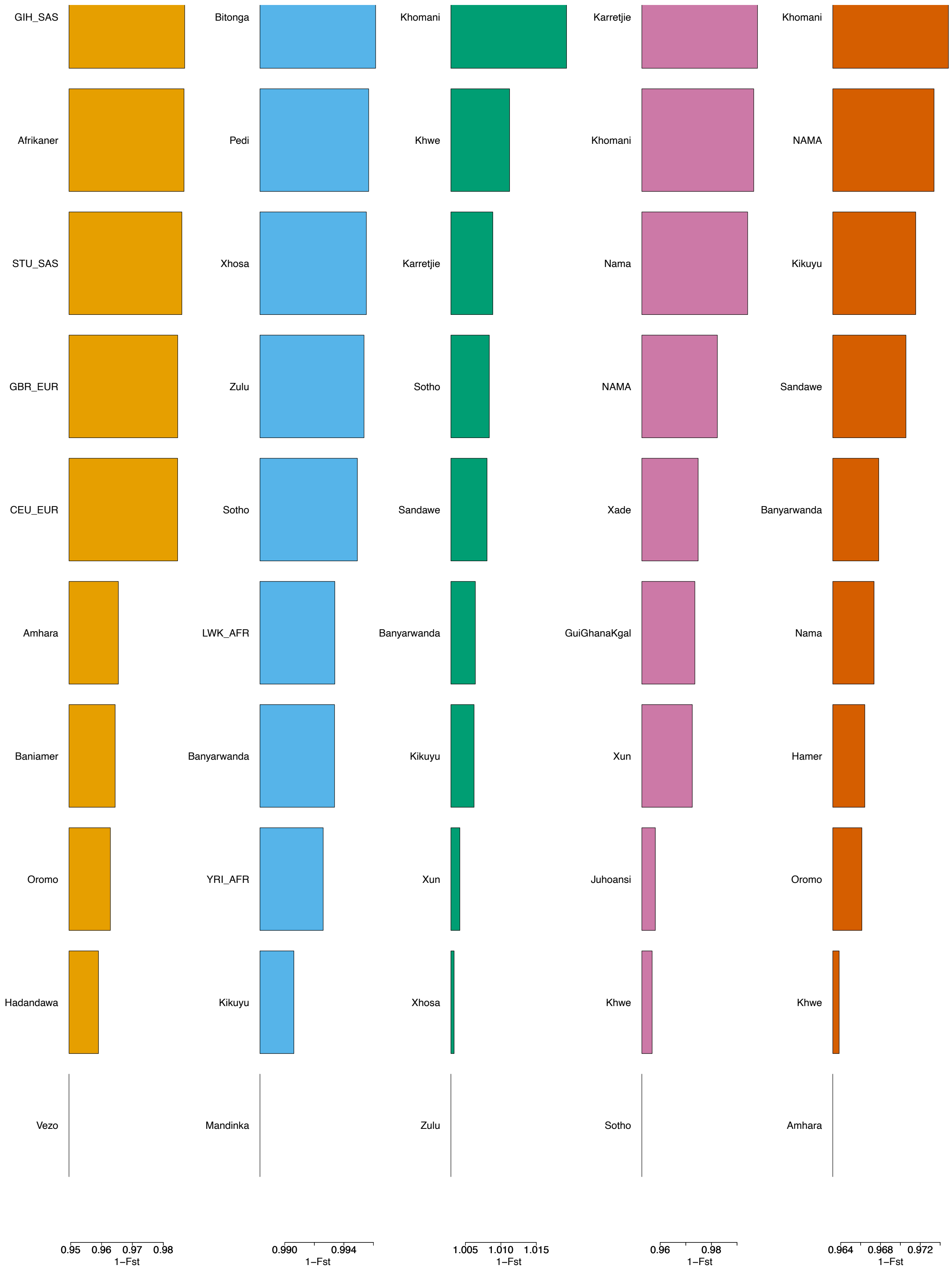


Supplementary Figure 32: 1-$F_{st}$ values from MOSAIC for the 5 constructed ancestries for Heidelberg.

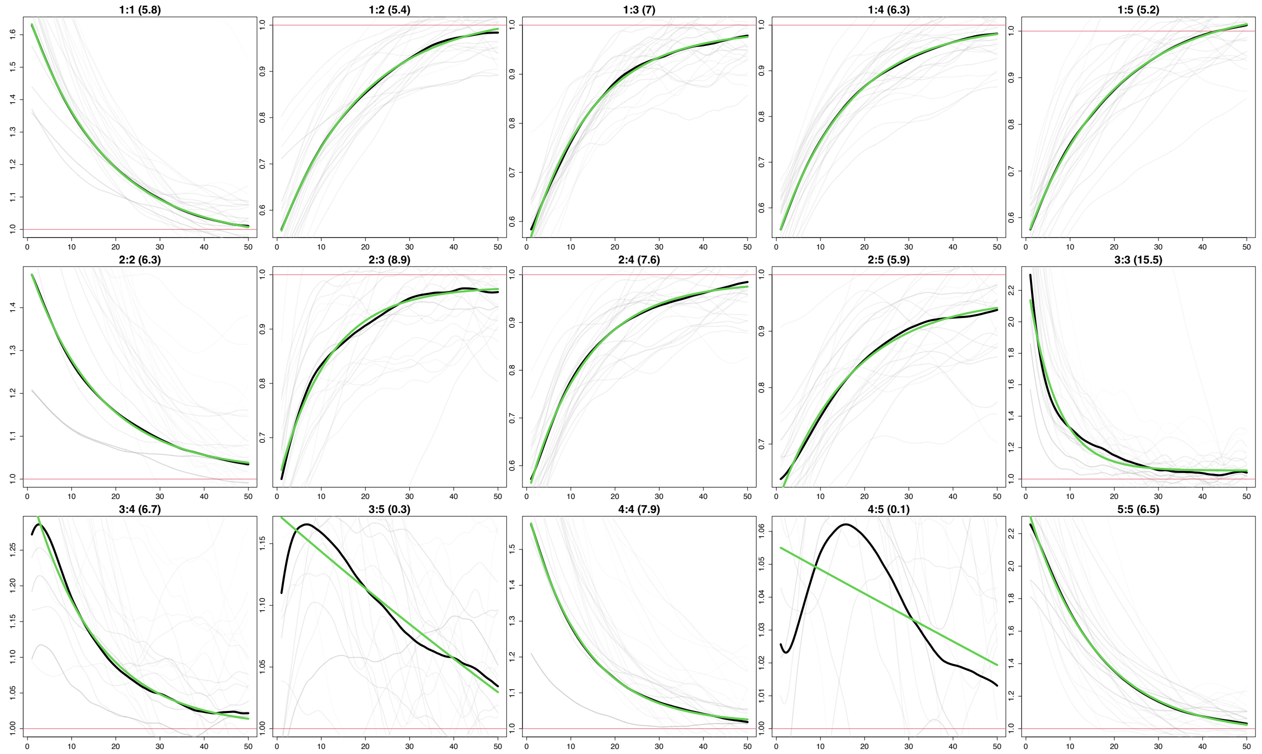


Supplementary Figure 33: Co-Ancestry curves from MOSAIC for the 5 constructed ancestries for Heidelberg. The y-axis depicts the relative probability of jointly copying two chunks from the donor populations and x-axis is the genetic distance in centimorgans. The black lines represent the empirical coancestry curves across all target individuals, the light grey are per individual, and the green lines represent the fitted single-event coancestry curve. Above each plot, the number of the populations being examined is noted as a:b, with the number of generations since the admixture event in parentheses behind it. The number represents the order the ancestries appear in in the 1-$F_{st}$ plot.

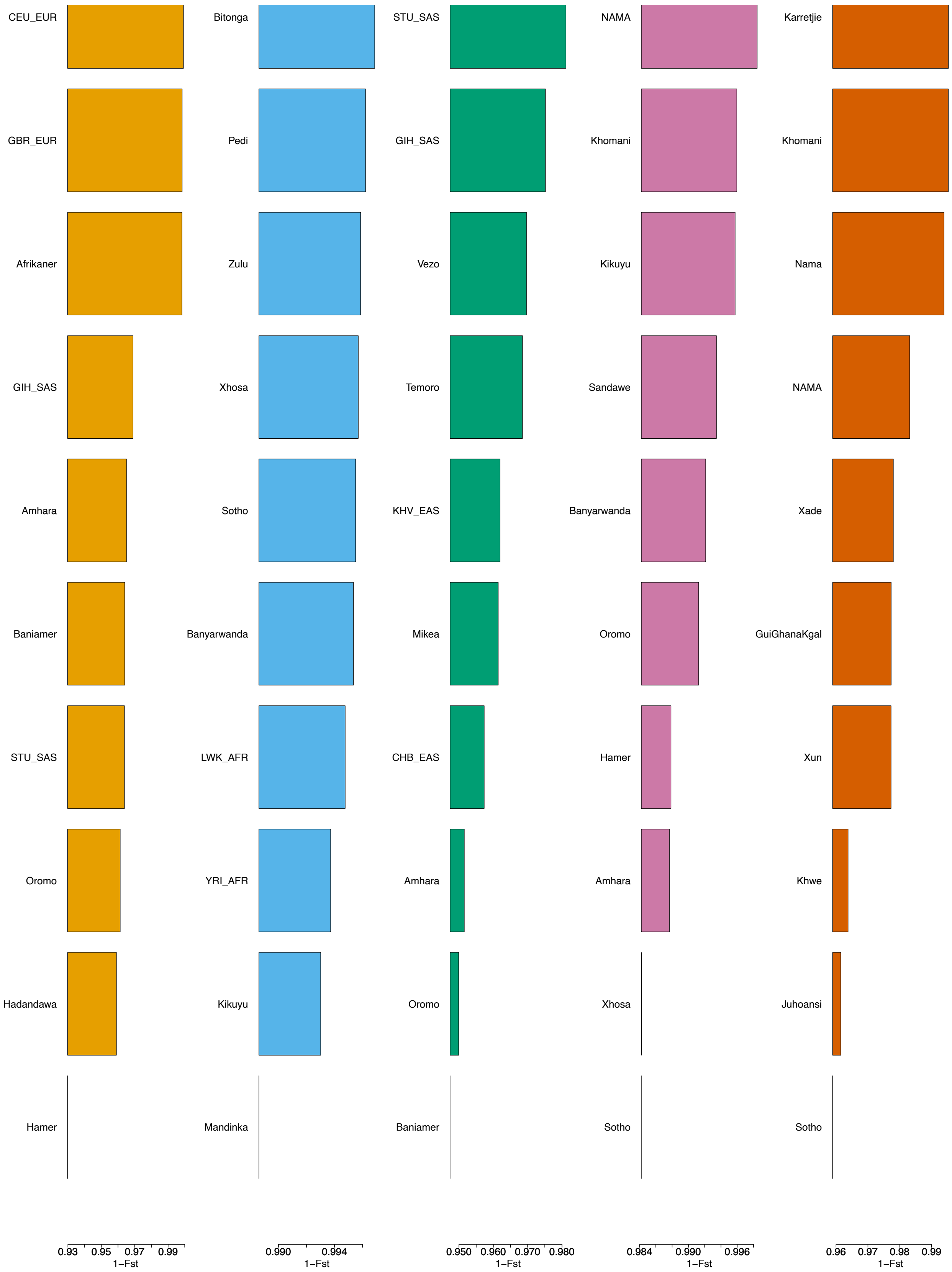


Supplementary Figure 34: 1-$F_{st}$ values from MOSAIC for the 5 constructed ancestries for Kranshoek.

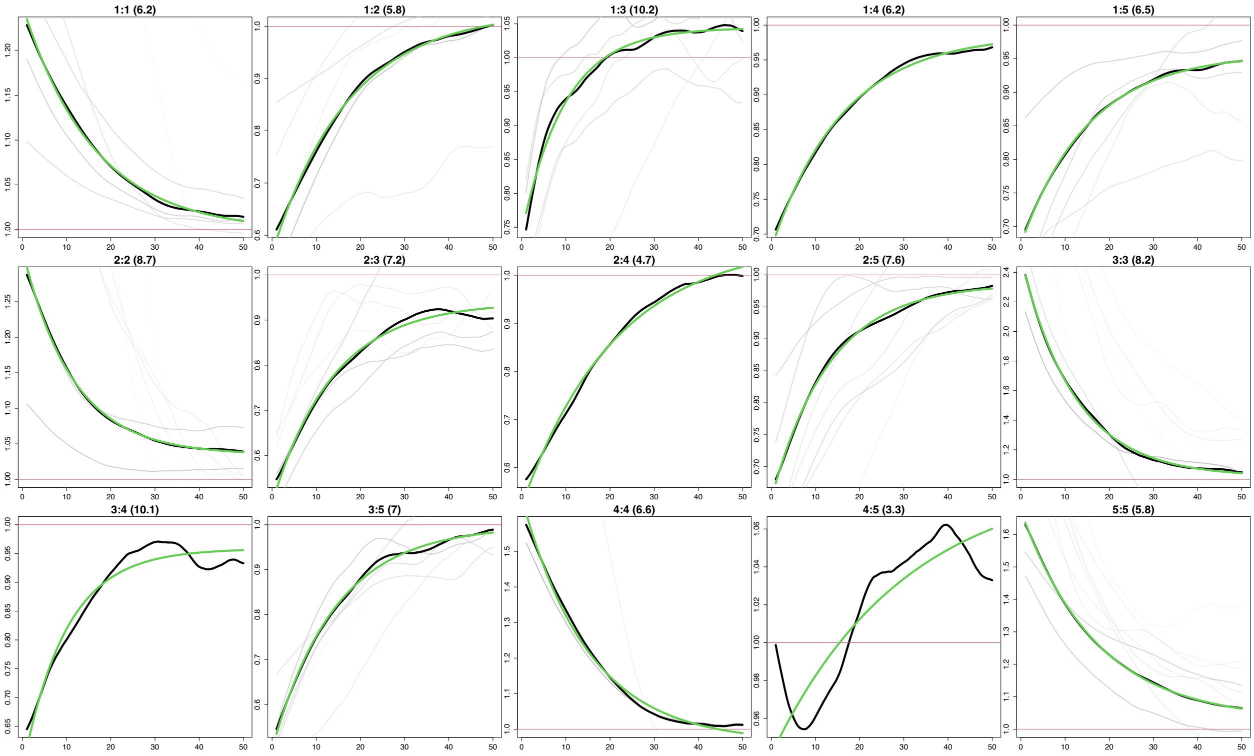


Supplementary Figure 35: Co-Ancestry curves from MOSAIC for the 5 constructed ancestries for Kranshoek. The y-axis depicts the relative probability of jointly copying two chunks from the donor populations and x-axis is the genetic distance in centimorgans. The black lines represent the empirical coancestry curves across all target individuals, the light grey are per individual, and the green lines represent the fitted single-event coancestry curve. Above each plot, the number of the populations being examined is noted as a:b, with the number of generations since the admixture event in parentheses behind it. The number represents the order the ancestries appear in in the 1-$F_{st}$ plot.

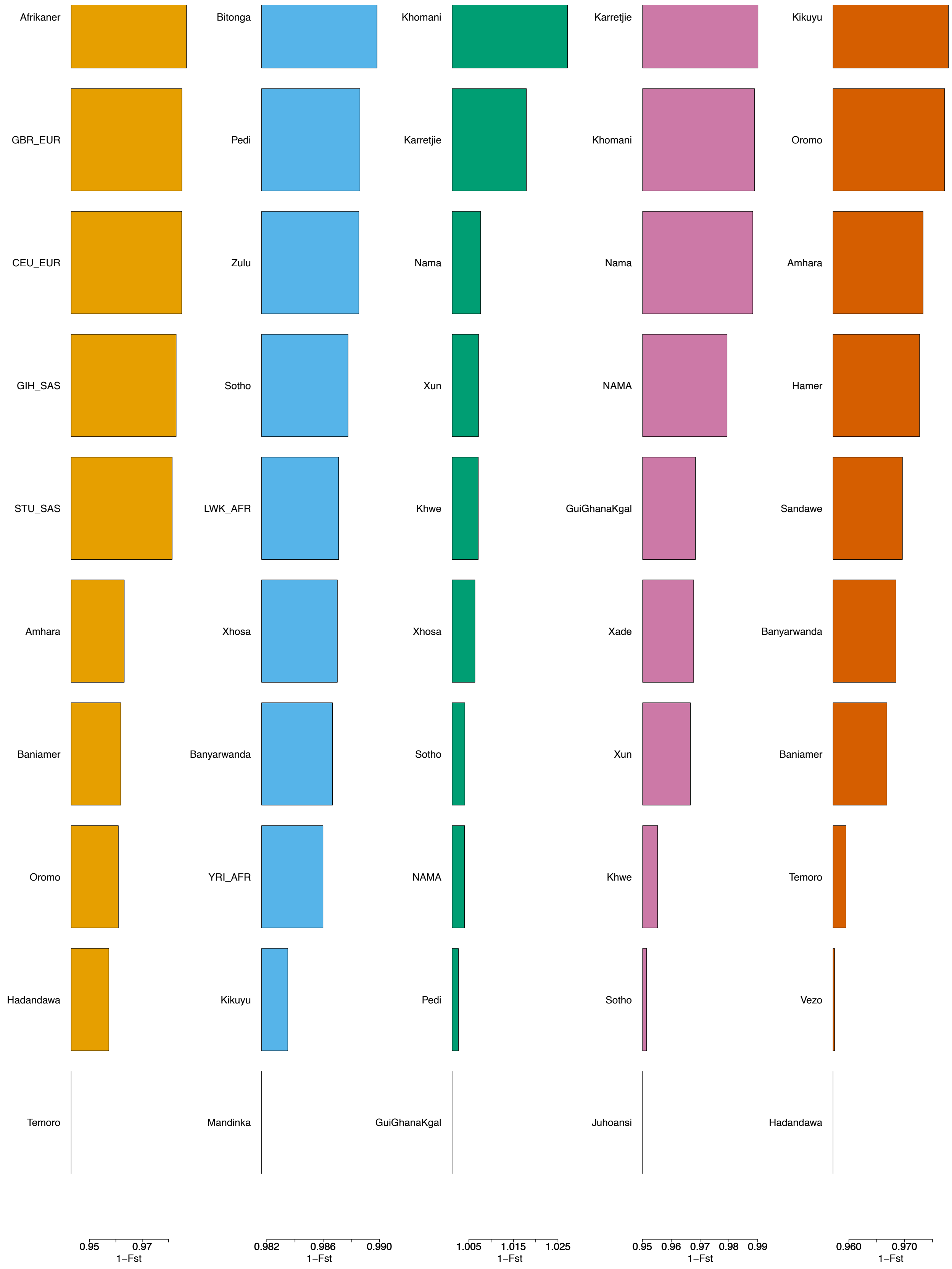


Supplementary Figure 36: 1-$F_{st}$ values from MOSAIC for the 5 constructed ancestries for Melkhoutfontein.

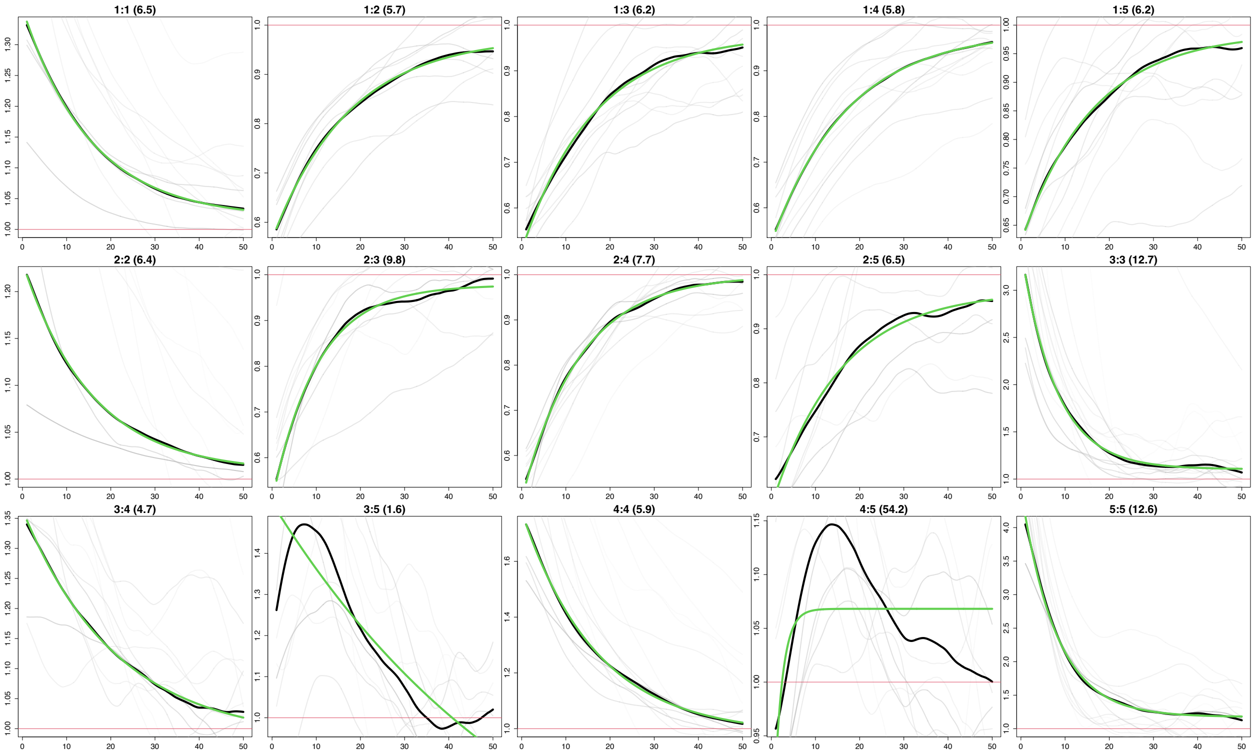


Supplementary Figure 37: Co-Ancestry curves from MOSAIC for the 5 constructed ancestries for Melkhoutfontein. The y-axis depicts the relative probability of jointly copying two chunks from the donor populations and x-axis is the genetic distance in centimorgans. The black lines represent the empirical coancestry curves across all target individuals, the light grey are per individual, and the green lines represent the fitted single-event coancestry curve. Above each plot, the number of the populations being examined is noted as a:b, with the number of generations since the admixture event in parentheses behind it. The number represents the order the ancestries appear in in the 1-$F_{st}$ plot.

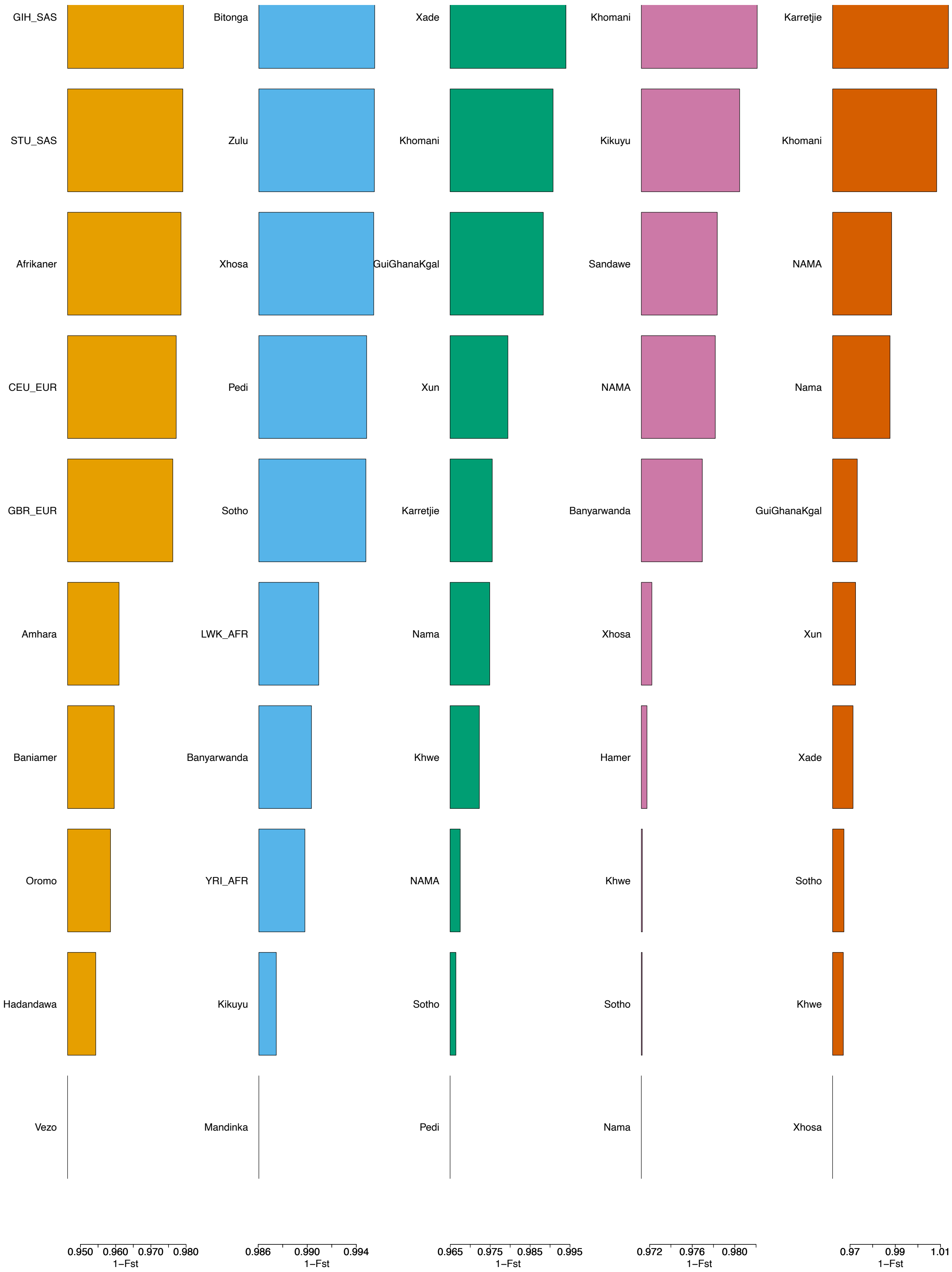


Supplementary Figure 38: 1-$F_{st}$ values from MOSAIC for the 5 constructed ancestries for Nieu-Bethesda.

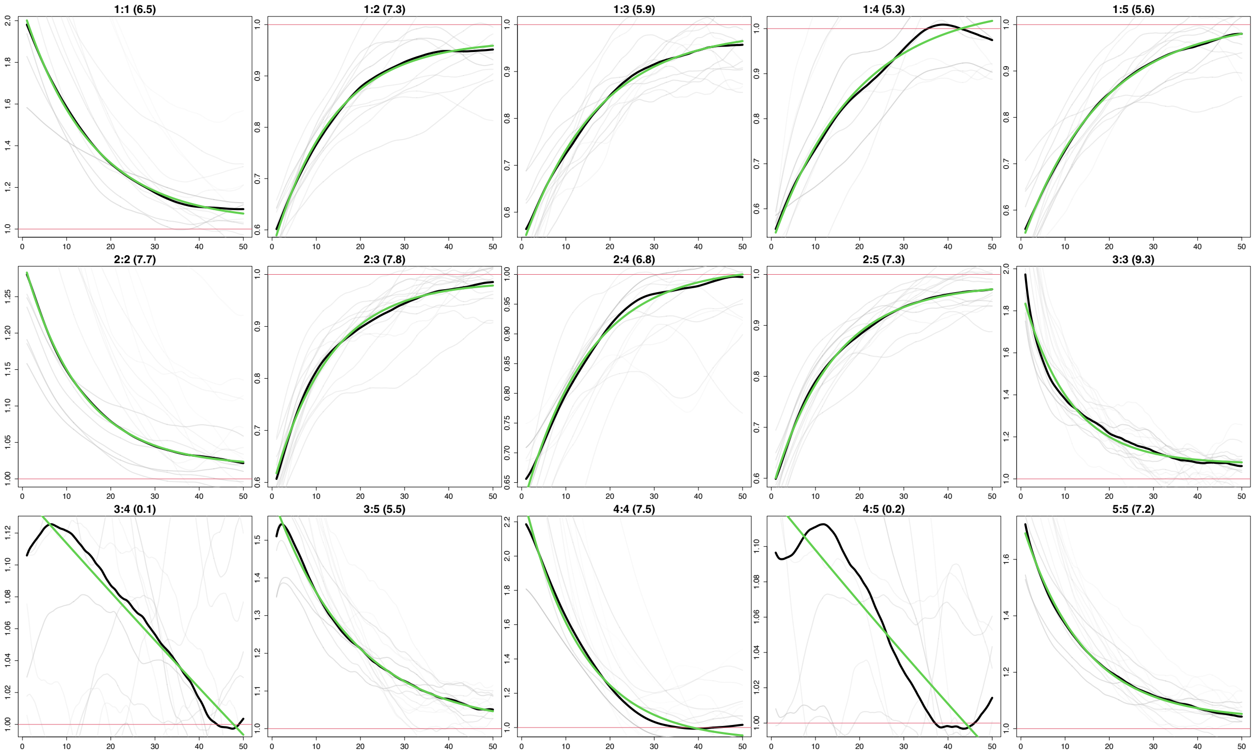


Supplementary Figure 39: Co-Ancestry curves from MOSAIC for the 5 constructed ancestries for Nieu-Bethesda. The y-axis depicts the relative probability of jointly copying two chunks from the donor populations and x-axis is the genetic distance in centimorgans. The black lines represent the empirical coancestry curves across all target individuals, the light grey are per individual, and the green lines represent the fitted single-event coancestry curve. Above each plot, the number of the populations being examined is noted as a:b, with the number of generations since the admixture event in parentheses behind it. The number represents the order the ancestries appear in in the 1-$F_{st}$ plot.

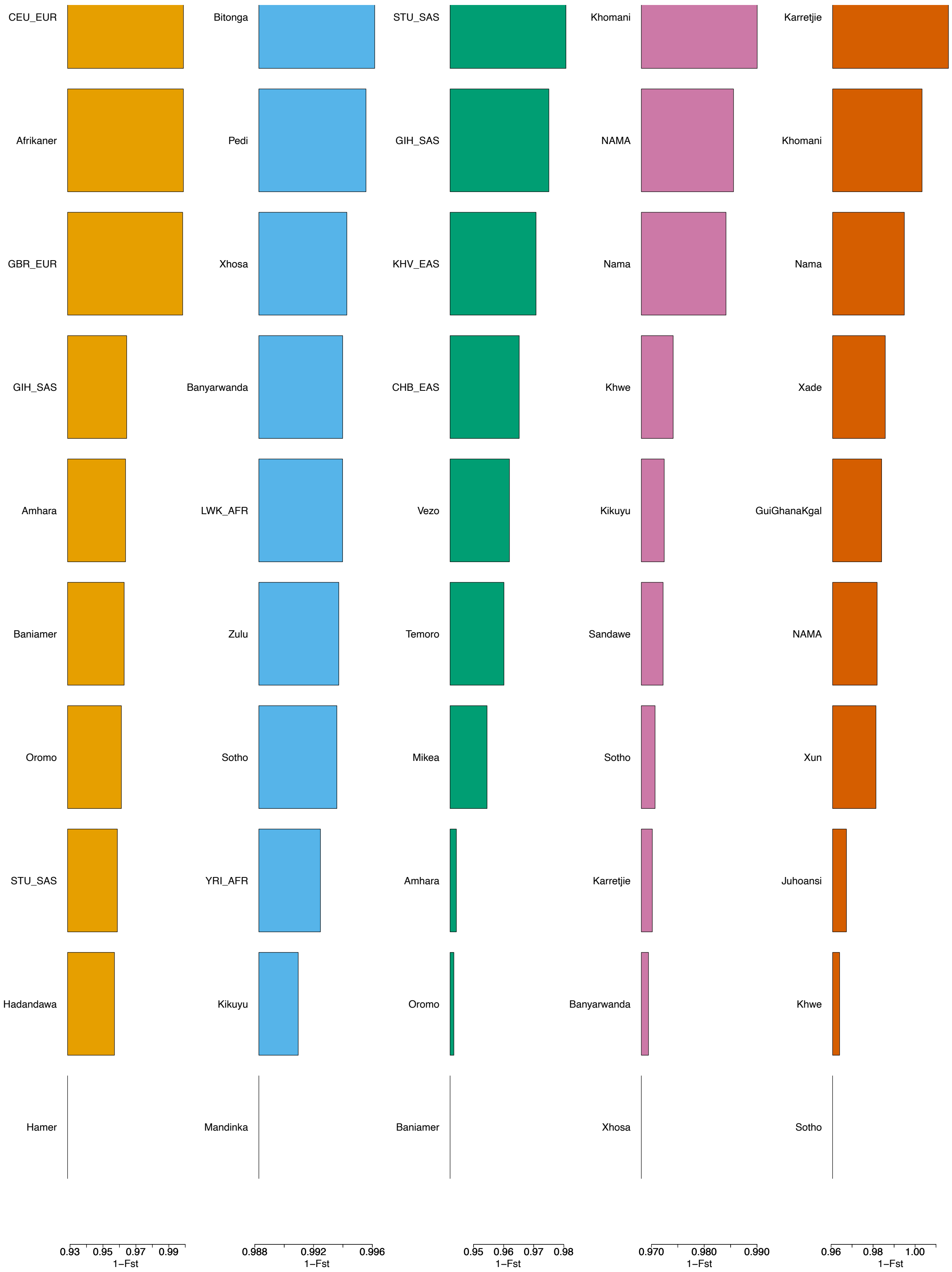


Supplementary Figure 40: 1-$F_{st}$ values from MOSAIC for the 5 constructed ancestries for Oudtshoorn.

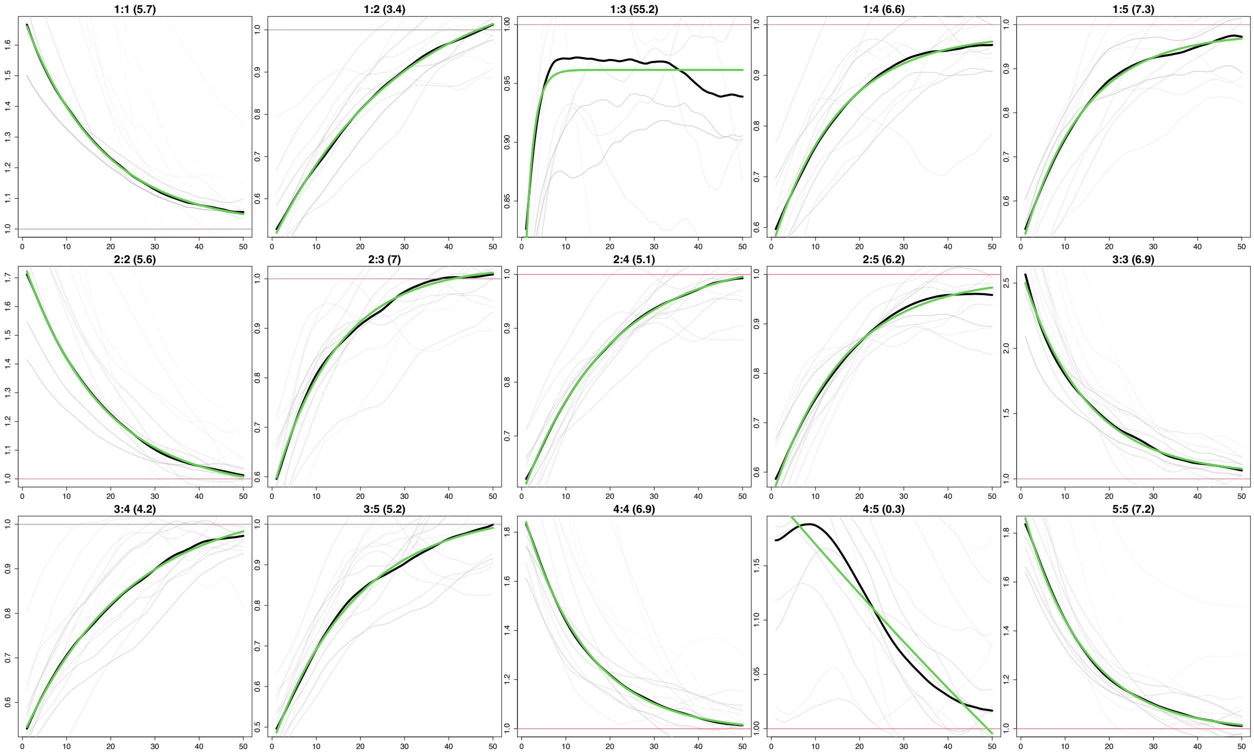


Supplementary Figure 41: Co-Ancestry curves from MOSAIC for the 5 constructed ancestries for Oudtshoorn. The y-axis depicts the relative probability of jointly copying two chunks from the donor populations and x-axis is the genetic distance in centimorgans. The black lines represent the empirical coancestry curves across all target individuals, the light grey are per individual, and the green lines represent the fitted single-event coancestry curve. Above each plot, the number of the populations being examined is noted as a:b, with the number of generations since the admixture event in parentheses behind it. The number represents the order the ancestries appear in in the 1-$F_{st}$ plot.

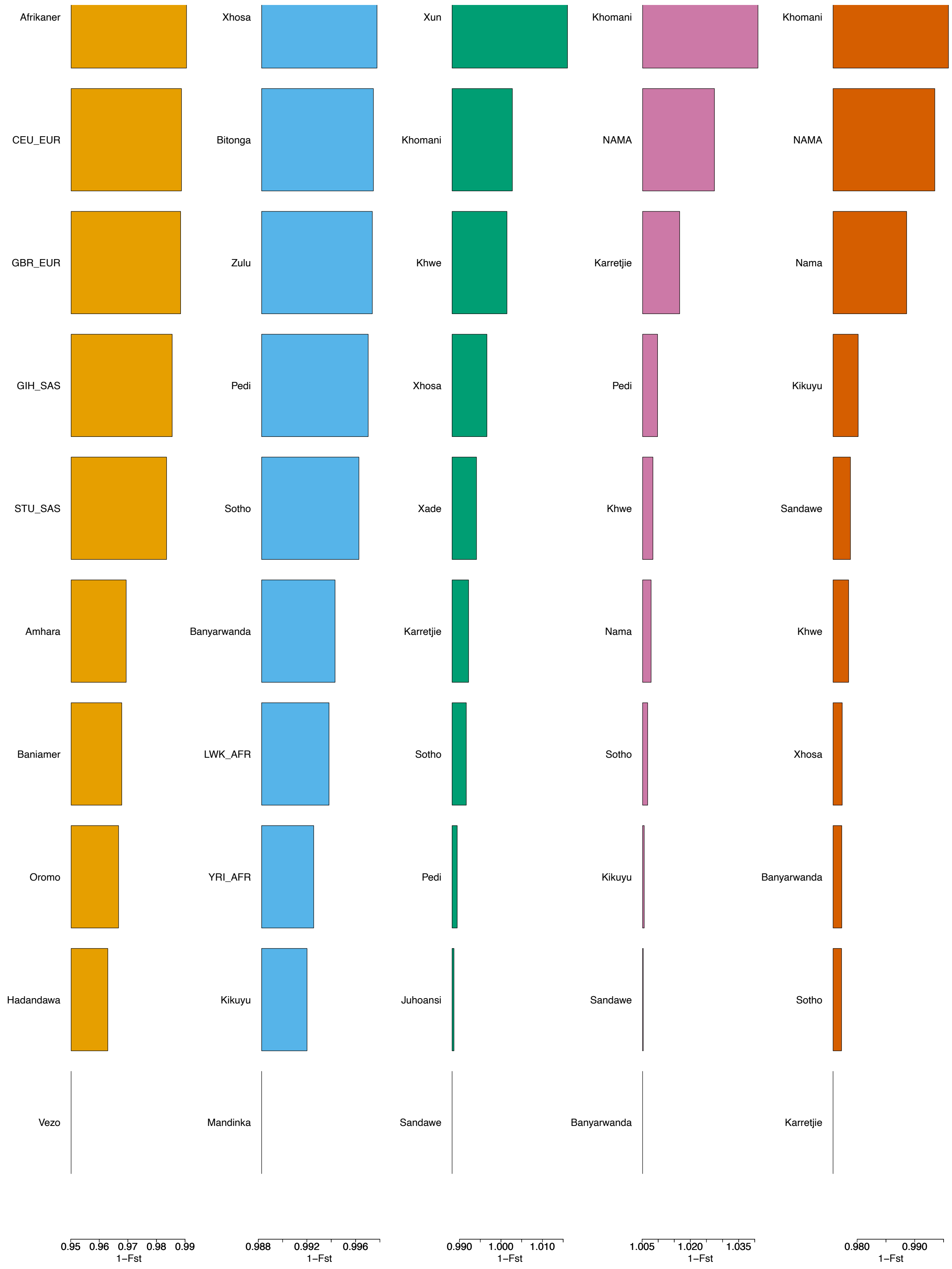


Supplementary Figure 42: 1-$F_{st}$ values from MOSAIC for the 5 constructed ancestries for Prince Albert.

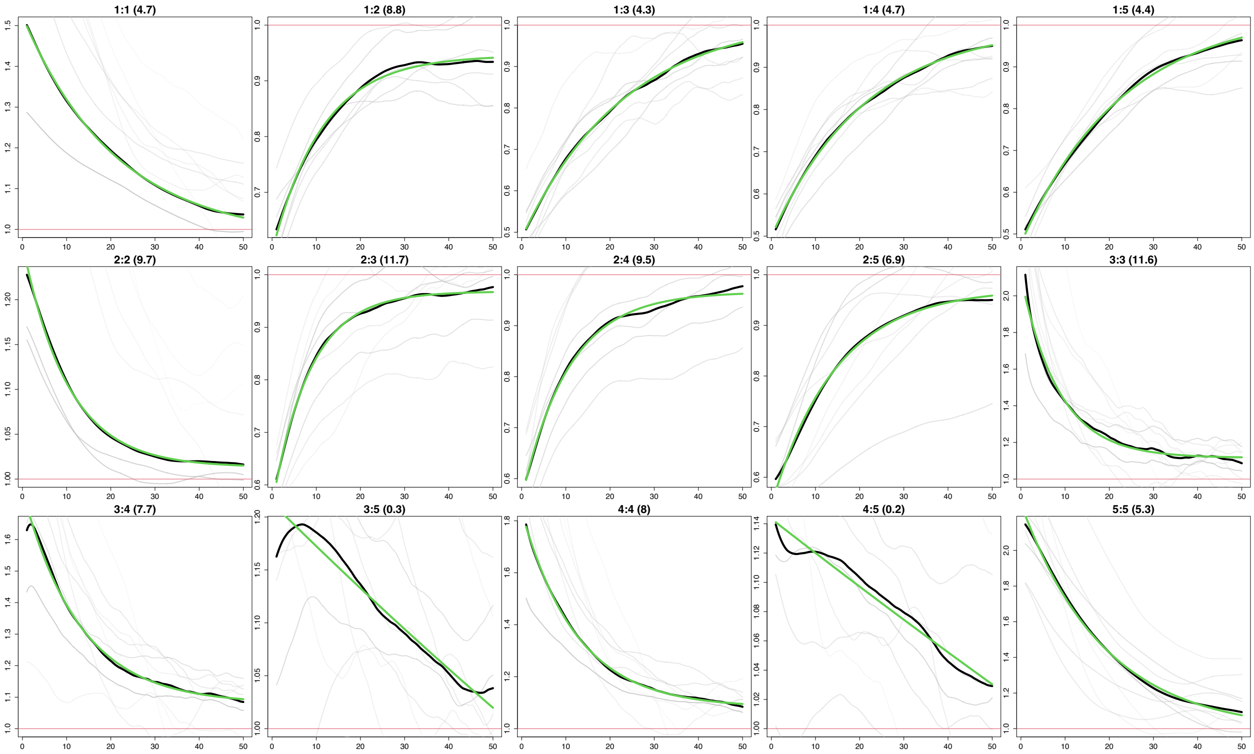


Supplementary Figure 43: Co-Ancestry curves from MOSAIC for the 5 constructed ancestries for Prince Albert. The y-axis depicts the relative probability of jointly copying two chunks from the donor populations and x-axis is the genetic distance in centimorgans. The black lines represent the empirical coancestry curves across all target individuals, the light grey are per individual, and the green lines represent the fitted single-event coancestry curve. Above each plot, the number of the populations being examined is noted as a:b, with the number of generations since the admixture event in parentheses behind it. The number represents the order the ancestries appear in in the 1-$F_{st}$ plot.

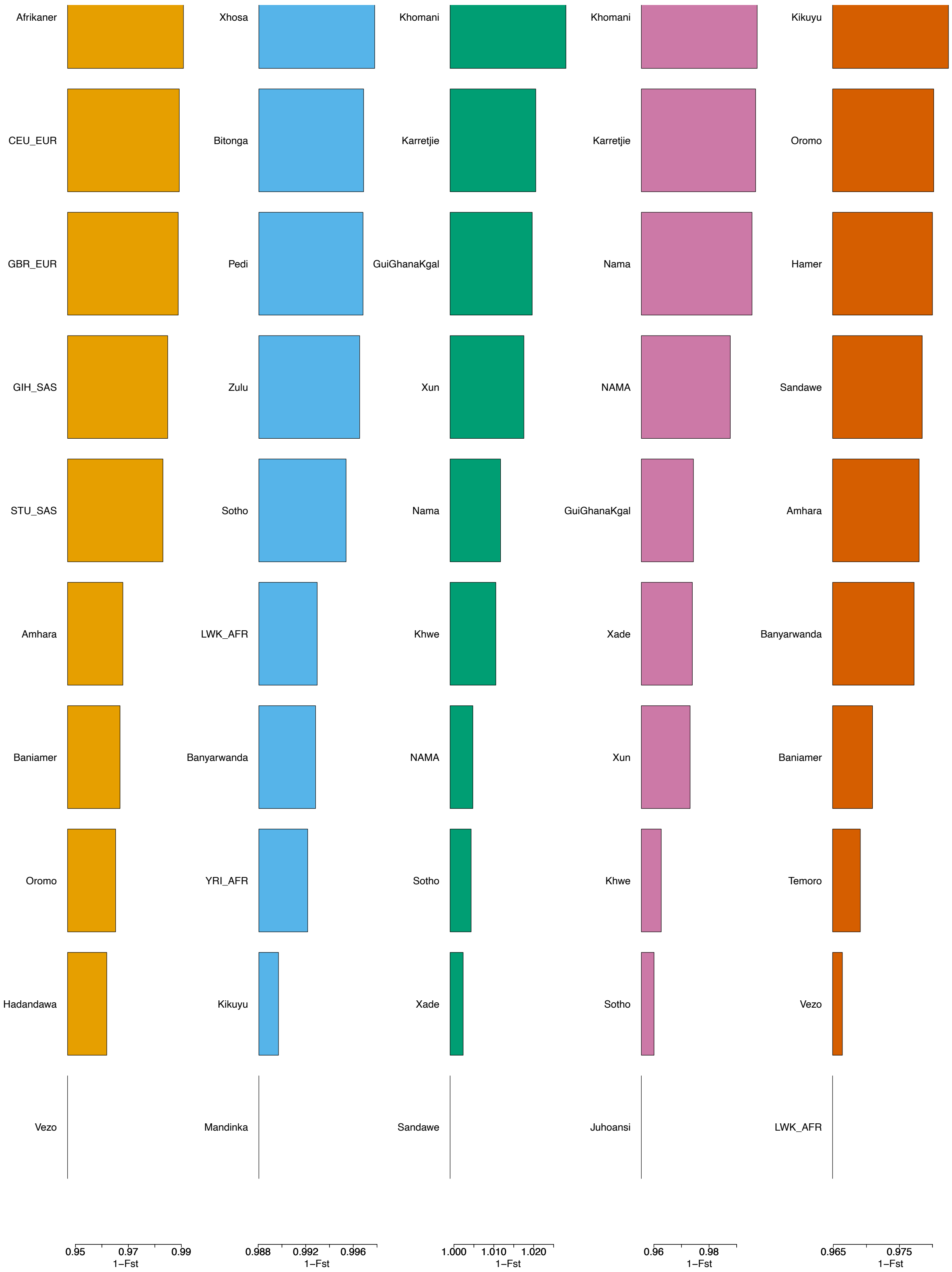


Supplementary Figure 44: 1-$F_{st}$ values from MOSAIC for the 5 constructed ancestries for Railton.

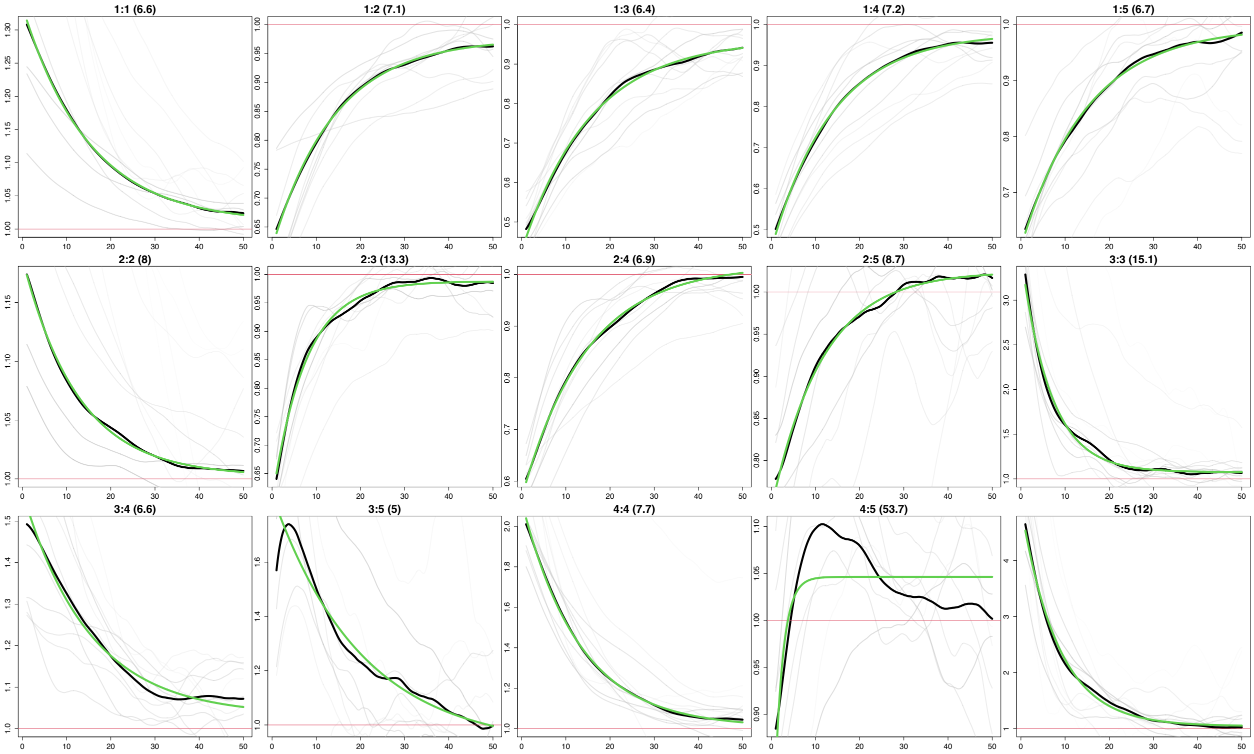


Supplementary Figure 45: Co-Ancestry curves from MOSAIC for the 5 constructed ancestries for Railton. The y-axis depicts the relative probability of jointly copying two chunks from the donor populations and x-axis is the genetic distance in centimorgans. The black lines represent the empirical coancestry curves across all target individuals, the light grey are per individual, and the green lines represent the fitted single-event coancestry curve. Above each plot, the number of the populations being examined is noted as a:b, with the number of generations since the admixture event in parentheses behind it. The number represents the order the ancestries appear in in the 1-$F_{st}$ plot.

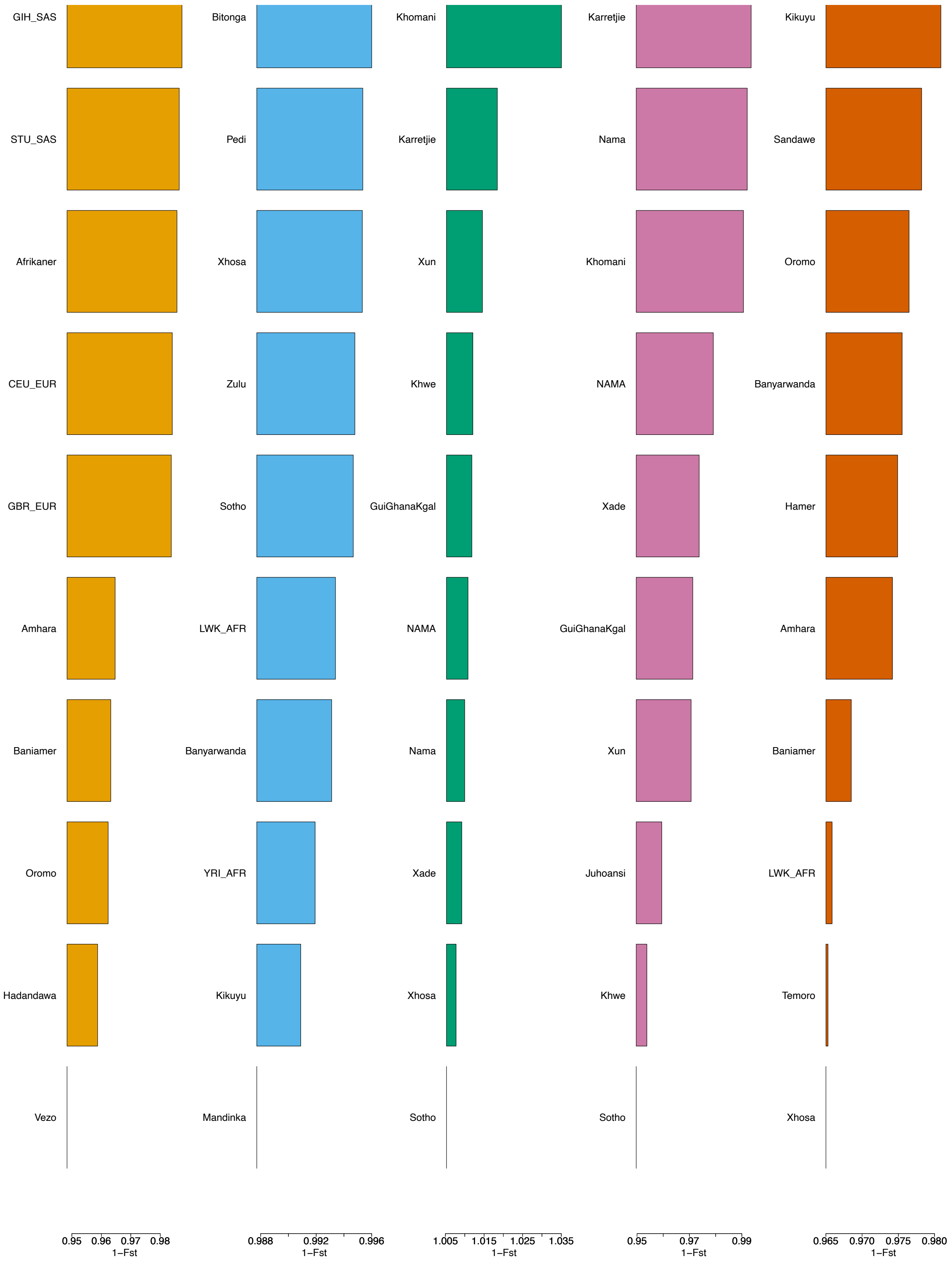


Supplementary Figure 46: 1-$F_{st}$ values from MOSAIC for the 5 constructed ancestries for Riversdale.

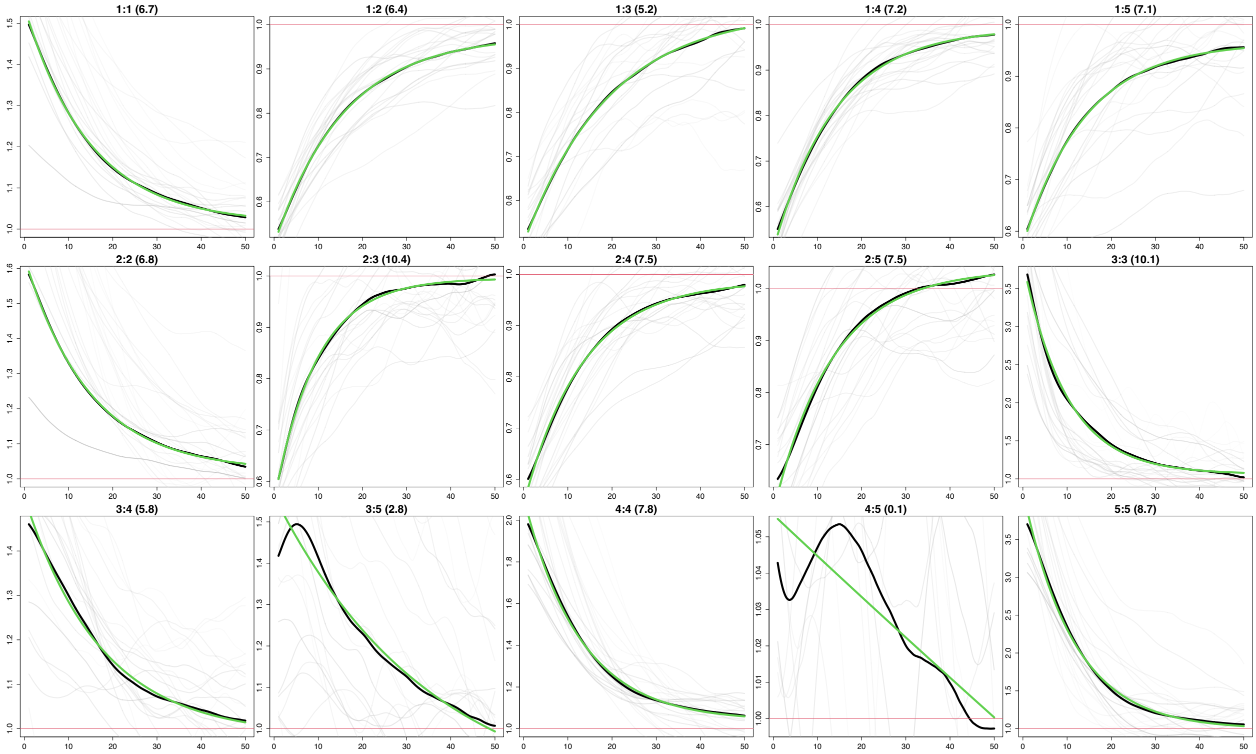


Supplementary Figure 47: Co-Ancestry curves from MOSAIC for the 5 constructed ancestries for Riversdale. The y-axis depicts the relative probability of jointly copying two chunks from the donor populations and x-axis is the genetic distance in centimorgans. The black lines represent the empirical coancestry curves across all target individuals, the light grey are per individual, and the green lines represent the fitted single-event coancestry curve. Above each plot, the number of the populations being examined is noted as a:b, with the number of generations since the admixture event in parentheses behind it. The number represents the order the ancestries appear in in the 1-$F_{st}$ plot.

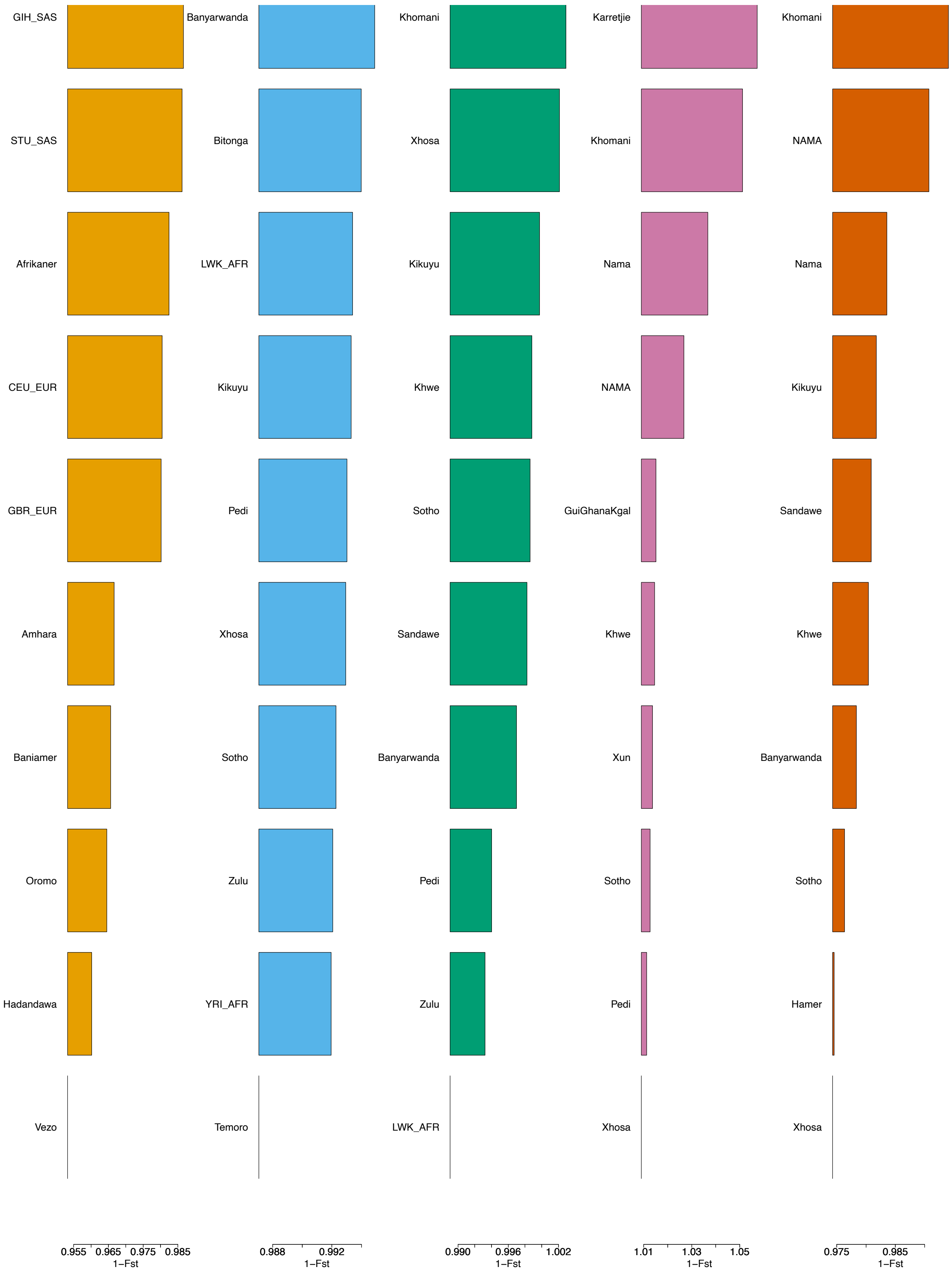


Supplementary Figure 48: 1-$F_{st}$ values from MOSAIC for the 5 constructed ancestries for RotterdamFarm.

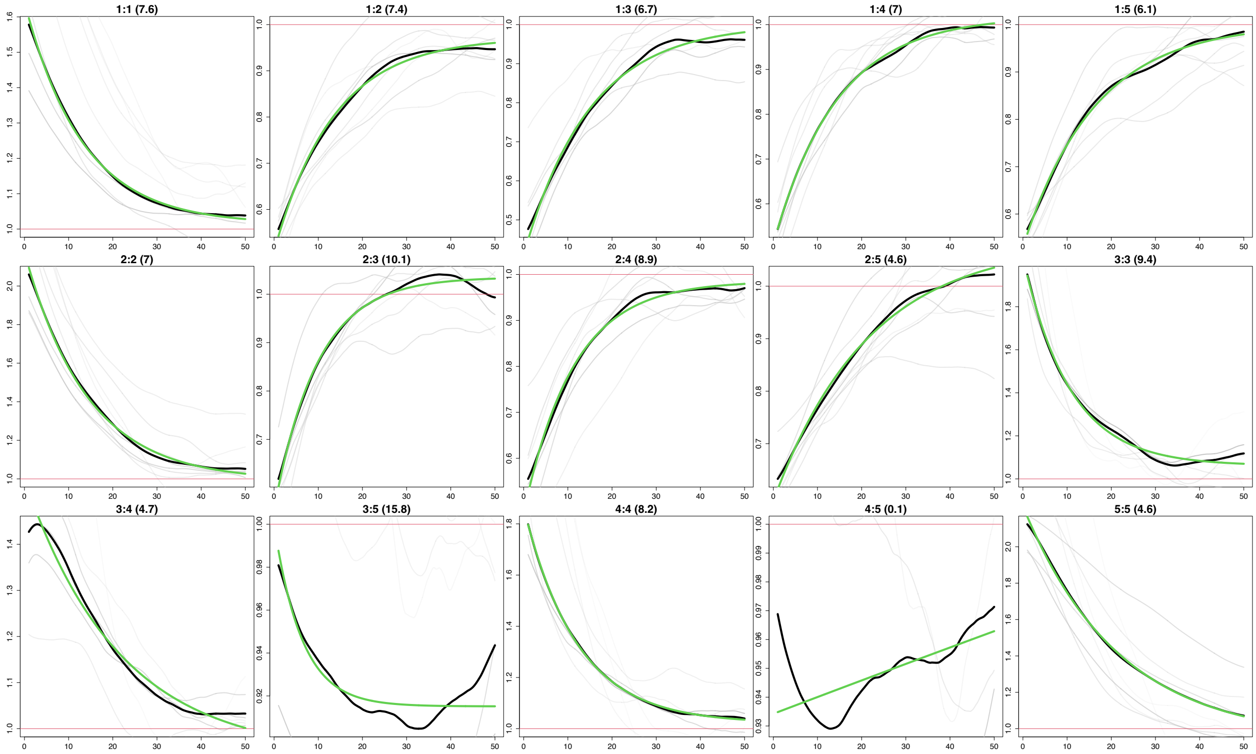


Supplementary Figure 49: Co-Ancestry curves from MOSAIC for the 5 constructed ancestries for Rotterdam-Farm. The y-axis depicts the relative probability of jointly copying two chunks from the donor populations and x-axis is the genetic distance in centimorgans. The black lines represent the empirical coancestry curves across all target individuals, the light grey are per individual, and the green lines represent the fitted single-event coancestry curve. Above each plot, the number of the populations being examined is noted as a:b, with the number of generations since the admixture event in parentheses behind it. The number represents the order the ancestries appear in in the 1-$F_{st}$ plot.

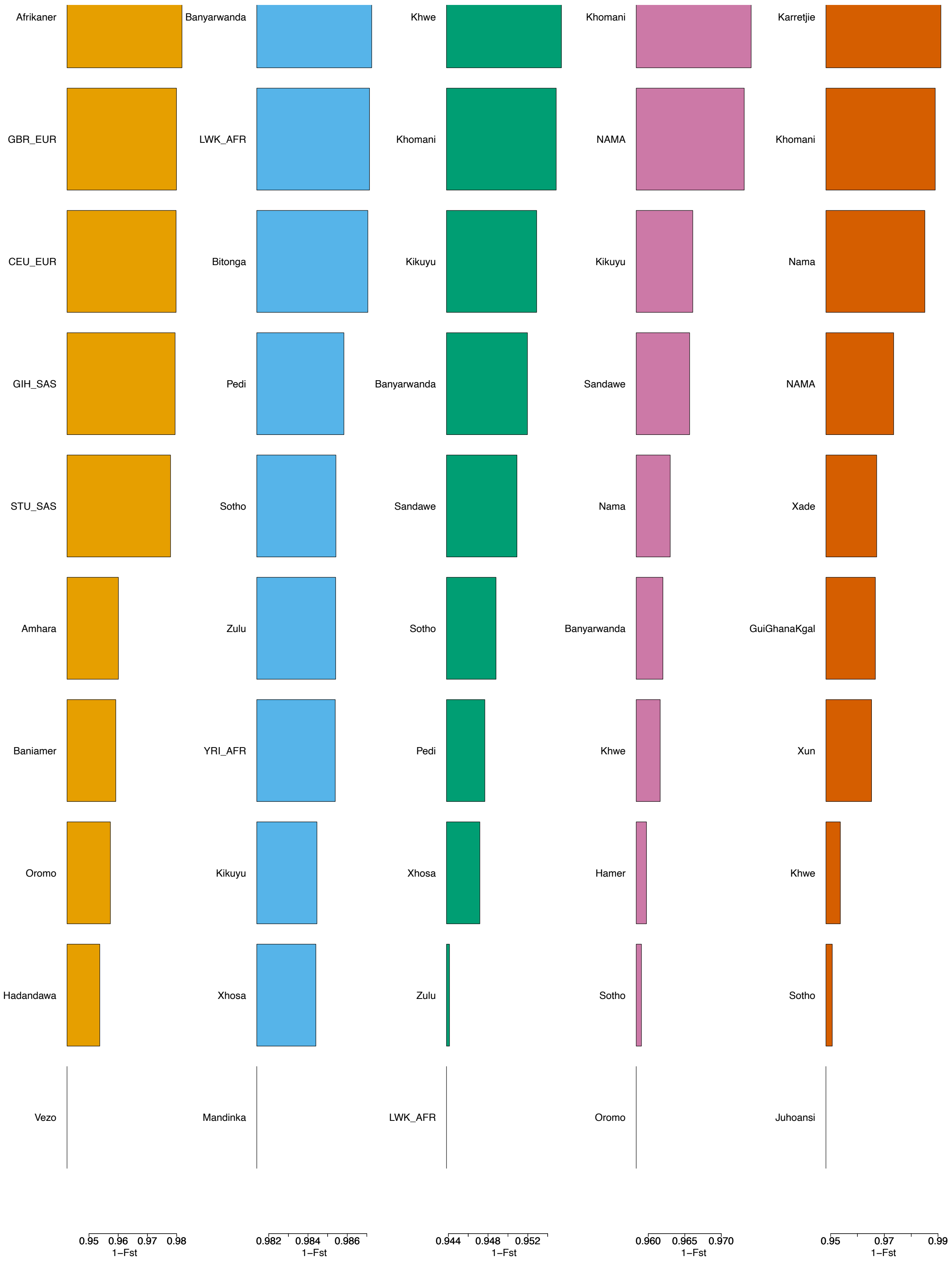


Supplementary Figure 50: 1-$F_{st}$ values from MOSAIC for the 5 constructed ancestries for Slangriver.

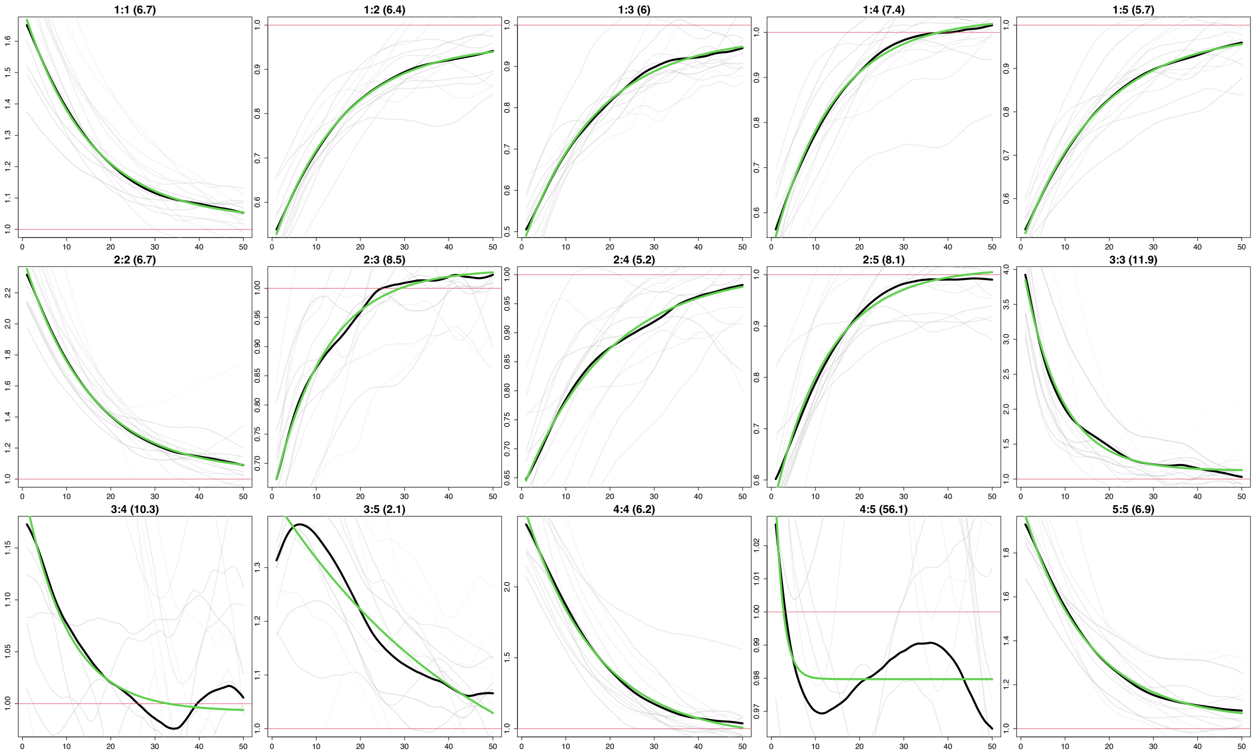


Supplementary Figure 51: Co-Ancestry curves from MOSAIC for the 5 constructed ancestries for Slangriver. The y-axis depicts the relative probability of jointly copying two chunks from the donor populations and x-axis is the genetic distance in centimorgans. The black lines represent the empirical coancestry curves across all target individuals, the light grey are per individual, and the green lines represent the fitted single-event coancestry curve. Above each plot, the number of the populations being examined is noted as a:b, with the number of generations since the admixture event in parentheses behind it. The number represents the order the ancestries appear in in the 1-$F_{st}$ plot.

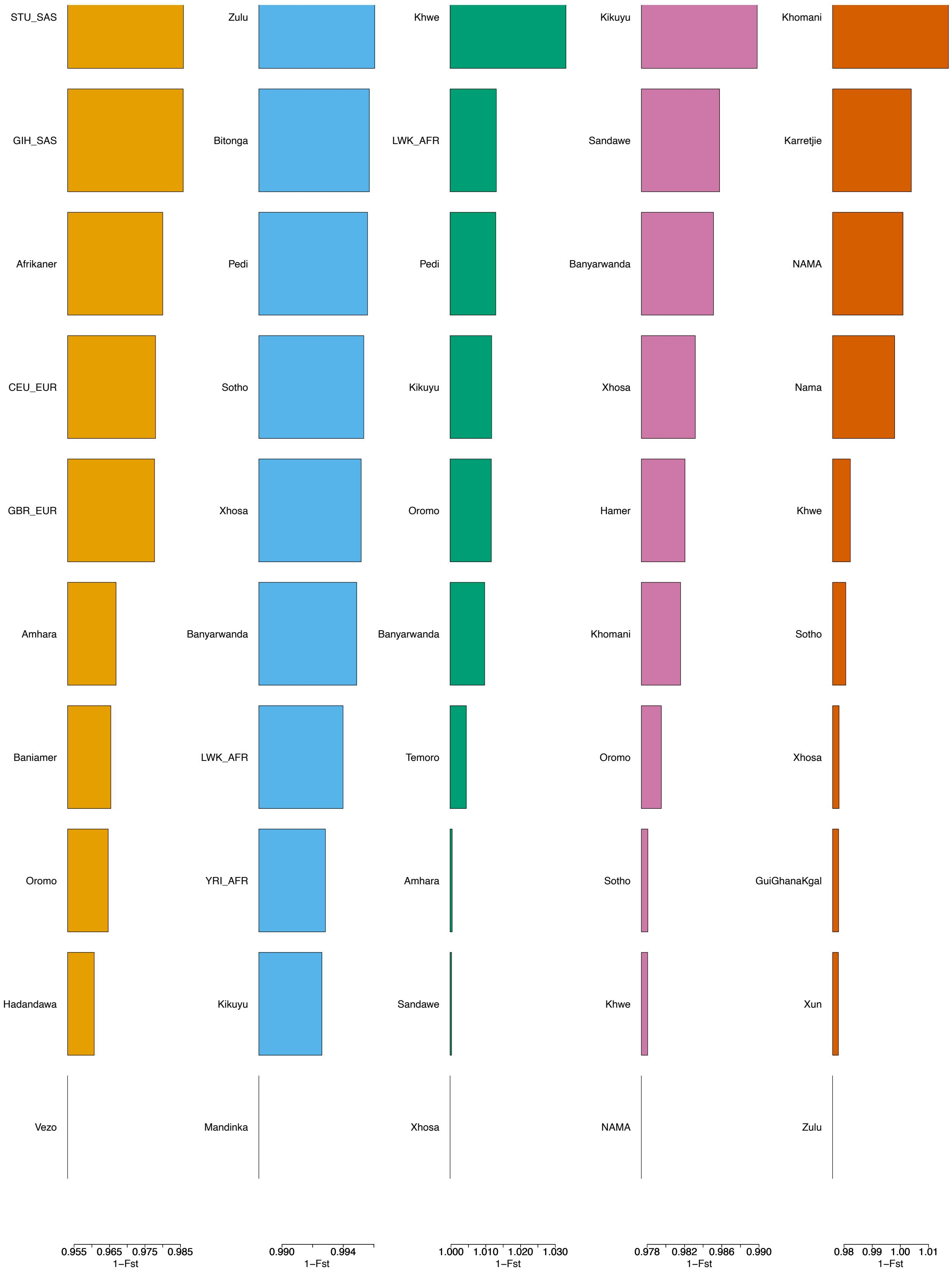


Supplementary Figure 52: 1-$F_{st}$ values from MOSAIC for the 5 constructed ancestries for Stormsvlei.

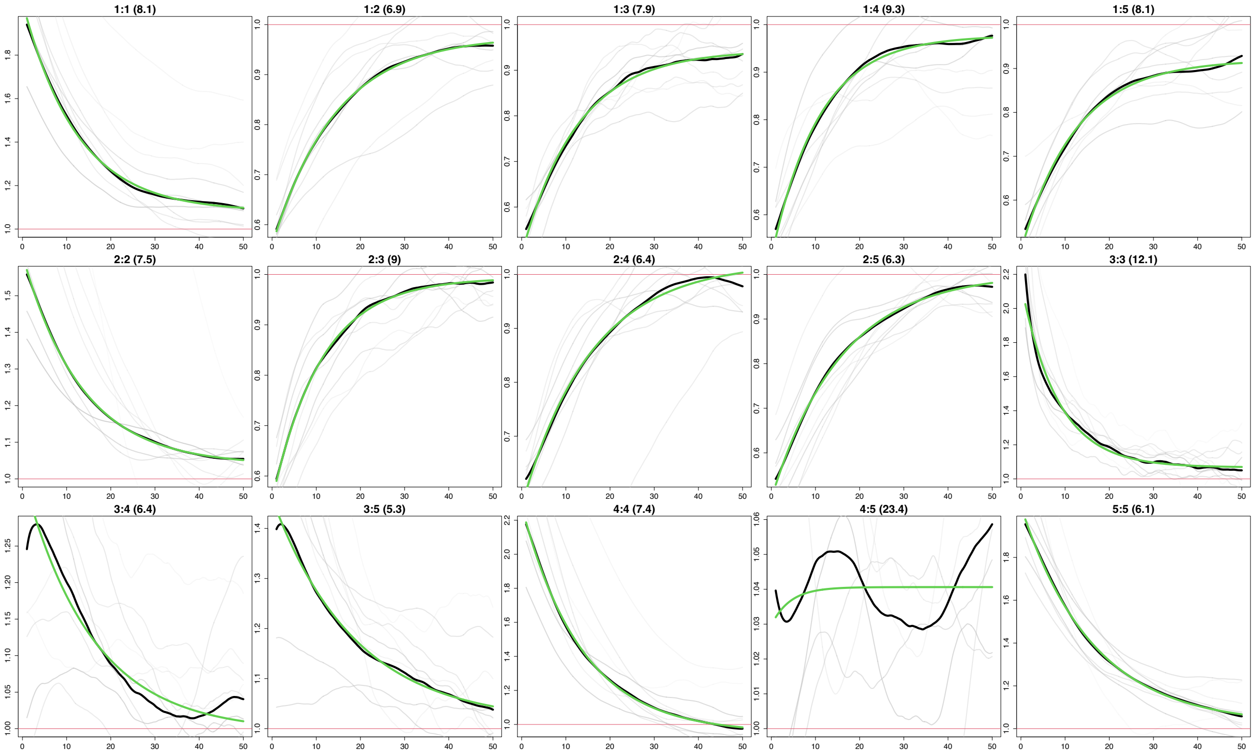


Supplementary Figure 53: Co-Ancestry curves from MOSAIC for the 5 constructed ancestries for Stormsvlei. The y-axis depicts the relative probability of jointly copying two chunks from the donor populations and x-axis is the genetic distance in centimorgans. The black lines represent the empirical coancestry curves across all target individuals, the light grey are per individual, and the green lines represent the fitted single-event coancestry curve. Above each plot, the number of the populations being examined is noted as a:b, with the number of generations since the admixture event in parentheses behind it. The number represents the order the ancestries appear in in the 1-$F_{st}$ plot.

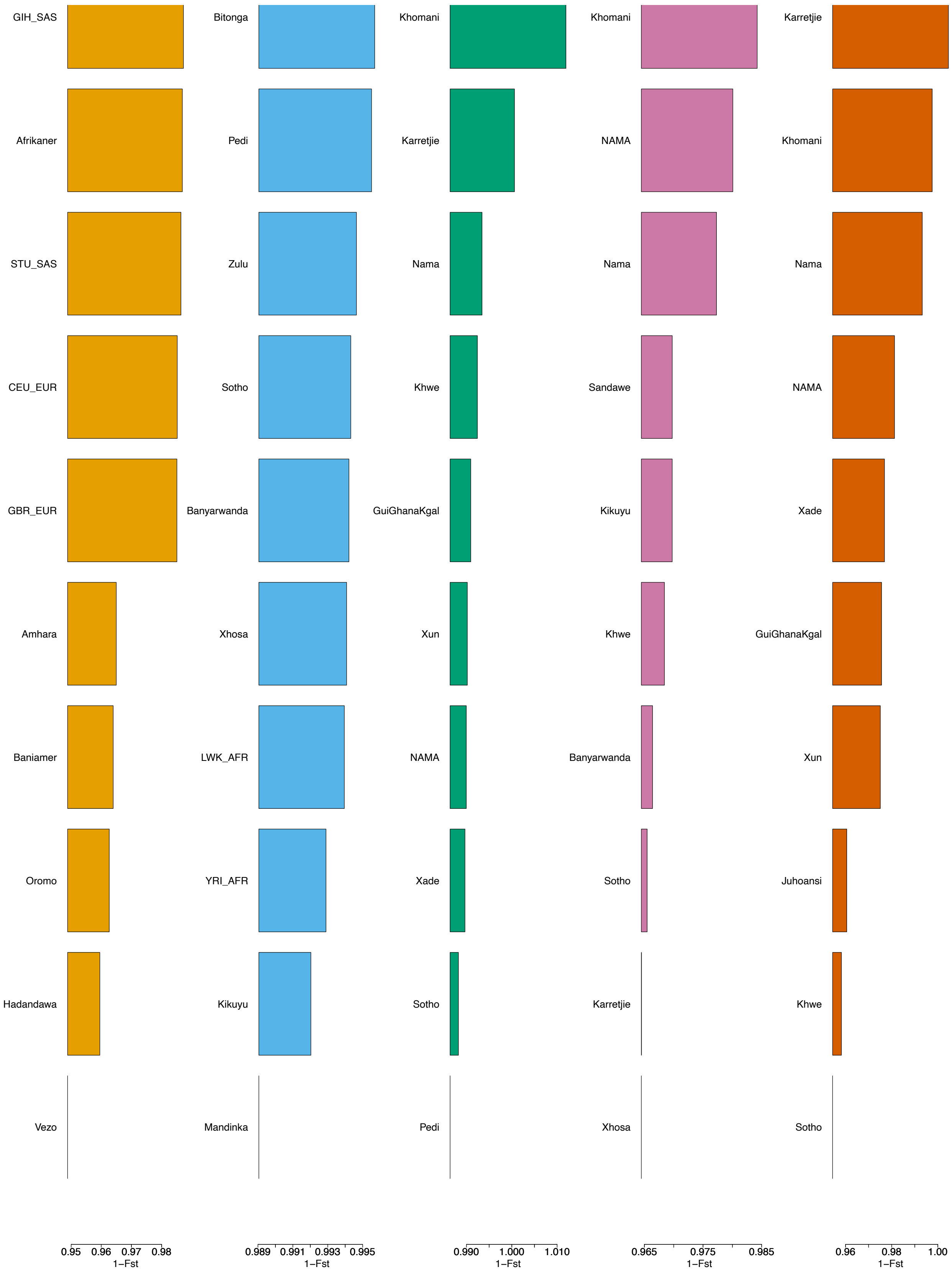


Supplementary Figure 54: Co-Ancestry curves from MOSAIC for the 5 constructed ancestries for Suurbraak.

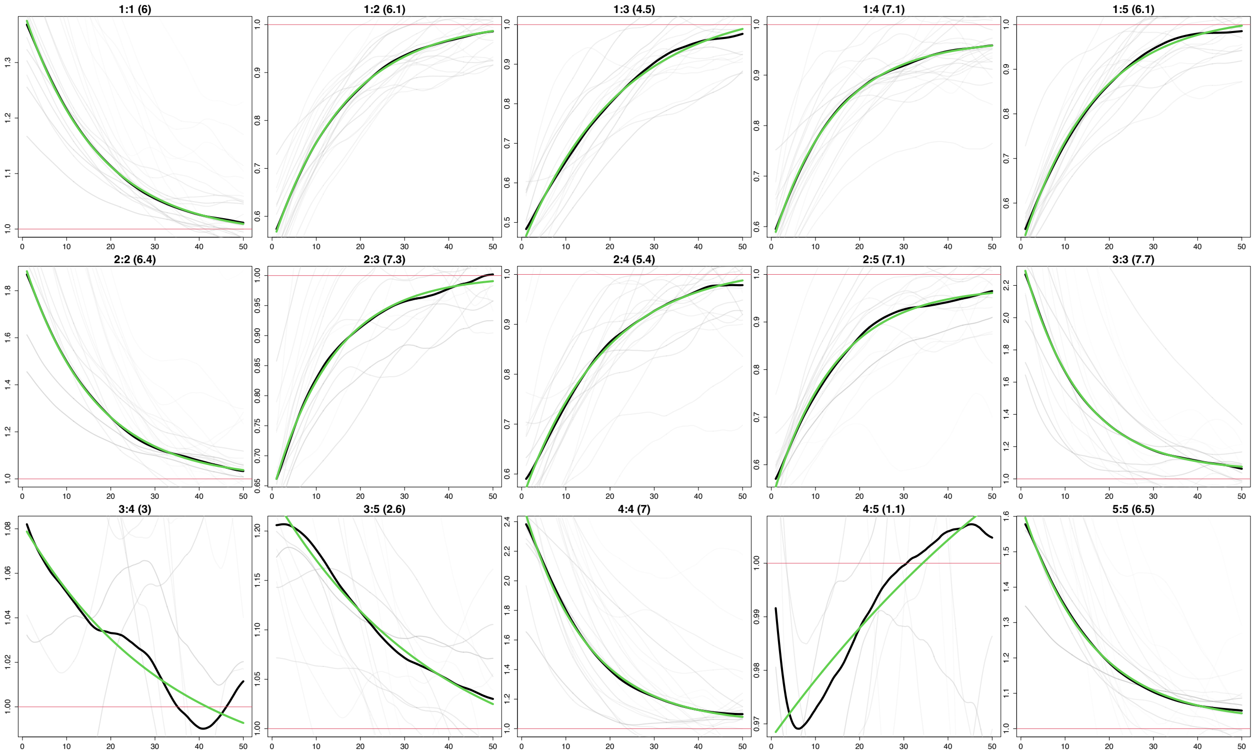


Supplementary Figure 55: Co-Ancestry curves from MOSAIC for the 5 constructed ancestries for Suurbraak. The y-axis depicts the relative probability of jointly copying two chunks from the donor populations and x-axis is the genetic distance in centimorgans. The black lines represent the empirical coancestry curves across all target individuals, the light grey are per individual, and the green lines represent the fitted single-event coancestry curve. Above each plot, the number of the populations being examined is noted as a:b, with the number of generations since the admixture event in parentheses behind it. The number represents the order the ancestries appear in in the 1-$F_{st}$ plot.

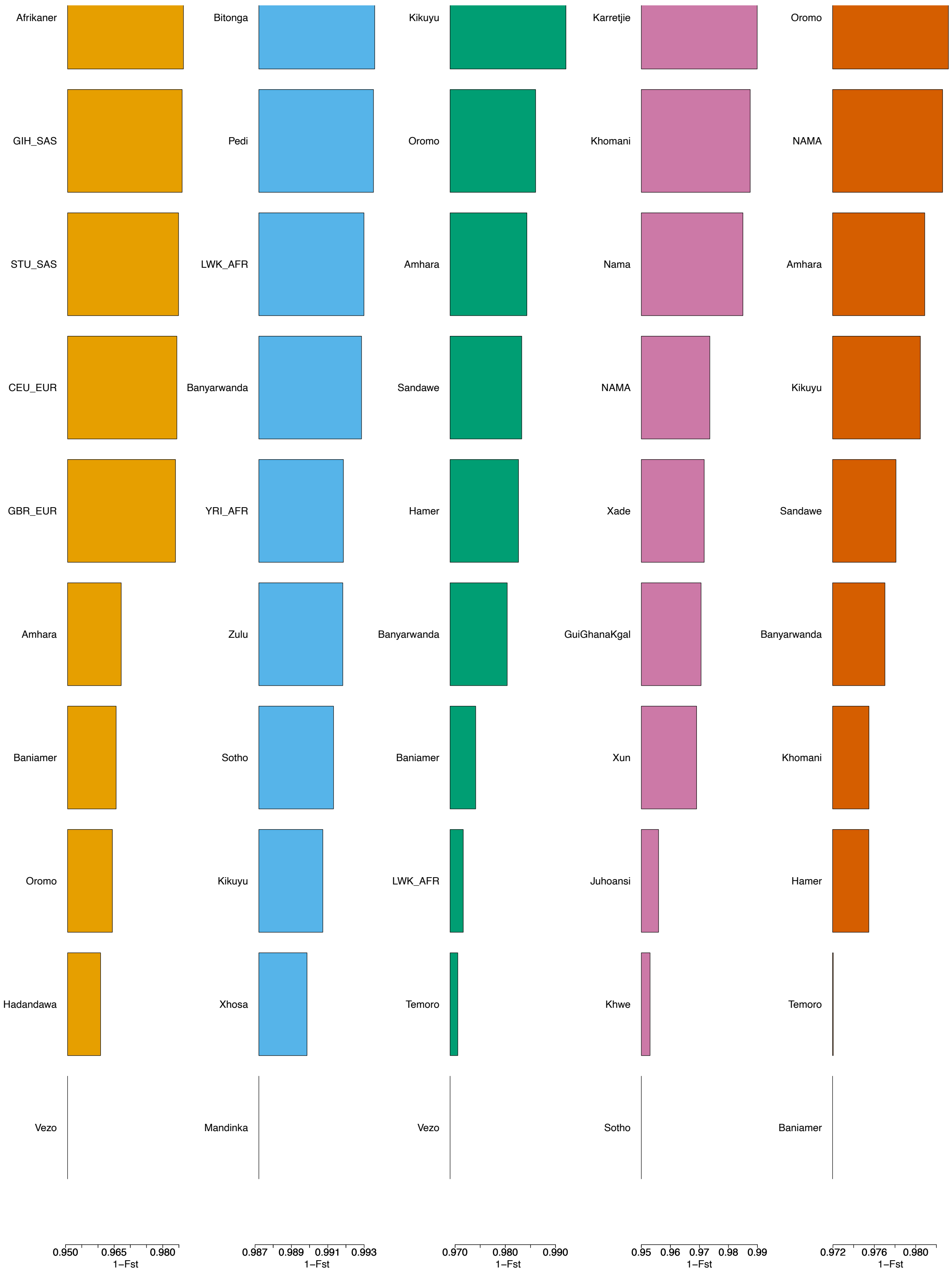


Supplementary Figure 56: 1-$F_{st}$ values from MOSAIC for the 5 constructed ancestries for Swellendam. The y-axis depicts the relative probability of jointly copying two chunks from the donor populations and x-axis is the genetic distance in centimorgans. The black lines represent the empirical coancestry curves across all target individuals, the light grey are per individual, and the green lines represent the fitted single-event coancestry curve. Above each plot, the number of the populations being examined is noted as a:b, with the number of generations since the admixture event in parentheses behind it. The number represents the order the ancestries appear in in the 1-$F_{st}$ plot.

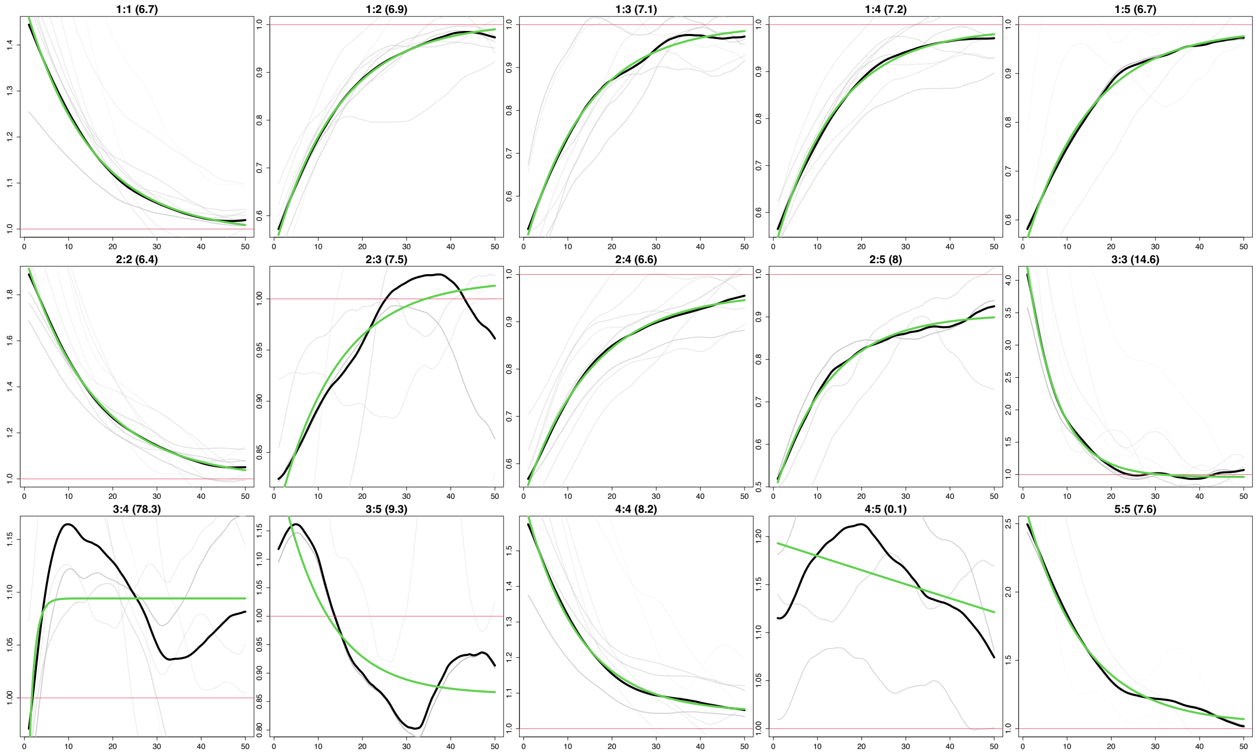


Supplementary Figure 57: Co-Ancestry curves from MOSAIC for the 5 constructed ancestries for Swellendam.
